# An aphid host-responsive RNA transcript that migrates systemically in plants promotes aphid reproduction

**DOI:** 10.1101/813964

**Authors:** Yazhou Chen, Archana Singh, Gemy G. Kaithakottill, Thomas C. Mathers, Matteo Gravino, Sam T. Mugford, Cock van Oosterhout, David Swarbreck, Saskia A. Hogenhout

## Abstract

Aphids are sap-feeding insects that colonize a broad range of plant species and often cause feeding damage and transmit plant pathogens, including bacteria, viruses and viroids. These insects feed from the plant vascular tissue, predominantly the phloem. However, it remains largely unknown how aphids, and other sap-feeding insects, establish intimate long-term interactions with plants. To identify aphid virulence factors, we took advantage of the ability of the green peach aphid *Myzus persicae* to colonize divergent plant species. We found that a *M. persicae* clone of near-identical females establishes stable colonies on nine plant species of five representative plant eudicot and monocot families that span the angiosperm phylogeny. Members of the novel aphid *Ya* family are differentially expressed in aphids on the nine plant species, are co-regulated and organized as tandem repeats in aphid genomes. Interestingly, aphids translocate *Ya* transcripts into plants and some transcripts migrate systemically within several plant species. RNAi-mediated knock down of *Ya* genes reduces *M. persicae* fecundity and *M. persicae* produces more progeny on transgenic plants that heterologously produce one of the systemically migrating *Ya* transcripts as a long non-coding (lnc)RNA. Taken together, our work led to the discovery of a new host-responsive aphid gene family that operate as virulence factors. Transcripts of this family translocate into plants, including a lncRNA that migrates systemically and promotes aphid reproduction.

**Significance Statement:** The green peach aphid *Myzus persicae* causes yield losses of a diverse range of economically important crops primarily as a vector of more than 100 different plant pathogens. We found that a single genotype of *M. persicae* is able to colonize nine plant species, including diverse dicots and maize, indicating that this aphid is truly polyphagous. Members of a new aphid *Ya* family undergoes coordinated expression changes in *M. persicae* depending on the plant species. The aphids translocate *Ya* transcripts into plants during feeding and these RNAs migrate to systemic leaves. Moreover, heterologous *in planta* expression of *M. persicae Ya1* as a long non-coding RNA promotes aphid reproduction. Our findings indicate that cross kingdom deployment of RNA is more common than thought.

## Introduction

Sap-feeding insects of the order Hemiptera include aphids, whiteflies, leafhoppers and planthoppers that have piercing-sucking mouthparts, named stylets, for feeding. Many species cause direct feeding damage, known for example as hopper burn (1, 2), though global economic yield losses caused by these insects are most often due to their abilities to transmit a diverse range of plant pathogens that include viruses, bacteria and plasmodium-like organisms, and also naked RNA molecules known as viroids (3, 4, 5-7, 8). The majority of insect herbivores are specialized to feed on one or a few closely related plant species (9). Nonetheless, some hemipteran insects are polyphagous. This includes the green peach aphid *Myzus persicae*, which is known to reproduce on over 400 plant species, and is also able to transmit divergent plant pathogens, including >100 plant viruses (10) and the potato spindle viroid (11). The factors involved in the ability of sap-feeding insects to establish intimate interactions with their plant hosts remain still largely unknown.

Sap-feeding insects of the order Hemiptera have stylets as mouthparts that are specialized to feed on plant sap, often from the phloem or xylem of the plant vascular tissue. The way by which sap-feeding insects move their stylets within plant tissues to reach the vascular tissues is arguably best investigated for aphids. These insects establish a long-term feeding site in the phloem sieve cells. However, before reaching the phloem, the stylets probe epidermis, mesophyll and other cells with each probe consisting of a short period of cell content ingestion, often referred to as ‘tasting’, followed by a short period of salivation (12). As soon as the stylets reach the phloem, aphids deposit saliva into the sieve cells followed by long periods of ingestion of phloem sap (13). The saliva introduced into cells is known as ‘watery’ saliva and is rich in proteins (14), some of which were shown to be effectors that modulate plant processes (15 - 21). The cycles of ‘tasting’ and salivation during the stylets path to the phloem are likely to help aphids perceive and adjust to their hosts.

We previously found that a parthenogenetically-reproducing *M. persicae* colony consisting of largely genetically identical females can adjust to the divergent plant species *Brassica rapa*, *Arabidopsis thaliana* and *Nicotiana benthamiana* via differential co-regulation of tandemly repeated gene families, including that of *Cathepsin B* (*CathB*), which are virulence factors that optimize the ability of *M. persicae* to colonize specific plant species (22). Interestingly, *CathB* genes are also differentially regulated in aphids on healthy versus those on virus-infected plants (23) and viruses are known to modulate plant defense responses (24, 25). These studies provide evidence that *M. persicae* has the ability to perceive the host plant status and adjust its gene expression accordingly.

Here, we build on our previous data, taking advantage of the ability of *M. persicae* to colonize divergent plant species to better understand how aphids colonize plants. We show that a single asexually-reproducing *M. persicae* clone forms stable colonies on nine plant species from five plant families. Via weighted gene co-expression network analysis (WGNCA) (26), we identified a novel gene family, named the *Ya* family, of which members adjust gene expression in a coordinated manner in response to the different plant species and that are organized as tandem repeats in the *M. persicae* genome. *Ya* transcripts translocate into plants during aphid feeding and migrate systemically, and the *M. persicae Ya1* transcript promotes aphid fecundity as a long non-coding (lnc) RNA when produced in plants. Our work shows evidence that the establishment of parasitic interactions between divergent organisms involves the translocation of a lncRNA virulence factor.

## Results

### *M. persicae* clone O establishes stable colonies on nine plant species that span the Angiosperm phylogeny

To establish *M. persicae* colonies on divergent plant hosts, we selected plant species across representative plant families across the Angiosperm phylogeny (27), including *B. rapa* (Br) and *A. thaliana* (At) of the Brassicaceae, *N. benthamiana* (Nb) and *Solanum tuberosum* (St) of Solanaceae, *Pisum sativum* (Ps) and *Phaseolus vulgaris* (Pv) of Fabaceae, *Helianthus annuus* (Ha) and *Chrysanthemum indicum* (Ci) of Asteraceae, and the monocot plant species, *Zea mays* (Zm) of the Poaceae (Fig. 1A). Individuals of asexually reproducing females of *M. persicae* clone O that were maintained on Br for at least three years were transferred to Br (control) and At, Nb, St, Ps, Pv, Ha, Ci, and Zm (Fig. 1A). *M. persicae* clone O achieved 100% survival rates and established stable colonies on these plant species (Fig. 1B; **Suppl. Fig. S1A-D**). Aphids survived equally well on Br and At from the start, and took about four weeks to achieve 100% survival rates on Nb, St, Ps, Ci and Zm, and 10 weeks on Pv and Ha.

**Fig. 1.**
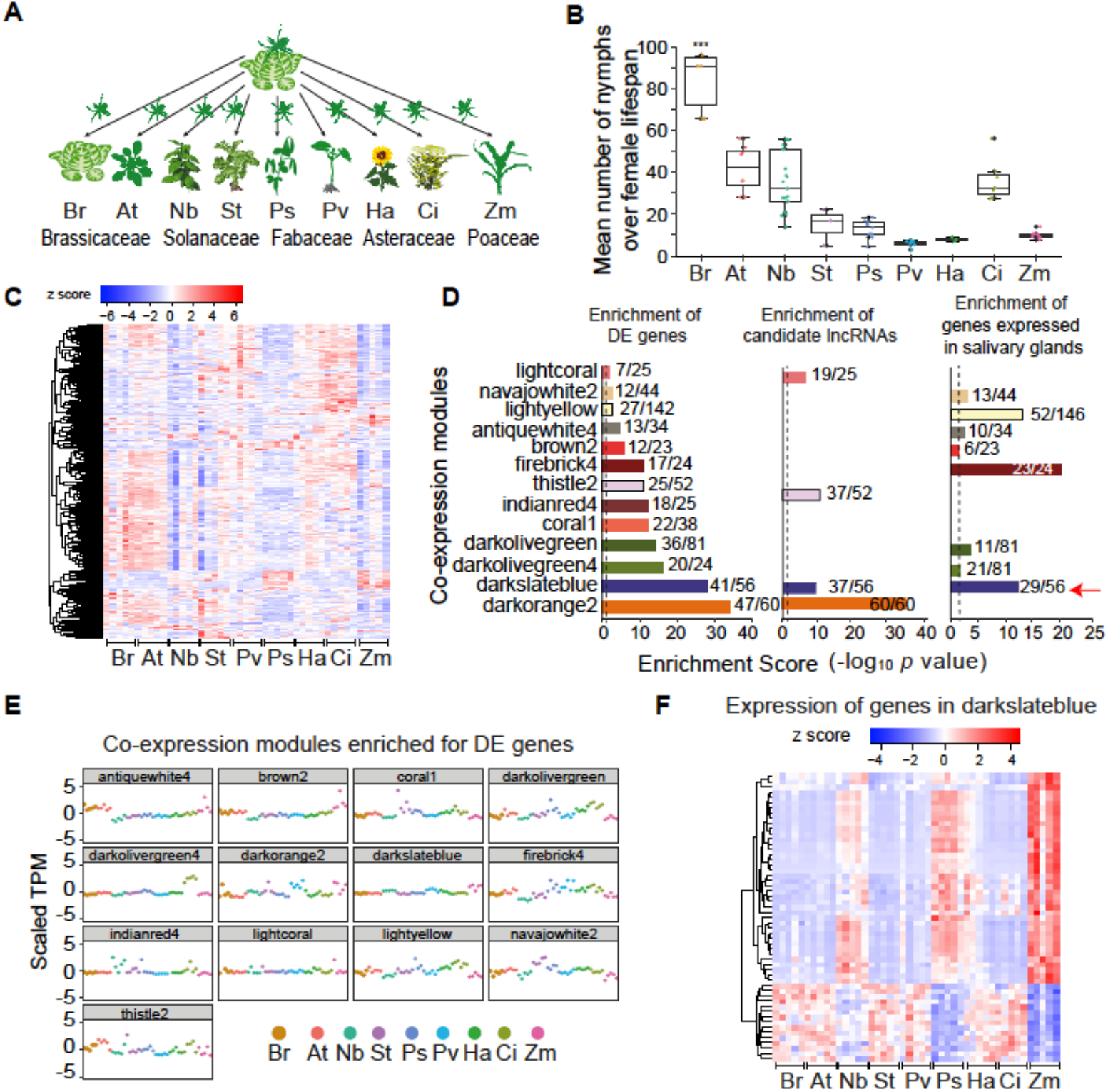
Colonisation of *M. persicae* on divergent host plant species involves co-expression of genes enriched in salivary gland expression and that encode lncRNAs. (**A**) Schematic overview of the experimental setup. *B. rapa* (Br), *A. thaliana* (At), *N. benthamiana* (Nb), *S. tuberosum* (St), *C. indicum* (Ci), *H. annuus* (Ha), *P. sativum* (Ps), *P. vulgaris* (Pv) and *Zea mays* (Zm). (**B**) Reproduction levels of stable *M. persicae* colonies on each of the nine plant species as shown in A. Box plots show mean numbers of nymphs produced over the lifespan of an adult female on each plant species. (**C**) Heatmaps of log transformed TPM values of 1 984 DE genes of *M*. *persicae* colonies on Br versus colonies on one or more of the other 8 plant species as shown in A at 5 biological replicates per host. (**D**) Enrichment scores of 13 WGNCA modules. The x/y above the bars indicating gene numbers in the enriched category (x) and module (y). Dashed vertical lines, enrichment score 2 (*p* value = 0.01). Red arrow, darkslateblue module. (**E**) Scatter plots of 13 WGCNA modules enriched for DE genes. (**F**) Heatmap of log transformed TPM values of genes in the darkslateblue module.

### Genes differentially expressed in *M. persicae* on nine plant species are enriched for salivary gland and candidate long non-coding RNAs

Next, we generated transcriptomes of the stable *M. persicae* colonies on the nine plant species. RNA-seq data of five independent biological replicates per plant species were combined with previous RNA-seq data (LIB1777, 22) to give a total of 1.38 billion RNA-seq reads that were assembled into transcripts using a genome-guided approach (**Suppl. Table S1; GSE129667**). We identified in total 45 972 transcripts (corresponding to 19 556 genes) with 30 127 (65.5%; 18 529 genes) annotated previously (22) and 15 845 (34.5%) novel transcripts (**Suppl. Fig. S2A**). Of the 45 972 transcripts, 6 581 transcripts (3 025 genes) had low protein-coding potentials and were assigned candidate lncRNAs (**Suppl. Fig. S2B**).

Relatively to the *M. persicae* colonies on Br (original host), 2 490 (1 984 genes) of the 45 972 transcripts (5.41%) were significantly differentially expressed (DE) in the aphids on at least one or more of the other 8 host plant species (fold change ≥ 2, *p* value ≤ 0.05, FDR ≤ 5%) (**Suppl. Table S2**) and the majority of these show host-specific expression patterns (Fig. 1C). The DE transcripts were enriched in functions of oxidation-reduction processes, proteolysis (including *CathB*), and sensory perception of taste (**Suppl. Fig. S3**). Based on a mean gene expression value of TPM > 5 across five biological replicates per hosts, 11 824 genes (out of 19 556 total) were selected for WGCNA, which identified 77 co-expression modules comprising 7 864 genes (**Suppl. Fig. S4**; **Suppl. Table S3**). Of the 1 984 DE genes, 1 364 (68%) were included among the genes of the 77 co-expression modules, and 13 modules were enriched for DE genes (313 DE genes of the total DE genes (16%); Fisher exact test, *p* value < 0.05) (Fig. 1D; **Suppl. Table S3**). Heatmaps based on normalized TPM values of these 13 modules showed different expression patterns in aphids depending on the plant host species (Fig. 1D; **Suppl. Fig. S5**), suggesting that *M. persicae* coordinates the expression of groups of genes in response to the plant species.

Remarkably, the darkslateblue, darkorange2, lightcoral, and thistle2 modules among the 13 enriched for DE genes were also enriched for candidate lncRNAs (Fig. 1E; **Suppl. Table S3**). As well, 8 out of 13 modules, including the darkslateblue, were enriched for *M. persicae* genes expressed in the salivary glands (28) (Fig. 1E; **Suppl. Table S3**) and 3 out of 13 modules were enriched for gut-expressed genes (**Suppl. Fig. S6**; **Suppl. Table S3**). We also noticed that 12 out of 13 modules included genes that lie adjacently to each other as tandem repeats within the aphid genome (**Suppl. Table S3**). The number of gene repeats varied from 2 to 8, with the latter being in the darkslateblue module. Hence, different attributes were enriched in the darkslateblue module, including expression in the salivary glands, candidate lncRNAs and tandemly repeated genes. The darkslateblue module is the only one among the 77 modules that contained two groups of genes with exact opposite host-responsive expression patterns (Fig. 1F).

### *M. persicae* preferentially translocates RNA transcripts of DE and candidate lncRNA genes into plants

Given the enrichment of differentially expressed salivary gland genes that encode candidate lncRNAs, we investigated if these transcripts translocate into plants when aphids feed. Leaves of 4-week-old *A. thaliana* plants were caged with 20 adult aphids (feeding site) or no aphids (control) for 24 hours (Fig. 2A; **Suppl. Table S4**). Reads that uniquely aligned to the *M. persicae* genome were identified, and at TPM ≥ 50, these corresponded to between 1 837 to 3 154 aphid transcripts, depending on the biological replicate, in leaf sections exposed to aphids only (**Suppl. Table S5**). Based on the presence in at least three biological replicates, 3 186 *M. persicae* transcripts of which 201 are candidate lncRNAs were found present in feeding sites (Fig. 2A). The candidate lncRNAs were enriched among the transcripts found in the feeding site (Fisher’s Exact Test, *p* = 1.02E-48) (Fig. 2A; **Suppl. Table S4**). The candidate lncRNAs in the feeding site belonged to ten co-expression modules that were enriched for DE genes, including the darkslateblue module (Fig. 2B). Moreover, the *M. persicae* transcripts in the feeding site were enriched for DE genes and salivary gland transcripts (Fisher’s Exact Test, *p* = 3.7E −13 and *p* = 0.03, respectively) and in both categories the transcripts were also enriched for candidate lncRNAs (Fisher’s Exact Test, *p* = 0.002 and *p* = 3.8E-35, respectively) (Fig. 2C). Therefore, *M. persicae* may preferentially translocate candidate lncRNAs of salivary gland expressed genes that are DE in aphids on divergent hosts into feeding sites.

**Fig. 2.**
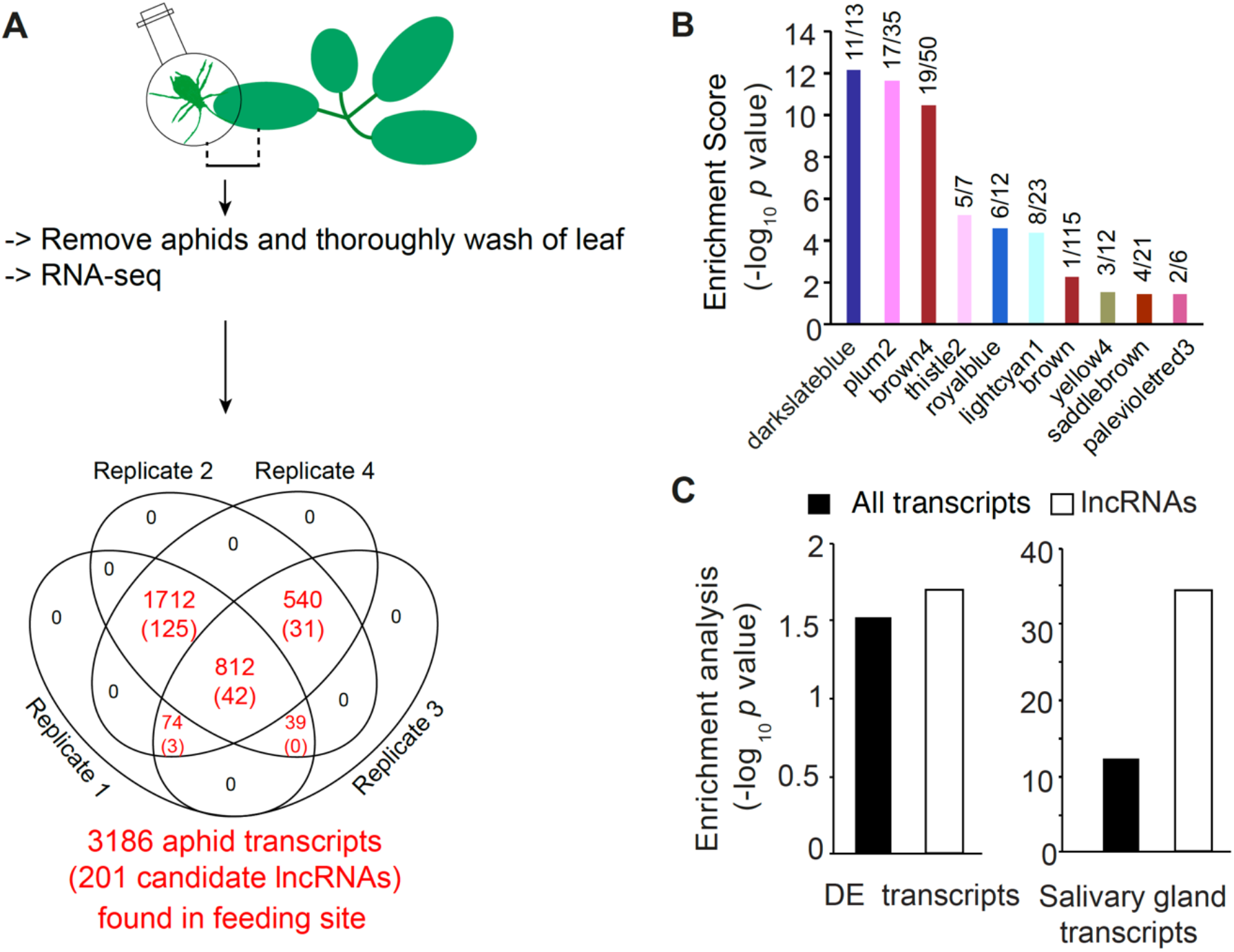
Aphid transcripts at the aphid feeding site are enriched for differential expression in aphids on nine plant species, candidate lncRNA and expression in aphid salivary glands. (**A**) Schematic overview of the experimental setup. Venn diagram shows the number of aphid transcripts found in plants at the aphid feeding site at TPM ≥ 50 in one of the replicates and/or presence of transcripts in at least three biological replicates. (**B**) Transcripts in the feeding site were enriched for candidate lncRNAs in 10 out of the 13 co-expression modules enriched for DE genes (see Fig. 1D); x/y above the bars indicating lncRNAs (x) and the total number of transcripts that belong to the module found in the feeding site (y). (**C**) Enrichment of genes that are DE (left graph) and expressed in salivary glands (28) (right graph) of aphid transcripts found in the plant at the aphid feeding. Black bars, all aphid transcripts. White bars, candidate lncRNAs.

### The co-regulated *M. persicae* genes of the darkslateblue module includes all 30 members of the *Ya* family tandem repeat

Upon further analyses of genes in the darkslateblue module, 37 out of total of 56 genes were found to encode candidate lncRNAs. Of these 37, 23 were identified to have high sequence similarities (± 80%) and located in close proximity to one another on six scaffolds. These genes were organized in series of tandem repeats with less than 1-kb distance between the gene copies. Given these characteristics, the genes appear to belong to a gene family.

Tandemly duplicated genes with highly similar sequences are often misannotated due to incorrect alignment of RNA-seq reads. Therefore, we manually this gene family by searching via BLAST the entire *M. persicae* genome for regions that align to a 148-bp nucleotide sequence that was found to be conserved among the 23 candidate lncRNA genes (**Suppl. Fig. S7**). This resulted in the identification of 7 additional genes to make a 30-member gene family (**Suppl. Table S6**). We updated the transcriptome assembly for this gene locus with the manually annotation (GSE129667). We named this gene family the *Ya* family (Yá means aphid in Chinese).

The 3’ ends of the 30 *Ya* transcripts were manually corrected based on the presence of a poly(A) signal and 5’ ends via identification of conserved sequences among the *Ya* transcripts. All *Ya* genes have a three-exon structure, and they show a modest to high sequence conservation (ranging between 84.6 – 99.1 % nucleotide identities compared to *Ya1*), including a region that corresponds to a small open reading frame (ORF) that may translate into a peptide of 38 amino acids in all 30 *Ya* transcripts (Fig. 3A; **Suppl. Fig. S8**).

**Fig. 3.**
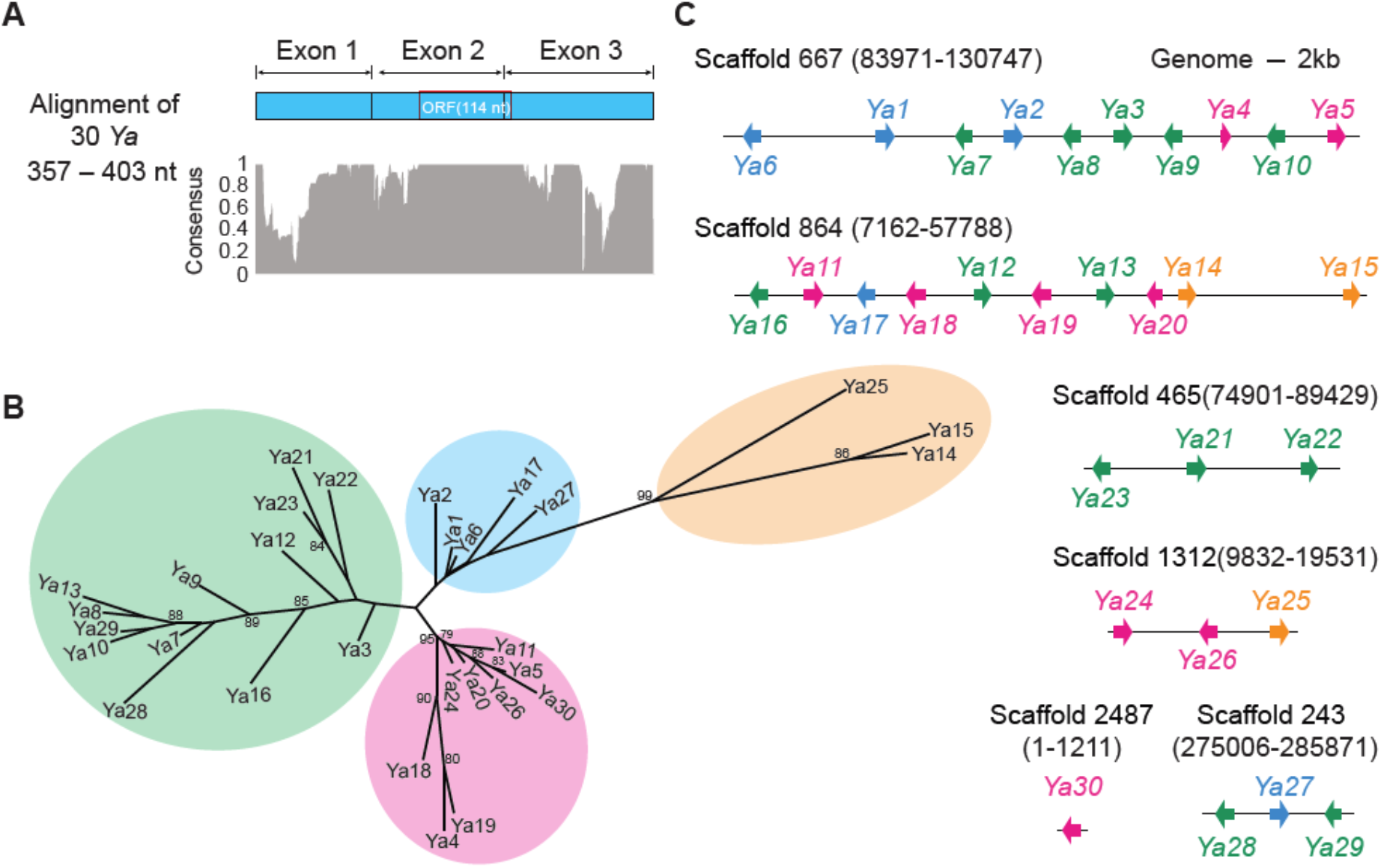
Annotation of the *Ya* gene family in the *M. persicae* genome. (**A**) Level of conservation (consensus) among aligned nucleotide sequences of mature transcripts of 30 *Ya* genes (See **Suppl. Fig. S8** for alignment). The small open reading frame (ORF) may translate into a 38 amino acid (aa) peptide. (**B**) Phylogeny based on the alignment of 30 *Ya* transcripts shown in A. (**C**) Locations of *Ya* gene repeats on *M. persicae* genomic scaffolds. *Ya* genes are shown as box arrows of which the colours match the clades of the Ya phylogenetic tree shown in B.

Phylogenetic analyses based on nucleotide sequence alignment (**Suppl. Fig. S8**) group the *Ya* genes into roughly 4 distinct clades (Fig. 3B). As well, we confirmed that the *Ya* genes form several tandem repeats in the *M. persicae* genome in which the *Ya* genes were often, but not always, organized in pairs facing outwards on opposite genomic strands (Fig. 3C). The pairs belong to the same or different phylogenetic clusters of the *Ya* phylogenetic tree. Repeating the *M. persicae* nine-host swap WGCNA analysis with the new set of manually annotated 30 *M. persicae Ya* genes grouped all *Ya* members within the same (darkslateblue) co-expression module (**Suppl. Fig. S9**), as before, suggesting that the 30 *Ya* genes are co-regulated.

*Ya* genes were also found in five other aphid species for which whole-genome sequences are available. To further compare the number of *Ya* genes among aphid species, the 148-bp *Ya1* exon 2 sequence was BLAST searched against genome assemblies of *Aphis glycines*, *Acyrthosiphon pisum, Diuraphis noxia*, *M. cerasi* and *Rhopalosiphum padi* and subsequently against public transcriptome datasets for these species. Transcripts that matched the genomic sequences were assigned as *Ya* genes, resulting in the identification of 13 candidate *Ya* genes in *Aphis glycines*, 17 in *Diuraphis noxia*, 29 in *M. cerasi* and 33 in *Rhopalosiphum padi*. In all five aphid species, *Ya* genes are tandem repeats in genomes. We did not find *Ya* genes in genomes of hemipteran insect species beyond aphids.

Therefore, *Ya* genes are unique to aphids and are part of larger families that form tandem repeats in aphid genomes and are differentially expressed and co-regulated in *M. persicae* on nine divergent host plant species.

### Translocated *M. persicae Ya* transcripts migrate systemically in plants

To asses if aphid *Ya* transcripts migrate systemically within plants, we caged twenty adult aphids on leaves of 4-week-old *A. thaliana* plants for 24 hrs, and then examined the presence of aphid transcripts in the caged area (feeding site), the area from the petiole to the aphid cage on the leaf (near feeding site) and a systemic leaf (Fig. 4A). We designed specific primers for the nine *Ya* transcripts found in the feeding site (**Suppl. Fig. S10, Suppl. Table S7**). For the six *Ya* transcripts *Ya1*, *Ya2*, *Ya3*, *Ya6*, *Ya11*, *Ya17* the correct sizes of amplification products were obtained and the sequences of these amplification products matched those of the *Ya* transcripts, whereas the presence of *Ya6*, *Ya21* and *Y23* transcripts was not confirmed (**Suppl. Fig. S10**). *Ya1*, *Ya2*, and *Ya17* transcripts were detected in feeding and systemic sites indicating that these transcripts migrated away from the feeding site into systemic tissues (**Suppl. Fig. S10**). We focused further analyses on the *Ya1* transcript, because the heatmap of the darkslateblue module suggested that *Ya1* is upregulated in aphids on Br and At (**Suppl. Fig. S9**). The predicted size of the *Ya1* transcript is 382 nt (Fig. 4B, **Suppl. Fig. S11A**). The presence of *Ya* transcripts in aphids and plants were analysed by reverse transcriptase PCRs with a series of specific primers and by northern blot hybridizations with a *Ya1* probe. PCRs showed that a 357-nt *Ya* transcript that starts at nucleotide 25 of exon 1 was detected in the aphid, and not in plants, whereas a shorter *Ya1* transcript of 273 nt that starts at nucleotide 110 near the start of the sequence corresponding to exon 2 was detected in the plant and migrated systemically (Fig. 4B, **Suppl. Fig. S11*A* and *B***). 3’ RACE experiments showed that the *Ya1* transcript has a poly(A) tail (**Suppl. Fig. S11C**). Northern blots confirmed the sizes of the *Ya* transcripts in aphids and plants; the fragments that hybridized to the *Ya1-SP6* in aphids were larger than the 291 nt *Ya1-SP6* transcript, whereas the Ya1 fragments in plants were shorter than this transcript (Fig. 4B). These data indicate that the first ±100 nt at the 5’ end of the 357-nt aphid *Ya1* transcript is processed to produce a 273-nt *Ya1* transcript upon translocation into plants.

**Fig. 4.**
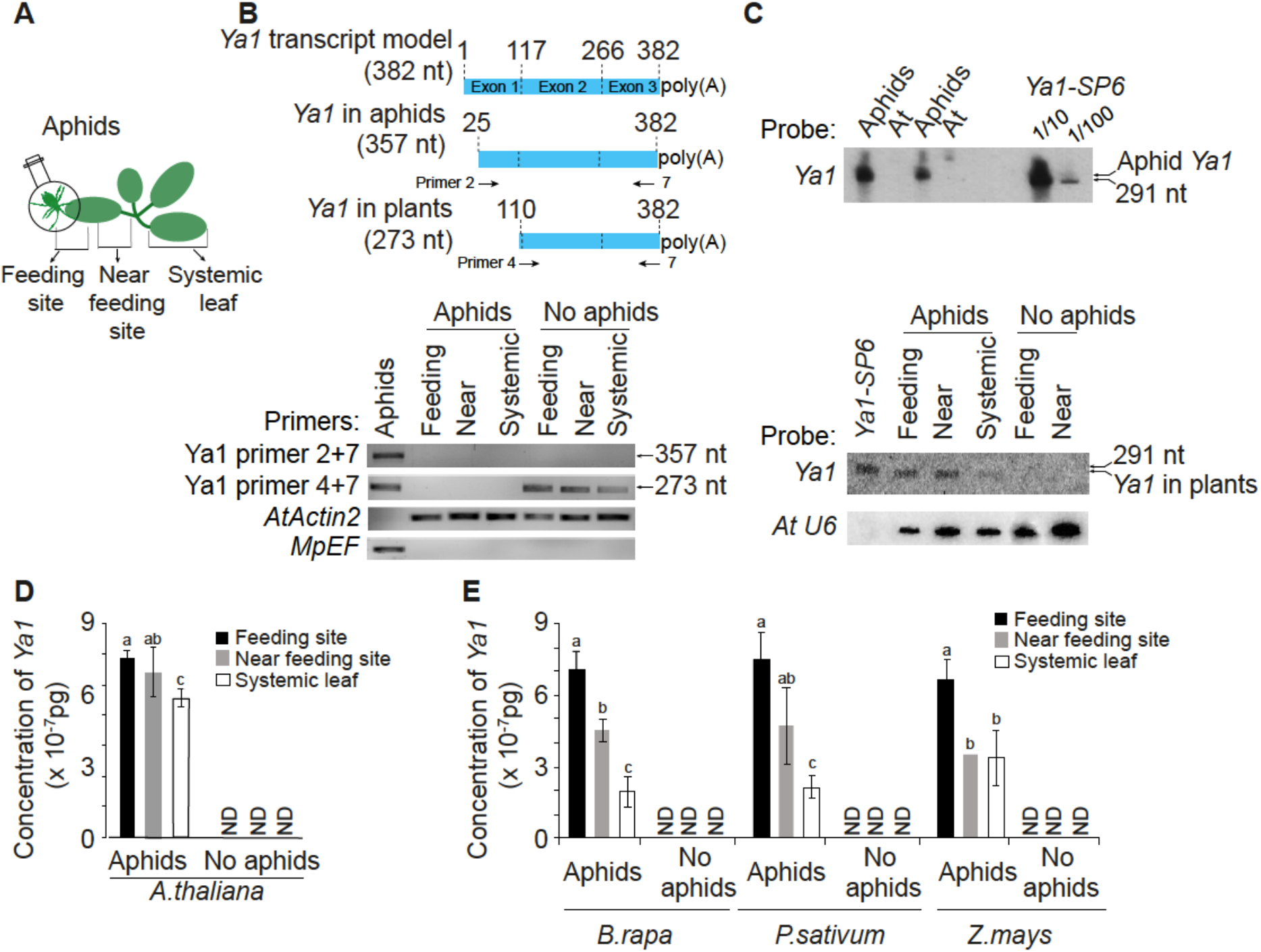
The *M. persicae Ya1* transcript translocates and systemically migrates in plants. (**A**) Schematic overview of the experimental setup. (**B**) RT-PCR of Ya1 transcripts in aphids and aphid-exposed plants. The 357-nt Ya1 transcript was found in aphids, and not in plants. A 273-nt transcript was found at aphid feeding sites and to migrate systemically in plants. *A. thaliana* actin 2 (AtActin2) and *M. persicae* elongation factor (MpEF) were used to control for presence of RNA. (**C**) Northern blot hybridizations with a Ya1 probe to detect Ya transcripts in aphids and plants. Arrows at right of the blots indicate the locations of the in vitro synthesized 291-nt *Ya1-SP6* RNA transcript and *Ya* transcripts found in aphids and plants. Blots were stripped and labelled with the *A. thaliana U6* probe to assess RNA loading levels among the samples. (**D, E**) Quantitative RT-PCR to detect systemic migration of *M. persicae* in (D) *A. thaliana* and (E) *B. rapa*, *P. sativum*, and *Z. mays*. Y-axes, *Ya1* concentrations based on a standard curve (**Suppl. Fig. S13**). ND, not detected. Bars indicate mean ± standard deviation (SD) of concentrations of *Ya1* and two independent biological replicates. Different letters above the bars indicate significant differences between groups (*p* value < 0.01, Student *t* test).

Northern blots and qRT-PCRs showed that the *Ya1* transcript gradually decreases in concentration from feeding sites to near feeding sites and systemic leaves of *A. thaliana* (Fig. 4C, 4D, **Suppl. Fig. S13**). *Ya1* also migrated systemically in *B. rapa*, *P. sativum* and *Z. mays* exposed to *M. persicae* (Fig. 4E, **Suppl. Fig. S12**). Sequencing of the PCR products derived from of the feeding sites, near feeding sites and systemic leaves of these three hosts revealed identical sequences to *Ya1*, which is different from the other *Ya* family members (**Suppl. Fig. S14**). Therefore, *M. persicae* deposits *Ya1* transcripts into four divergent host plant species during feeding and these transcripts migrate systemically within these plants away from aphid feeding sites.

### *M. persicae Ya1* is a virulence factor

Differential expression of *Ya1* in *M. persicae* among hosts and migration of *Ya1* in various of hosts suggest *Ya1* may play roles in aphid-host interactions. To investigate this, we generated transgenic lines expressing dsRNA corresponding to the *Ya1* sequence for plant-mediated RNAi of *Ya1* in *M. persicae*. Two independent *A. thaliana* lines 1-5 and 2-8 successfully knocked down *Ya1* expression compared to WT Col-0 and lines producing dsRNA corresponding to GFP (dsGFP) in *M. persicae* (Fig. 5A). *M. persicae* on *A. thaliana* lines 1-5 and 2-8 had reduced fecundity compared to WT Col-0 and dsGFP lines (Fig. 5B). Therefore, knock down of the expression of *Ya1* reduces *M. persicae* reproduction on *A. thaliana*.

**Fig. 5.**
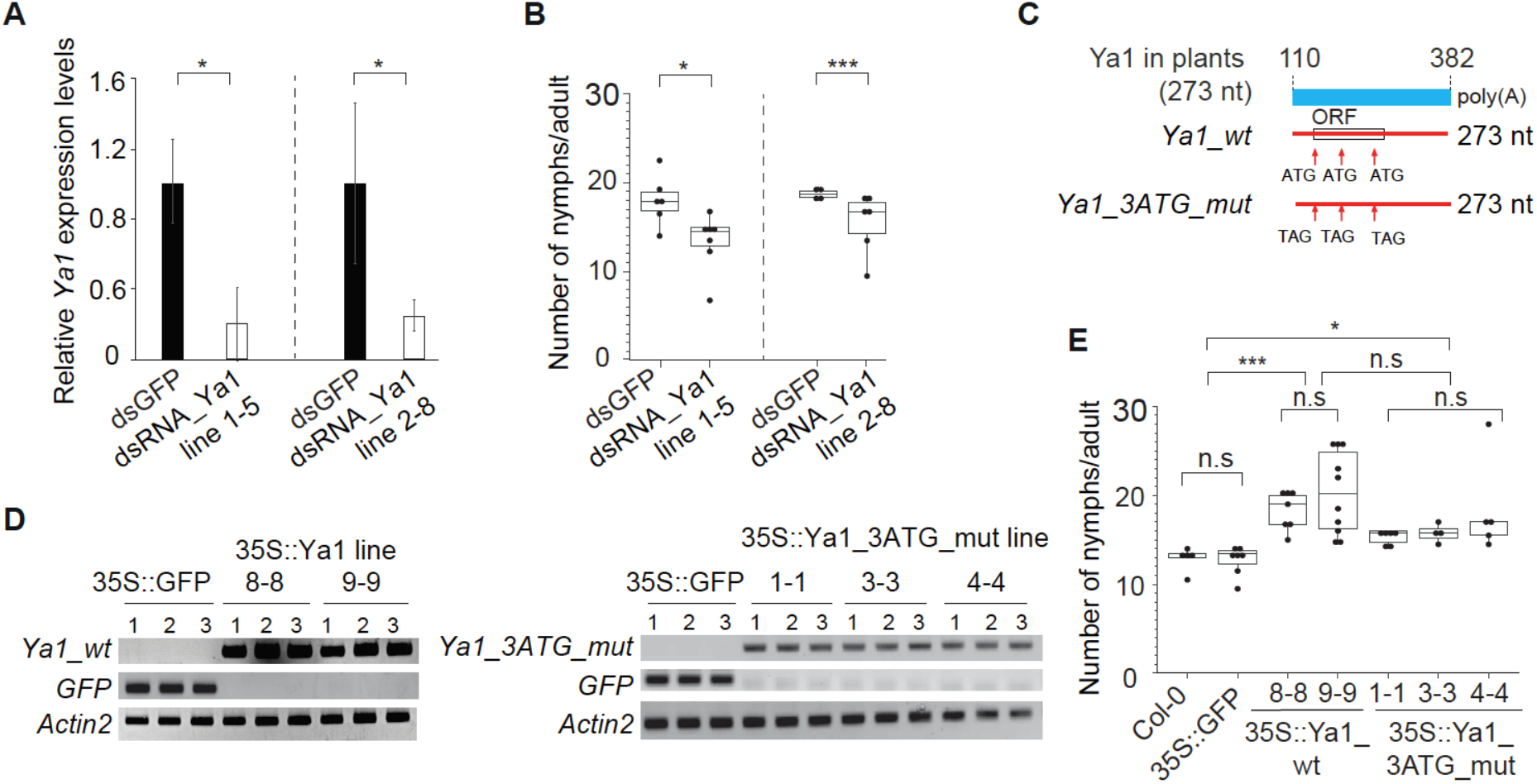
*M. persicae Ya* lncRNA promotes *M. persicae* colonization on *A. thaliana*. (A) Quantitative RT-PCRs showing knock-down of *Ya1* in *M. persicae* reared on plants that stably produce dsRNA_Ya1 relatively to those that produce dsRNA_GFP (controls). Y-axis indicates the *Ya1* expression levels relative to *M. persicae* EF reference gene. Bars represent relative mean ± standard deviation (SD) expression values at n = 4-8 in which n is one aphid per plant. \**p* < 0.01, Student *t* test. (B) Reduction in fecundity of Ya1-RNAi aphids tested for *Ya1* knock down in A. Each data point (black dot) represents the number of nymphs produced by one adult female per plant. Box plots show distribution of data points collected from n = 4-8 female aphids per *A. thaliana* line as shown on the x-axis. (C) Mutation of the three ATGs within the ORF of 38-aa of the Ya1 transcript into TAG stop codons. (D) Detection of transgene transcripts by RT-PCRs at three plants per lines (1-3). Detection of the *A. thaliana* Actin2 transcript was used as control for presence of RNA. (E) Stable expression of *Ya1_wt* and *Ya1_3ATG* promotes *M. persicae* reproduction on plants. Box plots show distribution of nymphs produced by an adult female aphid per plant (black dots) collected from n = 4-10 female aphids per *A. thaliana* line. \*\**p* < 0.05, \*\*\**p* < 0.001, ANOVA followed by a Tukey–Kramer post-hoc test.

Given that aphids deposit *Ya1* into plants and that this RNA migrates systemically, we investigated if stable expression of *Ya1 in planta* affects *M. persicae* performance. We generated stable transgenic plants that produced the 273-nt (exons 2 + 3) Ya1 transcript (35S::Ya1 (Col-0) lines 8-8 and 9-9) and 273-nt *Ya1* mutants in which three ATG start sites within the 38-aa open reading frame were mutated to stop codons (35S::Ya1_mut (Col-0) lines 1-1, 3-3 and 4-4) (Fig. 5C and D, **Suppl. Fig. S15**). *M. persicae* produced more progeny on both 35S::Ya1 and 35S::Ya1_mut compared to 35S:GFP and Col-0 plants (Fig. 5E, **Suppl. Fig. S16**), indicating that the *Ya1* RNA transcript modulates plant processes that leads to an increase of *M. persicae* fecundity. Taken together, our data show that *M. persicae* translocate the *Ya* transcript into plants during feeding and this transcript migrates systemically within plants and promotes fecundity of this aphid. Therefore, *Ya1* is an aphid lncRNA virulence factor.

## Discussion

We found that progeny derived from a single asexually-reproducing *M. persicae* clone O female stably colonizes nine divergent plant species of five families that span the Angiosperm phylogeny (27), demonstrating that *M. persicae* is truly polyphagous. We identified *M. persicae* genes that show coordinated up- and down-regulation depending on the plant species the aphid is colonizing and that are organized as repeats in the aphid genome. The genes are organized in co-expression modules of which 13 are enriched for DE genes, genes expressed in the salivary glands and candidate lncRNAs. One of these modules includes all members of the *Ya* family with 30 genes that are tandemly repeated in the *M. persicae* genome. Moreover, transcripts from six *Ya* genes translocate into plants during aphid feeding and three migrate systemically away from the aphid feeding site to distant leaves. Via RNAi-mediated knock down of *Ya* expression we show evidence that *Ya* genes contribute to aphid performance on *A. thaliana* and are therefore virulence factors. Heterologous expression of the *Ya1* transcript without the small putative ORF *in planta* promotes aphid performance indicating that the *Ya1* RNA transcript acts as a lncRNA virulence factor.

Because genome annotation pipelines are generally focused on protein-coding genes, the majority *Ya* family members were not annotated in the first version of the *M. persicae* genome annotation (22). Moreover, due to the *Ya* family being organised as tandem repeats of highly conserved sequences, genome-guided transcript assembly generated misassembled transcripts. To overcome this we did thorough manual annotation of the *Ya* region and were able to identify the 30-member *Ya* family, characterize the expression patterns of each of the family members in *M. persicae* in response to the nine plant species, and to assess which family members translocate to plants during aphid feeding. We show that *M. persicae Ya* members are characterized by a 3-exon gene model and produce transcripts ranging from 357 – 403 nucleotides in lengths. When analysing other hemipterans, we identified tandemly repeated *Ya* family members only in the genomes of only other aphid species among hemipterans for which genome sequence data are available, suggesting that these lncRNAs are unique to aphids.

Despite *Ya* genes being annotated as genes with low-coding potential, *Ya* transcripts have a conserved small ORF that potentially translates into a 38 amino acid peptide. Mutations that prevent translation of this peptide within the *Ya1* transcript in plants does not affect the ability of *Ya1* to promote aphid fecundity, indicating that the *Ya1* has virulence activity as a lncRNA. ORFs have been detected in number of transcripts known to function as lncRNAs, including RNAs associated with ribosome function (30; 31) and X-inactive specific transcript (*Xist*) that regulates the X chromosome inactivation process (32), and it is being debated whether lncRNA ORFs may be translated in some situations (33). Whether the small ORF found to be conserved across *Ya genes* has a function remains to be determined, but based on our annotations and functional analyses of *Ya1*, at least one member of the *Ya* gene family appears to function as a lncRNA.

Several parasites translocate small RNAs into their hosts (34 - 36). The functions of these transkingdom small RNAs are known for only a few parasites. For example, small RNAs of the fungal plant pathogen *Botrytis cinerea* interact with the AGO1 protein of the *A. thaliana* RNA interference (RNAi) machinery to suppress plant defense genes (37) and microRNAs of the parasitic plant dodder target host messenger RNAs involved in plant pathogenesis (38). In the opposite direction, plants export specific microRNAs to control virulence of a pathogenic fungus (39). In some parasite-host interactions, a large number of long transcripts (>200 nt) were found in hosts, including RNA transcripts of *Cryptosporidium parvum* in the nuclei of human intestinal epithelial cells (40) and transcripts of dodder that systemically migrate in *Solanum lycopersicum* and *A. thaliana* (41). However, it is unclear if these larger parasite RNAs modulate parasite-host interactions. Here we show that *M. persicae* RNA transcripts translocate into plants. These include transcripts of the *Ya* family that migrate systemically. Knock down of *Ya* gene expression via RNAi reduces aphid fecundity, whereas *in planta* expression of *Ya1* as a lncRNA promotes aphid fecundity. This suggests that the aphid *Ya* genes are virulence factors and that the *Ya1* transcript controls aphid performance via the plant.

The *M. persicae Ya1* lncRNA may be an effector that modulates specific plant processes. Interestingly, the entire *M. persicae Ya* family of 30 members is part of the darkslateblue module, which is the only module among the 77 co-expression modules that consists of two groups of genes with exact opposite expression patterns. The module also includes several protein-coding genes with similarities to haemolymph juvenile hormone binding and WD40 and EGF-like domain-containing proteins that have roles in signal transduction in insects. Given this finding, a non-mutually exclusive possibility is that *Ya1* and other *Ya* lncRNAs may have a sensing role within aphids; for example, lncRNAs not degraded in plants may migrate back into the aphid to regulate aphid gene expression, in agreement with lncRNAs often have functions in gene expression regulation (42 - 44). Whereas the mechanism by which *Ya1* controls aphid fecundity will have to be determined, results described herein indicate that members of the *Ya* gene family are aphid-specific virulence factors.

## Material and Methods

See Supplementary File 1.

## Acknowledgments

We thank Danielle Goff-Legett, Alexandra Kolodyazhnaya and Christian Aarssen for assisting with various experiments in this study, and Andrew David Lyle Nelson and Upendra Kumar Devisetty at The University of Arizona for their assistance with the initial steps of lncRNA identifications. We are grateful to Anna Jordan, Darrell Bean, Susannah Gill, and Ian Bedford of the JIC Entomology Facility for rearing and taking care of aphid colonies and Horticultural Services for growing the plants used in this study. The project was funded by the Biotechnology and Biological Sciences Research Council (BBSRC) Industrial Partnership Award with Syngenta Ltd grant BB/R009481/1 and the BBSRC Institute Strategic Program Grant (ISPG) Plant Health BB/P012574/1.

