## Supplementary File 1 for "An aphid host-responsive RNA transcript that migrates systemically in plants promotes aphid reproduction"

**Supplementary file S1**

### Materials and Methods

#### Plants and growth conditions

Seeds of *B. rapa*, *A. thaliana*, *N. benthamiana* were provided by John Innes Centre horticultural department. *Solanum tuberosum* (Desiree) were purchased from Marks & Spencer (Norwich, UK). Seeds of *C. indicum* (polar star), *H. annuus* (Helios X Helios Flame), *P. vulgaris* (NYFB) were purchased from Thompson Morgan (Ipswich, UK). Seeds of *P. sativum* (JI3253) were provided by Claire Domoney (John Innes Centre, UK). Seeds of *Z. mays* (Early sunglow corn) were provided by Ian Bedford (John Innes Centre, UK). Seeds were directly sown into soil (peat-based compost). Plants were grown in a controlled environmental room at a constant temperature of 22°C and 70% humidity under a 10 h day /14 h night cycle.

#### *M. persicae* transfer to 9 plant species

A colony of *M. persicae* clone O that started from a single female was established on *B. rapa* in a growth chamber (14 h light, 10 h dark at constant 20 °C, 75% humidity) in 2010. From this founder colony on *B. rapa*, approximately 500 asexual individuals were transferred to each of 9 plant species, including *B. rapa* (used as reference) and *A. thaliana*, as a representative of another plant species of the family Brassicaceae, and plant species belonging to 4 additional plant families, including *N. benthamiana* and *S. tuberosum* (Solanaceae), *C. indicum* and *H. annuus* (Asteraceae), *P. sativum* and *P. vulgaris* (*C. indicum* and *H. annuus*), and the monocot maize (*Z. mays*) (Poaceae). The aphids were maintained for about 4 generations on these plants, except for *C. indicum* and *P. vulgaris*, on which the aphids were reared for 10 generations. Aphid colonies on all 9 plant species were maintained in the same growth chamber (14 h light, 10 h dark at constant 20 °C, 75% humidity).

Stable aphid populations on the 9 hosts were analyzed for development time in days between births and emergence of adults, longevities in days between births to deaths, weights in mg per adult and reproduction rates measured by numbers of nymphs divided by the number of adults on days 7, 9 and 11 for colonies on *B. rapa*, *A. thaliana*, *N. benthamiana* and *S. tuberosum* and on days 7, 9, 11, 13, 15, 17 and 18 for colonies on *C. indicum*, *H. annuus*, *P. sativum*, *P. vulgaris* and *Z. mays*.

**Transcriptome sequencing of *M. persicae* on 9 hosts**

Approximately 100 aphids were harvested from each plant species, at 5 independent biological replicates per plant species, and snap frozen in liquid nitrogen. Aphid samples were ground to powder using 5 mm stainless steel beads (Qiagen, Germany) and a TissueLyser II (Qiagen, Germany), and RNA was extracted using Trizol reagent (Sigma, UK) and the Qiagen RNeasy MinElute Cleanup Kit (Qiagen, Germany), which included and on-column DNase digestion.

**RNA sequencing and transcriptome assembly**

Strand-specific libraries were constructed from mRNAs isolated from 1 ug of total aphid RNA using the poly-A method of the Illumina TruSeq RNA Library Preparation kit (Illumina, US) following the manufacturer's procedures. cDNA was synthesized by 10 cycles of PCR to amplify the fragments. Libraries were then pooled and sequenced on a HiSeq 2000 generating 150-bp paired-end sequences (Earlham Institute, Norwich, UK). A genome-guided transcriptome assembly was generated with RNA-seq data of the 45 libraries of the nine host experiments, five replicates each (data generated herein as described above) and RNA-seq data generated from library LIB1777 (1), as follows. Reads were trimmed for low quality and adapter using Trim Galore! v0.4.0 with default settings (2). Trimmed reads were aligned to *M. persicae* reference genome G006 v1 (1) by HISat2 version-2.0.5 (3). The RNA-seq reads of all 46 libraries were merged together in one BAM file (Binary version of Sequence Alignment/Map) using Samtools (v0.1.18) (4) and assembled to create transcript models by StringTie version-1.3.3 (5) guided by the reference genome (6). A consensus assembly was produced using StringTie merge. Transcripts with Fragments Per Kilobase of transcript per Million (FPKM) ≤ 0.2 were removed from downstream analyses. Details of all transcriptomic libraries generated for this study are listed in (**Suppl. Table S1**). GFF files of the transcriptome assembly were submitted to GEO (GSE129667, Transcriptome_Assembly_G006_V2.gff).

**Functional annotation and *M. persicae* lncRNAs identification**

The computational workflow for the lncRNAs identification of the *M. persicae* is shown in **Suppl. Fig. S2A**. lncRNA identification was performed on transcript models obtained from the transcriptome assembly described above. Mikado compare (7) was used to identify and subsequently exclude transcripts overlap over 10% of the length of annotated miRNA, tRNA, rRNA and transposons features. To identify putative Open Reading Frame (ORFs), we used TransDecoder (https://transdecoder.github.io) with the default parameters. To further maximize the sensitivity, we scanned all ORFs for homology to curated protein sequence (Arthropods) in the Swiss-Prot database (8), downloaded November 20, 2018 from http://www.uniprot.org/downloads. This was done by BlastP (v2.7.1, evalue 1e-5). HAMMER (v-3.0) was run against the Pfam database (9) with default parameter to search for protein domains. The coding potentials of the remaining transcripts were assessed using CPC2 (10) and those with a coding potential score < 0.5 were selected. To consider transcript as a candidate lncRNA, transcript must be larger than 200nt, not have a hit in the SwissProt, Pfam database, considered non-coding by the CPC2, and not be already classified as another class of functional RNA (rRNA, miRNA, tRNA, transposons). The candidate lncRNA were submitted as a separate gff file (GSE129667, Candidate_lncRNA.gff).

Functional annotation for the protein coding genes were generated using annotF v1.02 (https://github.com/EI-CoreBioinformatics/AnnotF) and was submitted to GEO (GSE129667, Functional_annotation.txt.gz). Assessments of whether *M. persicae* genes are expressed in the salivary glands and guts was done by performing blastn against the EST datasets of Ramsey et al., 2007 (28).

**Manual annotation of *Ya* genes**

A 148 bp nucleotide sequence (**Suppl. Fig. S7**) that includes the entire exon 2 of the *Ya* genes and that was found to be conserved among 23 lncRNA genes in the darkslateblue module was used to search the *M. persicae* reference genome G006 with Blastn (v2.22) using default parameters. Blast hits, with coverage more than 80 percent were converted to GFF and loaded to the Apollo browser (11) along with the annotated gene models (22) and all assembled transcripts herein before the merging step (see above). Gene models and corresponding transcripts that aligned to the 148-bp nucleotide sequence were selected and further curated by manually annotating the 3' ends of each of the transcripts based on the presence of a poly-A tail. The 5' ends were identified based on the most conserved sequence among all transcripts combined with existing RT-PCR data for *Ya1* (**Suppl. Fig. S11**). The curated annotation of the *Ya* family enabled more accurate transcript quantifications among its members. We updated the gene models in our previous gff file for the Ya locus and submitted as a separate file (GSE129667, Manual_Anno.V3.gff).

**Differential gene expression analysis**

Differentially expressed transcripts of *M. persicae* colonies on the 9-plant species was determined by comparing transcript expression levels of *M. persicae* colonies on *B. rapa* (original host) with those of colonies on the 8 other plant species (new hosts) (see experimental design in Fig. 1A) using the DESeq2 package in R (v1.2.10) (12) and transcript count per million (TPM) generated by Kallisto v0.42.3 (https://pachterlab.github.io/kallisto/) (**Suppl. Table S2**). Additional filtering was employed in DESeq2 to remove lowly expressed transcripts (mean count < 10) on the basis of normalized counts. Transcripts were considered differentially expressed if they had a *p* value less than

0.05 after accounting for a 5% FDR according to the Benjamini-Hochberg procedure and if log2Fold change was greater than 1.

**Co-expression analysis**

Weighted gene co-expression network analysis (WGCNA) was used to generate unsigned co-expression networks on nine host swap data (13). Genes with normalized count (TPM) > 5 in at least one sample were used for the co-expression analysis and clustered into network modules using the topological overlap measure (TOM). Genes were grouped by hierarchical clustering on the basis of dissimilarity of gene connectivity (1-TOM). The co-expression clusters were produced by cutreeDynamic in which the minimum size of modules was kept at 20 genes. The modules were randomly colour-labelled. An adjacency matrix was built by applying a power function ($\beta$) on the Pearson correlation matrix. A $\beta$ value of 18 was found to be optimal for balancing the scale-free property of the co-expression network and the sparsity of connections between genes.

**Identification *M. persicae* RNAs in plants**

To assess if aphid translocate transcripts into plants, twenty adult *M. persicae* were caged on rosette leaves of 4-week-old *A. thaliana* plants for 24 hrs. The caged leaf area with aphids was assigned ‘aphid feeding site’. Leaves on plants caged with empty cages were included as controls. Leaf areas covered by the cages were carefully washed three times with deionized water and three time with nuclease-free water. RNAs were isolated from four independent biological replicates of aphid-exposed leaves and non-exposed control leaves and processed for RNA-seq library synthesis and sequenced on the Ilumina HiSeq 25000 (Novogene, Beijing, China). Reads were trimmed to remove sequencing adapters and aligned to *A. thaliana* genome (TAIR10 database, http://arabidopsis.org) and the *M. persicae* G006 genome (22) with HISAT2 v2.0.5. Reads mapped to the *M. persicae* genome were retrieved and subjected to further filtering by mapping them back to the *A. thaliana* genome. Reads that did not align to the *A. thaliana* genome in the last step were considered as unique *M. persicae* mapping reads. Transcripts with TPM ≥ 50 in at least one sample and that were present in at least three samples were selected for further analysis.

**RT-PCR analyses to detect systemic migration of aphid transcripts in plants**

Systemic migration of aphid transcripts was determined by caging a leaf section with aphids and detection of aphid transcripts in the caged area (feeding site), next to the caged area of the same leaf (near-feeding site) and a systemic leaf (systemic site). See experimental setups shown in **Fig. 4A** and **Suppl. Fig. S12**. Plants exposed to cages without aphids were used as controls. For *A. thaliana* plants, aphids were caged at the distal halves of the 8$^{th}$ rosette leaf (14) of 4-week-old *A. thaliana* plants for 24 hrs. The proximal leaf area of the 8$^{th}$ leaf next to the cage near the petiole was assigned near-feeding site and the 5$^{th}$ leaf that is likely phloem-connected with the 8$^{th}$ leaf (14) of *A. thaliana* plants the systemic site. Similar setup as for *A. thaliana* were used for *B. rapa*, *P. sativum* and maize plants, except that for maize the near-systemic site was the middle of the leaf next to the caged area and systemic site the part of this leaf near the stem (**Suppl. Fig. S12**). Upon the 24 hr period of exposure to the cages with or without aphids, sections of leaves that were caged were immediately separated from the non-caged parts with scissors and the cages and aphids removed. Then the remaining parts of the leaves and systemic leaves were detached from the plants. Leaf sections detached from plants were cleaned with the brush to remove aphids and visible debris. The leaf tissues were submerged in 5 mL of MilliQ water in a 15 mL tube. The tube was shaken for 30 sec after which the MilliQ water was removed. This was repeated two more times with MilliQ water and three additional times with nuclease-free water. Samples were snap-frozen in liquid nitrogen and storage at -80 °C.

Total RNAs were isolated from the leaf tissues by RNeasy Plant Mini Kit (Qiagen, Germany) followed by a DNase treatment with RNase-free DNase I (Thermo Fisher Scientific, US). cDNA was synthesized from 1 µg total RNA at 20 µL reaction volume with poly(A) primers using the RevertAid First Strand cDNA Synthesis Kit (Thermo Fisher Scientific, US). The qRT-PCRs reactions were performed on a CFX96 Touch™ Real-Time PCR Detection System using transcript-specific primers. Each reaction was performed in a 20 µL reaction volume containing 10 µL SYBR Green (Maxima SYBR Green/ROX qPCR Master Mix, Thermo Fisher Scientific, USA), 0.4 µL Rox Reference Dye II, 1 µL of each primer (10 µM), 1 µL of 20 µL sample cDNA, and 7.6 µL UltraPure Distilled water (Invitrogen, US). The PCR cycles were: 95 $^{o}$C for 10 s, 40 cycles at 95 $^{o}$C for 20 s, 63 $^{o}$C for 30 s.

To compare *Ya1* transcript concentrations in the feeding and systemic sites, a standard *Ya1* concentration curve was generated (**Suppl. Fig. S13**). For this, the 273-nt Ya1 fragment cloned into the plasmid pBI121 under promoter 35S was used as the PCR template. The concentration curve was generated with a serial of dilution of the plasmid pBI121_35S::Ya1 from the highest concentration 1.44E$^{-10}$ g/ µL to the lowest concentration 1.44E$^{-18}$ g/ µL and primers (Ya1 primer6 and Ya1 primer9, **Suppl. Table S7**) using PCR conditions as described above.

**Primer design**

Primers to specifically amplify *M. persicae Ya* transcripts were designed with the PrimerQuest Tool (Integrated DNA Technologies, IA, USA) that predicted five to ten primers for each of the *Ya* transcripts. The primer pairs were aligned to the sequences of all *Ya* transcripts and the ones that matched unique sequences of one *Ya* transcript selected (**Suppl. Table S7**). *Ya* transcripts for which no unique primers were available were excluded from further analyses.

**Detection of aphid transcripts by northern blotting**

For northern blot analyses, 1 ug of total plant and aphid RNA were separated on a 6% denaturing polyacrylamide gel (PAGE) with 1 X TBE buffer (10 X TBE buffer stock, Thermo Fisher Scientific, USA) at 100 V for 60 min. The 273-nt *Ya1* fragment was cloned with a SP6 sequence at the 3 the end of *Ya1* and used to synthesize a 291-nt *Ya1-SP6* RNA of which 100 ng was ran alongside the total RNAs from aphids and plants on polyacrylamide gels. RNAs were transferred to a nylon membrane (Hybond N, Amersham, UK) by electroblotting at 0.8 A for 2 hrs

(BIO-RAD, USA) using 0.5 X TBE buffer at 60 V for 60 min. RNAs were cross-linked to the membranes by a UV cross linker (UVP Inc., CA, USA) using auto-crosslink function, twice on the side of the blot exposed to the gel and one time on the other slide of the blot.

*Ya1* transcript in aphids were detected via hybridization of the northern blots to a biotin-labelled anti-sense sequence of *Ya1*. Biotin-labelled anti-sense probe of *Ya1* was synthesized from anti-sense sequences of *Ya1* with the MAXIscript™ SP6/T7 Transcription Kit (Thermo Fisher Scientific, USA) and Biotin-16-dUTP (Roche, USA). The northern blots were performed according to the manufacturing manual of North2South Chemiluminescent Hybridization and Detection Kit (Thermo Fisher Scientific, US). Briefly, the blot was washed in North2South Hybridization Stringency Wash Buffer at room temperature for one time 20 min, in Wash Buffer containing 2X SSC/0.1% SDS at room temperature for three times 20 min, and Stringency Wash Buffer at 65 $^{o}$C for one time 20 min. The washing was done in 0.2 ml per cm$^2$ of blot. The blot was then incubated in the Substrate Working Solution containing equal volumes of the Luminol/Enhancer Solution and Stable Peroxide Solution for 5 min. To visualize the hybridization signal, the membrane was exposed to an X-ray film for an appropriate exposure time.

*Ya1* transcript in plants were detected via hybridization of the northern blots to a 273-nt *Ya1* fragment that was labeled using 3000 Ci/mmol of [a-32P] dATP (PerkinElmer Life Sciences. USA) with the Klenow DNA polymerase reaction as per manufacturer's instruction (Megaprime DNA Labeling System, GE Healthcare).The radioactively-labelled PCR probe was denatured at 95ºC for 5 min, transferred to ice and then incubated with the blot in 50 ml Hybridization buffer (Sigma-Aldrich) at 42°C for overnight. Washing was done three times at 42ºC with washing buffer (2X SSC/0.1% SDS). The membranes were exposed to storage Phosphor Screens (GE Healthcare) and hybridization signals were visualized using Typhoon Trio (GE Healthcare) scanner.

**Sequencing of 3' ends of aphid *Ya1* transcript**

Total RNA was extracted from *M. persicae* aphids reared on *B. rapa* and 3 ug of aphid total RNAs was added to a 80 μL ligation mixture containing 8 μL T4 RNA ligase buffer and 4 μL T4 RNA ligase (Thermo Fisher Scientific, USA), 8 μL ATP (Invitrogen, USA), 8 μL BSA (Invitrogen, USA), and 10 pmol 3' RACE RNA adaptor (**Suppl. Table S7**). RNA ligation was carried out at 16 °C overnight. The ligated RNA was converted into cDNA using oligo sequences complementary to the 3' RACE adaptor with RevertAid First Strand cDNA Synthesis Kit (Thermo Fisher Scientific, USA) (**Suppl. Table S7**). RACE PCRs were performed with the Ya1 forward primer GGACAAGTCCAATCTGC and the adapter primer CAAGCAGAAGACGGCATACGA (**Suppl. Table S7**) in a 50 μL reaction volume containing 0.5 μL Phusion DNA polymerase (NEB), 10 μL 5X Phusion HF buffer, 1 μL 10 mM dNTPs, 1 μL of each primer (10 μM), 1 μL of cDNA sample. The cycle programs were: 98 °C for 30 s, 35 cycles at 98 °C for 10 s, 60 °C for 30 s, 72 °C for 15 s, final extension 72 °C 10 mins. PCR produces were separated by 3% agarose gel. DNA bands were visualized under UV light, then cut to extract DNA using QIAquick Gel Extraction Kit (Qiagen, Germany). DNA were ligated to pGEM-T (Promega, USA) and sequenced with M13 primer.

**Plasmid construction**

To generate pJawohl8-RNAi constructs, a 273 bp of Ya1 was amplified from *M. persicae* cDNA by PCR with specific primers containing additional attB1 and attB2 linkers (**Suppl. Table S7**) for cloning with the Gateway system (Invitrogen, USA). The 273-bp Ya1 fragment was introduced into pDONR207 (Invitrogen, USA) plasmid using Gateway BP reaction and transformed into DH5α competent cells (Invitrogen, USA). Subsequent clones were sequenced to verify correct size and sequence of inserts. Via the Gateway LB reaction, inserts were transferred from pDONR207 into the plant transformation vector, pJawohl8-RNAi (kindly provided by I.E. Somssich, Max Planck Institute for Plant Breeding Research, Germany), which is a plasmid that enables the expression of the transgene as a double-stranded hairpin transcript, generating plasmid pJawohl8-RNAi_Ya1.

Plasmids pBI121_35S::Ya1 and pBI121_35S::Ya1_3ATGs_mut were constructed as follows. The fragment corresponding to the 273 nt Ya1 transcript was amplified from *M. persicae* cDNA by PCR with specific primers containing *Bam*HI and *Sac*I restriction sites in the forward and reverse primers, specifically (**Suppl. Table S7**). The *Bam*HI and *Sac*I digested PCR fragments were introduced into pBI121 to generate pBI121_35S::Ya1. Plasmid pBI121_35S::Ya1 was used in overlap PCR reactions (15) to generate the Ya1_3ATGs_mut construct in which the three ATGs at positions 41, 62, and 146 of Ya1 were converted into TAG stop codons. This involved the amplification of fragments 1, 2, 3 and 4 with primer pairs MutateF and ATG41_R, ATG41_F and ATG62_R, ATG62_F and ATG146_R and ATG146_F and MutateR, respectively (see **Suppl. Table S7 for primer sequences**), and subsequent amplification of a 81 bp fragment from fragments 1 and 2 with primers MutateF and ATG62_R, a 230 bp fragment from fragments 3 and 4 with primers ATG62_F and MutateR, and finally the amplification of the 273 nt Ya1 mutant in which three ATGs were mutated to three TAGs by combining the 81 bp and 230 bp fragments and primers MutateF and MutateR. The resulting Ya1_3ATGs_mut PCR fragment was digested with *Bam*HI and *Sac*I and introduced into pBI121 to generate pBI121_35S::Ya1_3ATGs_mut.

All plasmid inserts were sequenced for verification.

**Generation of transgenic *A. thaliana* plants**

pJawohl8-RNAi_Ya1, pBI121_35S::Ya1, and pBI121_35S::Ya1_3ATGs_mut plasmids were introduced into *A. tumefaciens* strain GV3101 that carried the helper plasmid pMP90RK for subsequent transformation of *A. thaliana* Col-0 using the floral dip method (Bechtold et al., 1993). Transgenic seeds were selected on Murashige and Skoog (MS) medium supplemented with 20 μg/mL phosphinothricin (BASTA) to select dsRNA_Ya1 transformants or on MS containing 50 μg/mL kanamycin to selection of 35S::Ya1 and 35S::Ya1_3ATGs_mut transformants. F2 seeds were germinated on MS medium supplemented with 20 μg/mL BASTA or 50 μg/mL kanamycin for selection. F2 seedlings with 3:1 alive/dead segregation (evidence of single insertion) were taken forward to the F3 stage. Seeds from F3 plants were sown on MS with BASTA (for dsRNA_Ya1 plants) or on MS with kanamycin (for 35S::Ya1 and 35S::Ya1_3ATGs_mut) and lines with 100% survival ratio (homozygous) were selected.

**Knock down of aphid transcripts by plant-mediated RNA interference (RNAi)**

Seeds of the dsRNA_Ya1 homozygous lines (expressing dsRNA corresponding to *Ya1*) were sown on MS medium and, after one week, seedlings were transferred to single pots (8 cm diameter) and transferred to a controlled environmental growth room at temperature 24 $^0$C day/20 $^0$C night under 10 hours of light. One *M. persicae* adult was confined to single 4-weeks-old Arabidopsis lines in sealed experimental cages (15.5 cm diameter and 30 cm height) containing the entire plant. One day later, the adult was removed, five nymphs remained on the plants and become adults in five days. Adults were harvested for qRT-PCR to confirm *Ya1* silencing compared to adults on dsGFP plants.

For qRT-PCRs, total RNA was isolated from aphids using Trizol reagent (Sigma) and subsequent DNase treatment using an RNase-free DNase I (Thermo Fisher Scientific, US). cDNA was synthesized from 1 μg total RNA with RevertAid First Strand cDNA Synthesis Kit (Thermo Fisher Scientific, US). The qRT-PCRs reactions were performed on CFX96 Touch™ Real-Time PCR Detection System using gene-specific primers. Each reaction was performed in a 20 μL reaction volume containing 10 μL SYBR Green (Thermo Fisher Scientific, US), 0.4 μL Rox Reference Dye II, 1 μL of each primer (10 μM), 1 μL of sample cDNA, and 7.6 μL UltraPure Distilled water (Invitrogen, US). The cycle programs were: 95 $^o$C for 10 s, 40 cycles at 95 $^o$C for 20 s, 60 $^o$C for 30 s. Relative quantification was calculated using the comparative $2^{-\triangle}$Ct method (16). All data were normalized to the level of elongation factor gene (MYZPE13164_G006_v1.0_000087220) from the same sample.

***M. persicae* fecundity assay**

Seeds of Arabidopsis lines were sown on MS medium and, after one week, seedlings were transferred to single pots (8 cm diameter) and transferred to a controlled environmental growth room at temperature 24 $^{0}$C day/20 $^{0}$C night under 10 hours of light. One *M. persicae* adult was confined to single 4-weeks-old Arabidopsis lines in sealed experimental cages that contained the entire plant. One day later, the adult was removed, and one nymph remained on the plants. Offspring produced on the 7$^{th}$, 9$^{th}$, 11$^{th}$ day of the experiment were scored and removed. This experiment was repeated three times to create data from three independent biological replicates with four to six plants per line per replicate.

**Statistical analyses**

All the data analyses were performed in R (v3.5.2). All statistical tests are described in the figure legends.
