## Supplementary material for "An aphid host-responsive RNA transcript that migrates systemically in plants promotes aphid reproduction": Suppl. Fig. S1

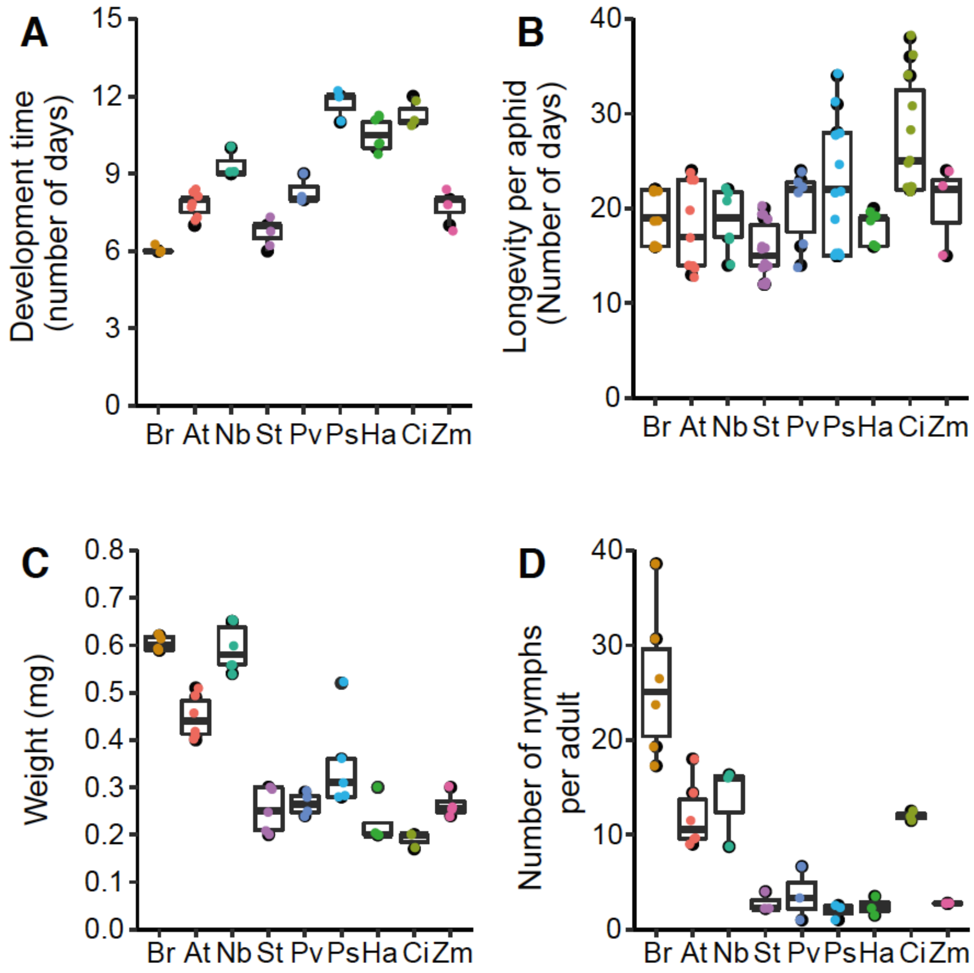


**Suppl. Fig. S1.** Performance parameters of stable colonies of *M. persicae* clone O on 9 divergent plant species. Colonies on *B. rapa* (Br), *A. thaliana* (At), *N. benthamiana* (Nb), *S. tuberosum* (St), *C. indicum* (Ci), *H. annuus* (Ha), *P. sativum* (Ps), *P. vulgaris* (Pv) and *Zea mays* (Zm), established as shown in **Fig. 1A**, were analysed for development time (A) in days between birth and emergence of adults (n = 3-7 plants with one aphid each), longevity (B) in days between birth to death (n = 3-15 plants with one aphid each), weight (C) in mg per adult (n = 3-6 plants with 100 adult aphids each), and number of nymphs per adult (D) calculated by dividing the number of nymphs by the number of adults on days 7, 9 and 11 (Br, At, Nb and St) or days 7, 9, 11, 13, 15, 17 and 18 (Ci, Ha, Ps, Pv and Zm) (n = 3-6 plants seeded with one aphid each).
