## Supplementary material for "An aphid host-responsive RNA transcript that migrates systemically in plants promotes aphid reproduction": Suppl. Fig. S2

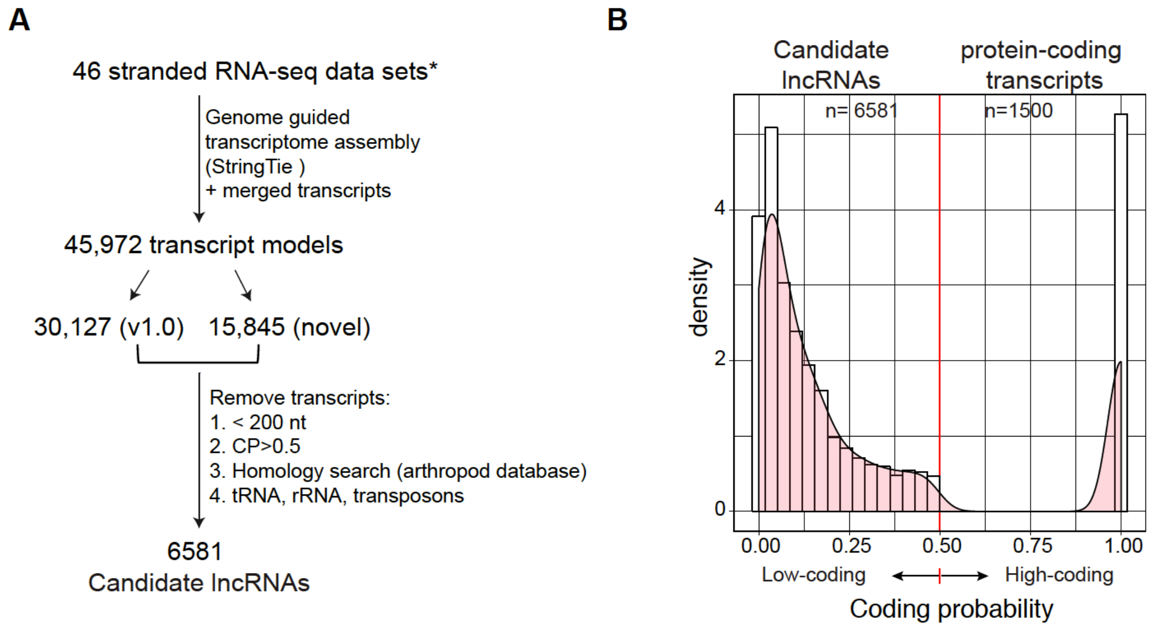


**Suppl. Fig. S2.** Annotation of *M. persicae* transcript models and candidate lncRNAs. (A) Transcript annotation pipeline. RNA-seq reads from 45 libraries generated from *M. persicae* on nine divergent hosts (data generated herein, **Fig. 1A**) and LIB1771 derived from a *M. persicae* colony on *B. rapa* (1) were assembled into transcripts using a genome-guided approach with StringTie (2). Of the 45 972 transcripts in total, 30 127 were annotated previously (1) and 15 845 are novel. The 6 581 transcripts that are candidate lncRNAs were identified upon removing transcripts of < 200 nt in size, transcripts with coding potential (CP) >0.5, as determined using the Coding Potential Calculator 2 (CPC2, http://cpc2.cbi.pku.edu.cn/) (3), and that have similarities of deduced protein sequences to known arthropods proteins, house-keeping RNAs (rRNA and tRNA) and transposons. (B) Distribution of CPC2 coding probability scores of 6 581 candidate lncRNAs identified in Fig. S2A and 1 500 randomly selected transcripts from *M. persicae* protein-coding sequences.
