## Supplementary material for "An aphid host-responsive RNA transcript that migrates systemically in plants promotes aphid reproduction": Suppl. Fig. S3

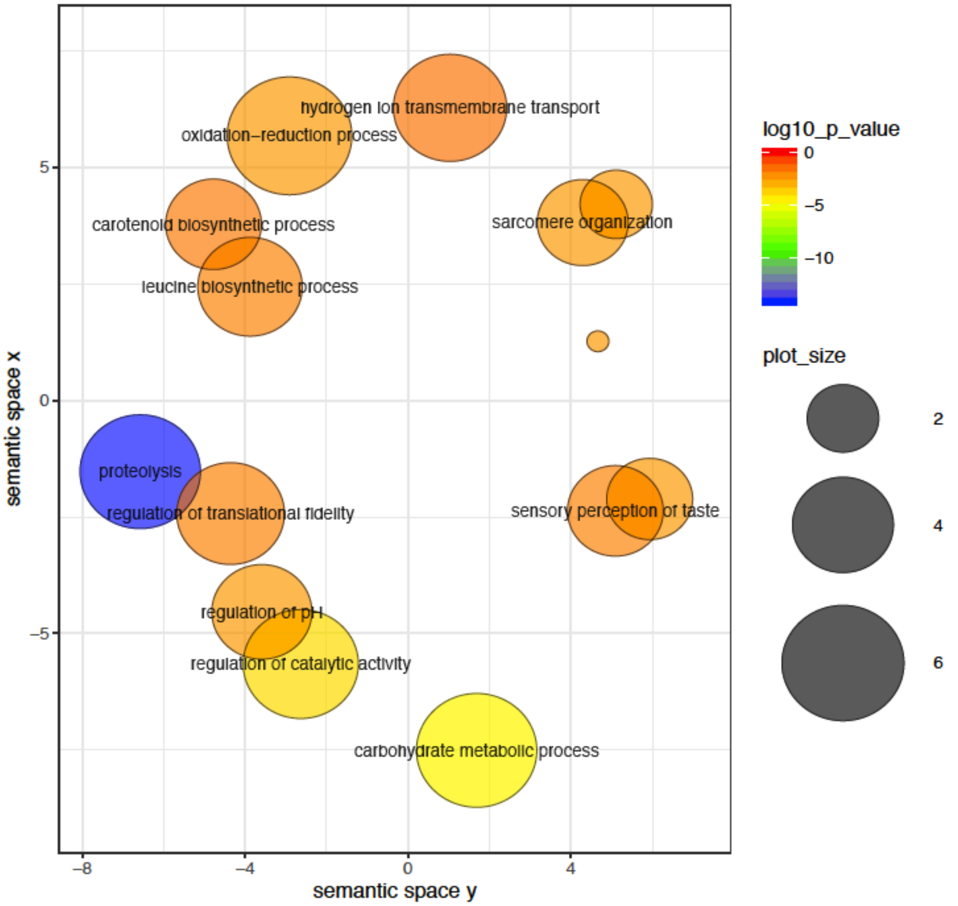


**Suppl. Fig. S3**. Biological process GO terms that are enriched among DE genes of *M. persicae* on 9 plants species. Graphs was generated from data shown in **Fig. 1C**. Enriched GO terms are represented by circles and are clustered according to semantic similarities to other GO terms in the gene ontology. Distance between the circles indicates the semantic similarity between the corresponding GO terms. Circle size is proportional to the frequency of the GO term, whereas colour indicates the $\log_{10}$(*p* value) for the enrichment calculated using GO-seq (red higher, blue lower). Only GO-terms having *p* value and padj < 0.05 are shown. Data were summarized and visualized by REVIGO online tool (http://revigo.irb.hr/) (1).
