## Supplementary material for "An aphid host-responsive RNA transcript that migrates systemically in plants promotes aphid reproduction": Suppl. Fig. S4

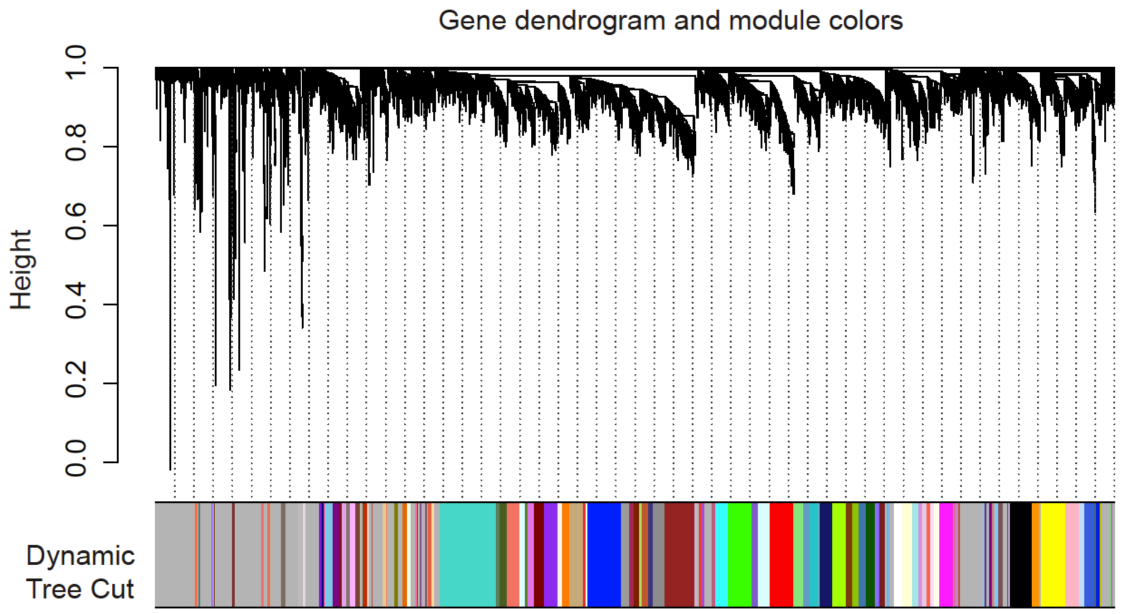


**Suppl. Fig. S4**. WGCNA gene co-expression modules of *M. persicae* on 9 plant species. The dendogram groups 11 824 *M. persicae* genes into 77 distinct modules (color-coded) using hierarchical clustering. Dynamic cut tree was used to identify modules, dividing modules at significant branch points in the dendrogram. The x-axis shows the width of each module that is defined by the number of genes. The y-axis corresponds to distance determined by the extent of topological overlap (1-TOM). Genes not assigned to a module are labelled in grey.
