## Supplementary material for "An aphid host-responsive RNA transcript that migrates systemically in plants promotes aphid reproduction": Suppl. Fig. S5

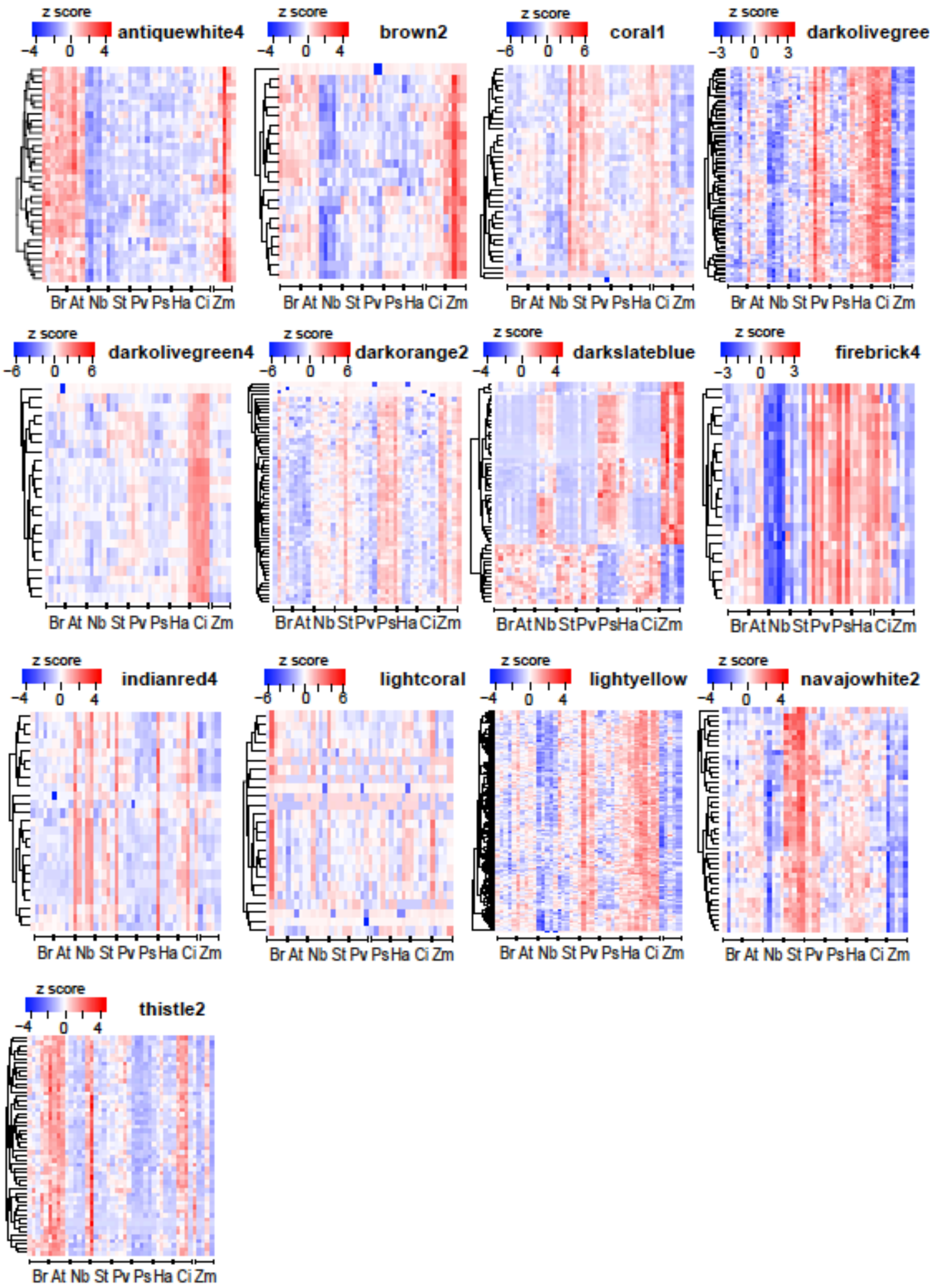


**Suppl. Fig. S5.** Heatmaps of 13 modules enriched for DE genes of *M. persicae* colonies on 9 plant species. The colour codes of the modules are indicated above the heatmaps. Hierarchical clustering was done based on plant species as follows: Br, *B. rapa*; At, *A. thaliana*; Nb, *N. benthamiana*; St, *S. tuberosum*; Ci, *C. indicum*; Ha, *H. annuus*; Ps, *P. sativum*; Pv, *P. vulgaris*; Zm, *Z. mays*. Rows are scaled based log transformed TPM values, which are shown using a z-score as indicated.
