## Supplementary material for "An aphid host-responsive RNA transcript that migrates systemically in plants promotes aphid reproduction": Suppl. Fig. S6

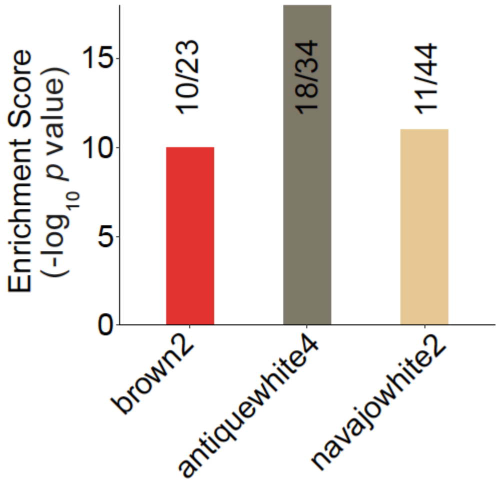


**Suppl. Fig. S6**. Modules enriched for genes expressed in *M. persicae* guts. *M. persicae* DE genes were searched against a gut EST dataset (1). The x/y numbers above the bars indicate the number of genes in the enriched category (x) and the total number of genes in the module (y).
