## Supplementary material for "An aphid host-responsive RNA transcript that migrates systemically in plants promotes aphid reproduction": Suppl. Fig. S7

```
>Ya1_exon2_148nt
TTTCTCTTTTAAACCTAAAAAACCAACCAACAAATCAAAAATGGGCGCTGAAAAGGTATCCATGAACA
TCGTTGTCGTCGGACAAGTCCAATCTGCCAAGGCCATCAAGACCGTCGCCAAGGCCTCCCCAGCCACC
AACCAATTGATG
```

**Suppl. Fig. S7.** Nucleotide sequence of *Ya1* exon 2 used to identify *Ya* family members in *M. persicae* and other aphid species**.**
