## Supplementary material for "An aphid host-responsive RNA transcript that migrates systemically in plants promotes aphid reproduction": Suppl. Fig. S8

See enclosed supplementary Fig. S8.

**Suppl. Fig. S8.** Alignment of transcripts of 30 *Ya* gene family members. The alignment was used to generate Fig. 3A and the phylogenetic tree of Fig. 3B. Alignment was performed on online version of MAFFT (version 7, https://mafft.cbrc.jp/alignment/server/index.html) (1) with default settings. The multiple alignment was subjected to GUIDANCE2 (http://guidance.tau.ac.il/ver2/) (2) to compute the residue-wise confidence scores and extract well-aligned residues. Nucleotides are highlighted (colour scale between red and turquoise) to indicate confidence of alignment (red is high confidence and turquoise is low). The bottom bar graph displays the scores of aligned nucleotides, as determined by GUIDANCE2.
