## Supplementary Fig. S8 for "An aphid host-responsive RNA transcript that migrates systemically in plants promotes aphid reproduction"

### MSA color-coded by GUIDANCE scores

|  |  |  |  |  |  |  |  |  |  |  |  |  |  |  |  |  |  |  |  |  |  |  |  |  |  |  |  |  |  |  |  |  |  |  |  |  |  |  |  |  |  |  |  |  |  |  |  |  |  |  |  |  |  |  |  |  |  |  |  |  |  |  |  |  |  |  |  |  |  |  |  |  |  |  |  |  |  |  |  |  |  |  |  |  |  |  |  |  |  |  |  |  |  |  |  |  |  |  |  |  |  |  |  |  |  |  |  |  |  |  |  |  |  |  |  |  |  |  |  |  |  |  |  |  |  |  |  |  |  |  |  |  |  |  |  |  |  |  |  |  |  |  |  |  |  |  |  |  |  |  |  |  |  |  |  |  |  |  |  |  |  |  |  |  |  |  |  |  |  |  |  |  |  |  |  |  |  |  |  |  |  |  |  |  |  |  |  |  |  |  |  |  |  |  |  |  |  |  |  |  |  |  |  |  |  |  |  |  |  |  |  |  |  |  |  |  |  |  |  |  |  |  |  |  |  |  |  |  |  |  |  |  |  |  |  |  |  |  |  |  |  |  |  |  |  |  |  |  |  |  |  |  |  |  |  |  |  |  |  |  |  |  |  |  |  |  |  |  |  |  |  |  |  |  |  |  |  |  |  |  |  |  |  |  |  |  |  |  |  |  |  |  |  |  |  |  |  |  |  |  |  |  |  |  |  |  |  |  |  |  |  |  |  |  |  |  |  |  |  |  |  |  |  |  |  |  |  |  |  |  |  |  |  |  |  |  |  |  |  |  |  |  |  |  |  |  |  |  |  |  |  |  |  |  |  |  |  |  |  |  |  |  |  |  |  |  |  |  |  |  |  |  |  |  |  |  |  |  |  |  |  |  |  |  |  |  |  |  |  |  |  |  |  |  |  |  |  |  |  |  |  |  |  |  |  |  |  |  |  |  |  |  |  |  |  |  |  |  |  |  |  |  |  |  |  |  |  |  |  |  |  |  |  |  |  |  |  |  |  |  |  |  |  |  |  |  |  |  |  |  |  |  |  |  |  |  |  |  |  |  |  |  |  |  |  |  |  |  |  |
| --- | --- | --- | --- | --- | --- | --- | --- | --- | --- | --- | --- | --- | --- | --- | --- | --- | --- | --- | --- | --- | --- | --- | --- | --- | --- | --- | --- | --- | --- | --- | --- | --- | --- | --- | --- | --- | --- | --- | --- | --- | --- | --- | --- | --- | --- | --- | --- | --- | --- | --- | --- | --- | --- | --- | --- | --- | --- | --- | --- | --- | --- | --- | --- | --- | --- | --- | --- | --- | --- | --- | --- | --- | --- | --- | --- | --- | --- | --- | --- | --- | --- | --- | --- | --- | --- | --- | --- | --- | --- | --- | --- | --- | --- | --- | --- | --- | --- | --- | --- | --- | --- | --- | --- | --- | --- | --- | --- | --- | --- | --- | --- | --- | --- | --- | --- | --- | --- | --- | --- | --- | --- | --- | --- | --- | --- | --- | --- | --- | --- | --- | --- | --- | --- | --- | --- | --- | --- | --- | --- | --- | --- | --- | --- | --- | --- | --- | --- | --- | --- | --- | --- | --- | --- | --- | --- | --- | --- | --- | --- | --- | --- | --- | --- | --- | --- | --- | --- | --- | --- | --- | --- | --- | --- | --- | --- | --- | --- | --- | --- | --- | --- | --- | --- | --- | --- | --- | --- | --- | --- | --- | --- | --- | --- | --- | --- | --- | --- | --- | --- | --- | --- | --- | --- | --- | --- | --- | --- | --- | --- | --- | --- | --- | --- | --- | --- | --- | --- | --- | --- | --- | --- | --- | --- | --- | --- | --- | --- | --- | --- | --- | --- | --- | --- | --- | --- | --- | --- | --- | --- | --- | --- | --- | --- | --- | --- | --- | --- | --- | --- | --- | --- | --- | --- | --- | --- | --- | --- | --- | --- | --- | --- | --- | --- | --- | --- | --- | --- | --- | --- | --- | --- | --- | --- | --- | --- | --- | --- | --- | --- | --- | --- | --- | --- | --- | --- | --- | --- | --- | --- | --- | --- | --- | --- | --- | --- | --- | --- | --- | --- | --- | --- | --- | --- | --- | --- | --- | --- | --- | --- | --- | --- | --- | --- | --- | --- | --- | --- | --- | --- | --- | --- | --- | --- | --- | --- | --- | --- | --- | --- | --- | --- | --- | --- | --- | --- | --- | --- | --- | --- | --- | --- | --- | --- | --- | --- | --- | --- | --- | --- | --- | --- | --- | --- | --- | --- | --- | --- | --- | --- | --- | --- | --- | --- | --- | --- | --- | --- | --- | --- | --- | --- | --- | --- | --- | --- | --- | --- | --- | --- | --- | --- | --- | --- | --- | --- | --- | --- | --- | --- | --- | --- | --- | --- | --- | --- | --- | --- | --- | --- | --- | --- | --- | --- | --- | --- | --- | --- | --- | --- | --- | --- | --- | --- | --- | --- | --- | --- | --- | --- | --- | --- | --- | --- | --- | --- | --- | --- | --- | --- | --- | --- | --- | --- | --- | --- | --- | --- | --- | --- | --- | --- | --- | --- | --- | --- | --- | --- | --- | --- | --- | --- | --- | --- | --- | --- | --- | --- | --- | --- | --- | --- | --- | --- | --- | --- | --- | --- | --- | --- |
|  |  | 1 |  |  |  |  |  |  |  |  | 1 | 0 |  |  |  |  |  |  |  |  | 2 | 0 |  |  |  |  |  |  |  |  | 3 | 0 |  |  |  |  |  |  |  |  | 4 | 0 |  |  |  |  |  |  |  |  | 5 | 0 |  |  |  |  |  |  |  |  | 6 | 0 |  |  |  |  |  |  |  |  | 7 | 0 |  |  |  |  |  |  |  |  | 8 | 0 |  |  |  |  |  |  |  |  | 9 | 0 |  |  |  |  |  |  |  |  | 1 | 0 | 0 |  |  |  |  |  |  |  | 1 | 1 | 0 |  |  |  |  |  |  |  | 1 | 2 | 0 |  |  |  |  |  |  |  | 1 | 3 | 0 |  |  |  |  |  |  |  | 1 | 4 | 0 |  |  |  |  |  |  |  | 1 | 5 | 0 |  |  |  |  |  |  |  | 1 | 6 | 0 |  |  |  |  |  |  |  | 1 | 7 | 0 |  |  |  |  |  |  |  | 1 | 8 | 0 |  |  |  |  |  |  |  | 1 | 9 | 0 |  |  |  |  |  |  |  | 2 | 0 | 0 |  |  |  |  |  |  |  | 2 | 1 | 0 |  |  |  |  |  |  |  | 2 | 2 | 0 |  |  |  |  |  |  |  | 2 | 3 | 0 |  |  |  |  |  |  |  | 2 | 4 | 0 |  |  |  |  |  |  |  | 2 | 5 | 0 |  |  |  |  |  |  |  | 2 | 6 | 0 |  |  |  |  |  |  |  | 2 | 7 | 0 |  |  |  |  |  |  |  | 2 | 8 | 0 |  |  |  |  |  |  |  | 2 | 9 | 0 |  |  |  |  |  |  |  | 3 | 0 | 0 |  |  |  |  |  |  |  | 3 | 1 | 0 |  |  |  |  |  |  |  | 3 | 2 | 0 |  |  |  |  |  |  |  | 3 | 3 | 0 |  |  |  |  |  |  |  | 3 | 4 | 0 |  |  |  |  |  |  |  | 3 | 5 | 0 |  |  |  |  |  |  |  | 3 | 6 | 0 |  |  |  |  |  |  |  | 3 | 7 | 0 |  |  |  |  |  |  |  | 3 | 8 | 0 |  |  |  |  |  |  |  | 3 | 9 | 0 |  |  |  |  |  |  |  | 4 | 0 | 0 |  |  |  |  |  |  |  | 4 | 1 | 0 |  |  |  |  |  |  |  | 4 | 2 | 0 |  |  |  |  |  |  |  | 4 | 3 | 0 |  |  |  |  |  |  |  | 4 | 4 | 0 |  |  |  |  |  |  |  | 4 | 5 | 0 |  |  |  |  |  |  |  | 4 | 6 | 0 |  |  |  |  |  |  |
|  | MYZPE13164\_G006\_V2.0\_0000 | A | T | C | A | C | G | C | C | A | C | A | C | - | - | - | - | - | - | - | - | - | - | - | - | - | G | C | C | A | C | C | - | - | - | - | - | T | T | C | C | C | C | C | T | T | T | C | T | A | C | T | - | - | - | - | - | - | - | - | - | G | G | C | T | G | A | C | G | A | A | A | A | A | G | T | A | T | A | T | - | A | A | G | G | C | G | C | C | C | G | A | G | A | T | C | A | C | G | A | T | T | C | C | G | G | C | A | T | A | A | C | A | C | A | T | - | T | T | T | T | T | G | T | G | C | A | G | A | T | C | T | A | T | T | C | T | T | T | G | T | T | T | T | - | - | - | - | - | - | C | A | C | - | T | T | T | A | T | T | T | - | - | C | A | C | A | A | A | C | A | A | A | A | - | - | - | A | C | C | A | A | C | A | A | A | T | C | A | A | A | A | A | T | G | G | G | C | G | C | T | G | A | A | A | A | A | G | T | A | A | C | C | A | T | G | A | A | C | A | T | C | G | T | C | G | T | C | G | T | C | G | G | A | C | A | G | G | T | C | C | A | A | T | C | T | G | C | C | A | A | G | G | C | C | A | T | C | A | A | A | T | C | G | G | T | C | G | C | C | A | A | A | G | C | C | A | C | C | C | C | A | G | C | C | A | C | C | A | G | C | C | A | A | T | T | G | A | T | G | A | A | C | T | G | T | T | A | A | T | T | - | - | - | T | A | T | G | T | A | G | T | C | - | - | C | A | T | C | C | A | - | - | - | - | - | - | - | - | - | T | C | A | C | A | A | A | C | T | C | A | A | A | A | A | T | G | C | C | A | T | A | G | C | C | C | G | A | A | C | A | G | T | A | C | T | A | T | - | - | - | - | - | - | - | - | - | T | A | T | A | T | T | G | T | A | A | T | A | A | T | T | C | - | - | - | - | - | - | - | - | - | - | - | - | - | - | - | - | - | - | - | A | T | A | T | G | A | G | T | A | T | T | G | A | C | A | C | C | G | C | A | A | A | A | T | A | A | A | T | C | A | G | A | T | A | T | T | T | - | - | - | - | - | - | - | - | - | - |
|  | MYZPE13164\_G006\_V2.0\_0000 | A | T | C | A | C | A | C | C | A | C | - | - | - | - | - | - | - | - | - | - | - | - | - | - | - | G | C | C | A | C | C | - | - | - | - | - | A | C | C | C | C | A | C | T | T | T | A | A | A | T | C | - | - | - | - | - | - | - | - | - | G | G | T | C | G | G | T | G | G | C | A | G | T | G | T | A | T | A | - | A | A | A | G | G | T | T | T | C | C | G | A | G | A | T | C | T | C | G | A | T | T | C | C | G | G | C | A | T | A | A | C | A | C | A | T | C | C | T | T | T | T | G | T | G | C | A | G | A | A | C | T | A | T | T | A | T | T | T | G | T | T | T | T | - | - | - | - | - | - | C | A | C | - | T | T | T | A | A | C | C | - | - | C | T | C | A | A | A | A | A | C | C | A | A | C | C | A | C | C | A | A | - | A | A | A | C | C | A | A | A | A | A | T | G | G | G | C | G | C | A | G | A | A | A | A | A | G | T | A | T | C | C | A | T | G | A | A | C | A | T | C | G | T | C | G | T | C | G | T | C | G | G | A | C | A | A | G | T | C | C | A | A | T | C | T | G | C | C | A | A | G | G | C | C | A | T | C | A | A | A | T | C | G | G | T | C | G | C | C | A | A | A | G | C | C | T | C | T | C | C | A | G | C | C | A | C | C | A | G | C | C | A | A | T | T | G | A | T | G | A | A | C | T | G | T | T | A | A | T | T | - | - | - | C | A | T | G | T | A | G | T | C | - | - | C | A | T | A | C | A | T | - | - | - | - | - | - | - | - | T | G | C | A | A | A | A | C | T | C | A | A | A | A | A | T | G | C | C | A | T | A | G | C | C | T | A | A | G | C | A | A | T | A | C | A | A | T | - | - | - | - | - | - | - | - | - | A | A | T | A | T | - | - | - | - | G | T | C | A | T | T | A | - | - | - | - | - | - | - | - | - | - | - | - | - | - | - | - | - | - | - | A | T | A | T | G | A | G | T | A | T | T | G | A | C | A | C | C | G | C | A | A | A | A | T | A | A | A | C | C | A | G | T | C | A | T | T | T | - | - | - | - | - | - | - | - | - | - |
|  | MYZPE13164\_G006\_V2.0\_0000 | A | T | C | A | C | A | C | C | A | C | - | - | - | - | - | - | - | - | - | - | - | - | - | - | - | G | C | C | A | C | T | - | - | - | - | - | G | C | C | C | C | A | C | T | T | T | A | A | A | T | C | - | - | - | - | - | - | - | - | - | G | G | T | C | G | G | T | G | G | C | A | G | T | G | T | A | T | A | T | A | A | A | G | G | T | T | T | C | C | A | A | G | A | T | C | T | C | G | A | T | T | C | C | G | G | C | A | T | A | A | C | A | T | A | T | C | C | T | T | T | T | G | T | G | C | A | G | A | A | C | T | A | T | T | A | T | T | T | G | T | T | T | C | - | - | - | - | - | - | C | A | C | - | T | T | T | A | A | T | C | - | - | C | T | C | A | A | A | A | A | C | C | A | A | C | C | A | C | C | A | A | - | A | A | A | C | C | A | A | A | A | A | T | G | G | G | C | G | C | A | G | A | A | A | A | A | G | T | A | T | C | C | A | T | G | A | A | T | A | T | C | G | T | C | G | T | C | G | T | C | G | G | A | C | A | A | G | T | C | C | A | A | T | C | T | G | C | C | A | A | G | G | C | C | A | T | C | A | A | A | T | C | G | G | T | C | G | C | C | A | A | A | G | C | C | A | C | C | C | C | A | G | C | A | A | C | C | A | G | C | C | A | A | T | T | G | A | T | G | A | A | C | T | G | T | T | A | A | T | T | - | - | - | C | A | T | G | A | A | G | T | C | - | - | C | A | T | A | C | A | T | - | - | - | - | - | - | - | - | T | G | C | A | A | A | A | C | T | C | A | A | A | A | A | T | G | C | C | A | T | A | G | C | C | C | A | A | A | C | A | G | T | A | C | A | A | T | - | - | - | - | - | - | - | - | - | A | A | T | A | T | - | - | - | - | G | T | A | A | T | T | C | - | - | - | - | - | - | - | - | - | - | - | - | - | - | - | - | - | - | - | A | T | A | T | T | T | G | T | G | T | T | G | A | C | A | C | C | T | A | A | A | A | A | T | A | A | A | T | C | A | G | T | - | - | - | - | - | - | - | - | - | - | - | - | - | - | - |
|  | MYZPE13164\_G006\_V2.0\_0001 | A | T | C | A | C | A | C | C | A | C | - | - | - | - | - | - | - | - | - | - | - | - | - | - | - | G | C | C | A | C | T | - | - | - | - | - | G | C | C | C | C | A | C | T | T | T | A | A | A | T | C | - | - | - | - | - | - | - | - | - | G | G | T | C | G | G | T | G | G | C | A | G | T | G | T | A | T | A | - | A | A | A | G | G | T | T | T | C | C | G | A | G | A | T | C | T | C | G | A | T | T | C | C | G | G | C | A | T | A | A | C | A | C | A | T | C | C | T | T | T | T | G | T | G | A | C | G | A | A | C | T | A | T | T | C | T | T | T | G | T | T | T | T | - | - | - | - | - | - | C | A | C | - | T | T | T | A | A | C | C | - | - | C | T | C | A | A | A | A | A | C | C | A | A | C | C | A | C | C | A | A | - | A | A | A | C | C | A | A | A | A | A | T | G | G | G | C | G | C | A | G | A | A | A | A | A | G | T | A | T | C | C | A | T | G | A | A | C | A | T | C | G | T | C | G | T | C | G | T | C | G | G | A | C | A | A | G | T | C | C | A | A | T | C | T | G | C | C | A | A | G | G | C | C | A | T | C | A | A | A | T | C | G | G | T | C | G | C | C | A | A | A | G | C | C | A | C | T | C | C | A | G | C | C | A | C | C | A | G | C | C | A | A | T | T | G | A | T | G | A | A | C | T | G | T | T | A | A | T | T | - | - | - | C | A | T | G | A | A | G | T | C | - | - | C | A | T | A | C | A | T | - | - | - | - | - | - | - | - | T | G | C | A | A | A | A | C | T | C | A | A | A | A | A | T | G | C | C | A | T | A | G | C | C | C | A | A | A | C | A | G | T | A | C | A | A | T | - | - | - | - | - | - | - | - | - | A | A | T | A | T | - | - | - | - | G | T | A | A | T | T | C | - | - | - | - | - | - | - | - | - | - | - | - | - | - | - | - | - | - | - | A | T | A | T | T | T | G | T | G | T | T | G | A | C | A | C | C | T | A | A | A | A | A | T | A | A | A | T | C | A | G | T | T | G | T | T | T | - | - | - | - | - | - | - | - | - | - |
|  | MYZPE13164\_G006\_V2.0\_0001 | A | T | C | A | C | G | C | C | A | C | - | - | - | - | - | - | - | - | - | - | - | - | - | - | - | C | C | A | G | C | C | - | - | - | - | - | G | T | C | C | C | A | C | T | T | T | G | A | A | T | C | - | - | - | - | - | - | - | - | - | G | G | T | C | G | G | T | G | G | C | A | G | T | G | T | A | T | A | T | A | A | A | G | G | T | G | T | C | C | G | A | G | A | T | C | T | C | G | A | T | T | C | C | G | G | C | A | T | A | A | C | A | C | A | T | C | C | T | T | T | T | G | T | G | C | A | G | A | T | A | T | A | T | T | C | T | T | T | G | T | T | T | T | - | - | - | - | - | - | C | A | C | - | T | T | T | A | A | C | C | - | - | C | T | C | A | A | A | A | A | C | C | A | - | - | - | A | C | C | A | A | - | A | A | A | C | C | A | A | A | A | A | T | G | G | G | C | G | C | T | G | A | A | A | A | A | G | T | A | T | C | C | A | T | G | A | A | C | A | T | C | G | T | C | G | T | C | G | T | C | G | G | A | C | A | A | G | T | C | C | A | A | T | C | T | G | C | C | A | A | G | G | C | C | A | T | C | A | A | A | T | C | G | G | T | C | G | C | C | A | A | A | G | C | C | A | C | T | C | C | A | G | C | C | A | C | C | A | G | C | C | A | A | T | T | G | A | T | G | A | A | C | T | G | T | T | A | A | T | T | - | - | - | C | A | T | G | T | A | G | T | C | - | - | A | A | T | A | C | A | T | - | - | - | - | - | - | - | - | T | G | C | A | A | A | A | C | T | C | A | A | A | A | A | T | G | C | C | A | T | A | T | C | C | C | A | A | A | C | A | G | T | A | C | A | A | T | - | - | - | - | - | - | - | - | - | A | A | T | A | T | - | - | - | - | G | T | A | A | C | T | C | - | - | - | - | - | - | - | - | - | - | - | - | - | - | - | - | - | - | - | A | T | A | T | C | T | G | T | A | T | T | G | A | C | A | C | C | G | C | A | A | A | A | T | A | A | A | T | C | A | G | T | C | G | T | T | T | - | - | - | - | - | - | - | - | - | - |
|  | MYZPE13164\_G006\_V2.0\_0001 | A | T | C | A | C | G | C | C | A | T | G | - | - | - | - | - | - | - | - | - | - | - | - | - | - | G | C | C | G | G | C | - | - | - | - | - | G | C | C | C | C | A | C | T | T | T | T | A | T | A | C | - | - | - | - | - | - | - | - | - | G | A | T | C | G | G | C | G | G | G | A | G | T | G | T | A | T | A | T | A | A | A | G | G | T | G | T | C | T | G | A | G | A | T | C | T | A | G | A | A | T | C | C | G | G | C | A | T | A | A | C | A | C | A | T | C | C | T | T | T | T | G | T | G | C | A | G | A | T | C | T | A | T | T | C | T | T | T | G | T | T | T | T | - | - | - | - | - | - | C | A | C | - | T | T | T | A | A | T | C | - | - | C | T | C | A | A | A | A | C | C | A | A | - | C | C | A | C | C | A | A | - | A | A | A | C | C | A | A | A | A | A | T | G | G | G | C | G | C | A | G | A | A | A | A | A | G | T | A | T | C | C | A | T | G | A | A | C | A | T | C | G | T | C | G | T | T | G | T | C | G | G | A | C | A | A | G | T | C | C | A | A | T | C | T | G | C | C | A | A | G | G | C | C | A | T | C | A | A | A | T | C | G | G | T | C | G | C | C | A | A | A | G | C | C | A | C | C | C | C | A | G | C | C | A | C | C | A | G | C | C | A | A | T | T | G | A | T | G | A | A | C | T | G | T | T | A | A | T | T | - | - | - | - | A | T | G | T | A | G | T | C | - | - | G | A | T | A | C | A | T | - | - | - | - | - | - | - | - | T | G | C | A | A | A | A | C | T | C | A | A | A | A | C | T | G | C | C | A | T | A | G | C | C | C | A | A | A | C | A | G | T | A | C | A | A | T | - | - | - | - | - | - | - | - | - | A | A | T | A | T | - | - | - | - | G | T | A | A | T | T | C | - | - | - | - | - | - | - | - | - | - | - | - | - | - | - | - | - | - | - | A | T | A | T | C | T | G | T | A | T | T | G | A | C | A | C | T | G | C | A | A | A | A | T | A | A | A | T | C | A | G | A | C | A | C | T | T | - | - | - | - | - | - | - | - | - | - |
|  | MYZPE13164\_G006\_V2.0\_0001 | A | T | C | A | C | G | C | C | A | C | G | C | T | G | C | C | - | - | - | - | - | - | - | - | - | G | C | C | G | C | C | - | - | - | - | - | G | C | C | C | C | A | C | T | T | A | A | C | A | C | T | - | - | - | - | - | - | - | - | - | T | G | T | T | G | A | T | G | G | A | A | G | T | G | T | A | T | A | T | A | A | A | G | G | C | G | T | C | C | G | A | G | A | T | C | T | T | G | A | T | T | C | C | G | G | C | A | T | A | A | C | A | C | A | T | C | T | T | T | T | T | G | T | - | - | - | G | A | T | C | T | A | T | T | C | T | T | T | G | T | T | T | T | - | - | - | - | - | - | C | A | C | - | T | T | T | A | A | T | C | - | - | C | T | C | A | A | A | A | A | C | C | A | - | - | - | A | C | C | A | A | - | A | A | A | C | C | A | A | A | A | A | T | G | G | G | C | G | C | T | G | A | A | A | A | A | G | T | A | A | C | C | A | T | G | A | A | C | A | T | C | G | T | T | G | T | C | G | T | T | G | G | A | C | A | A | G | T | C | C | A | A | T | C | T | G | C | C | A | A | G | G | C | C | A | T | C | A | A | A | T | C | G | G | T | C | G | C | C | A | A | A | G | C | C | A | C | C | C | C | A | G | C | C | A | C | C | A | G | C | C | A | A | T | T | G | A | T | G | A | A | C | T | G | C | T | A | A | T | T | - | - | - | T | A | T | G | T | A | G | T | C | - | - | C | A | T | A | C | A | T | - | - | - | - | - | - | - | - | T | C | C | A | A | A | A | C | T | C | A | A | A | A | A | T | G | C | C | A | T | A | G | T | C | C | A | A | A | C | A | A | T | A | C | A | A | T | - | - | - | - | - | - | - | - | - | A | A | T | A | T | - | - | - | - | G | T | A | A | C | T | C | A | T | - | - | - | - | - | - | - | - | - | - | - | - | - | - | - | - | A | A | T | A | T | C | A | G | T | A | T | T | G | C | C | A | T | C | G | C | A | A | A | A | T | A | A | A | T | C | A | G | T | C | A | T | T | T | - | - | - | - | - | - | - | - | - | - |
|  | MYZPE13164\_G006\_V2.0\_0001 | A | T | C | A | C | G | C | C | A | T | C | G | C | C | C | C | C | C | C | C | C | C | C | T | C | C | C | C | A | C | T | - | - | - | - | - | T | C | C | C | C | A | C | T | T | - | - | A | T | T | C | - | - | - | - | - | - | - | - | - | A | G | T | C | A | G | C | G | G | G | A | G | T | G | T | A | T | A | T | A | A | A | G | G | C | G | T | G | C | G | T | G | A | T | C | T | C | G | G | T | T | A | A | A | G | C | A | T | A | A | C | A | C | A | T | - | T | T | T | T | T | G | T | G | C | A | G | A | T | C | T | A | T | T | C | T | T | T | G | T | T | T | T | - | - | - | - | - | - | C | A | C | - | T | T | T | A | A | T | C | - | - | C | T | C | A | A | A | A | A | A | C | A | - | - | - | A | C | C | A | A | - | A | A | A | C | C | A | A | A | A | A | T | G | G | G | C | G | C | T | G | A | A | A | A | A | G | T | A | A | C | C | A | T | G | A | A | C | A | T | C | G | T | C | G | T | C | G | T | A | G | G | A | C | A | A | G | T | C | C | A | A | T | C | T | G | C | C | A | A | G | G | C | C | A | T | C | A | A | G | A | C | C | G | T | C | G | C | C | A | A | G | G | C | C | T | C | C | C | C | A | G | C | C | A | C | C | A | A | C | C | A | A | T | T | G | A | T | G | A | A | C | T | G | T | T | A | A | A | T | - | - | - | C | A | T | G | T | A | A | T | C | - | - | C | A | T | A | C | A | T | - | - | - | - | - | - | - | - | T | G | C | A | A | A | A | C | C | C | A | A | A | A | A | T | G | C | C | A | T | A | G | C | C | C | A | A | A | C | A | G | T | A | T | A | A | T | - | - | - | - | - | - | - | - | - | A | A | T | A | T | - | - | - | - | G | T | A | A | T | T | C | - | - | - | - | - | - | - | - | - | - | - | - | - | - | - | - | - | - | - | A | T | A | T | - | T | A | T | A | T | T | G | A | C | A | C | C | G | C | A | A | A | A | T | A | A | A | T | C | A | G | T | A | A | T | T | T | - | - | - | - | - | - | - | - | - | - |
|  | MYZPE13164\_G006\_V2.0\_0000 | A | T | C | A | C | G | C | C | A | T | - | - | - | - | - | - | - | - | - | - | - | - | - | - | - | - | - | - | - | - | - | - | - | - | - | - | C | A | T | C | C | C | C | A | C | T | T | A | T | T | C | - | - | - | - | - | - | - | - | - | A | T | T | C | A | G | C | G | G | G | A | G | T | G | T | A | T | A | T | A | A | A | G | T | C | G | T | C | C | G | A | G | A | T | C | T | C | A | A | T | T | C | T | G | G | C | A | T | A | A | C | A | C | A | T | T | T | T | T | T | T | G | T | A | C | A | G | A | T | C | T | A | T | T | C | A | T | T | G | T | T | T | T | - | - | - | - | - | - | C | A | C | - | T | T | T | A | A | A | C | - | - | C | T | C | A | A | A | A | A | C | C | C | - | - | - | A | C | C | A | A | - | A | A | A | C | C | - | A | A | A | A | T | G | G | G | C | G | C | A | G | A | A | A | A | G | G | T | A | T | C | C | A | T | G | A | A | C | A | T | C | G | T | T | G | T | C | G | T | C | G | G | A | C | A | A | G | T | C | A | A | A | T | C | T | G | C | C | A | A | G | G | C | C | G | T | C | A | A | G | A | C | C | G | T | C | T | C | C | A | A | G | G | C | C | A | C | A | C | C | A | G | C | C | A | C | C | A | A | C | C | A | A | C | T | G | A | T | G | A | A | C | T | G | T | T | A | A | A | T | - | - | - | C | A | T | G | T | A | G | T | C | T | A | C | A | T | A | C | A | T | - | - | - | - | - | - | - | - | T | G | G | A | A | A | A | C | T | C | A | A | A | A | A | T | G | C | C | A | T | A | G | C | C | C | A | A | A | T | A | G | T | A | C | A | A | T | - | - | - | - | - | - | - | - | - | A | A | T | A | T | - | - | - | - | G | T | A | A | T | T | C | - | - | - | - | - | - | - | - | - | - | - | - | - | - | - | - | - | - | - | A | T | T | T | C | A | A | T | A | T | T | T | A | C | A | A | C | A | C | A | A | A | A | T | A | A | A | T | C | A | A | T | A | A | T | T | T | - | - | - | - | - | - | - | - | - | - |
|  | MYZPE13164\_G006\_V2.0\_0001 | A | T | C | A | C | G | C | C | A | T | - | - | - | - | - | - | - | - | - | - | - | - | - | - | C | C | C | C | A | C | C | - | - | - | - | - | T | T | C | C | C | A | C | T | T | T | C | A | G | C | C | - | - | - | - | - | - | - | - | - | T | G | C | C | G | A | C | G | A | G | A | G | C | A | T | A | T | A | T | - | A | A | G | G | C | G | T | G | C | G | T | G | A | T | T | T | C | G | A | T | T | A | A | G | G | C | A | T | A | A | C | A | C | A | T | C | A | T | T | T | T | G | T | G | C | A | G | A | T | A | A | A | T | T | C | T | T | T | G | T | T | T | T | - | - | - | - | - | - | C | A | C | - | T | T | T | A | A | T | C | - | - | C | A | C | A | A | A | A | C | C | A | A | - | - | - | C | C | A | A | C | - | A | A | A | T | C | A | A | A | A | A | T | G | G | G | C | G | C | A | G | A | A | A | A | G | G | T | A | T | C | C | A | T | G | A | A | C | A | T | C | G | T | C | G | T | C | G | T | C | G | G | A | C | A | A | G | T | C | C | A | G | T | C | T | G | C | C | A | A | G | G | C | C | A | T | C | A | A | A | T | C | G | G | T | C | G | C | C | A | A | A | G | C | C | A | C | C | C | C | A | G | C | C | A | C | C | A | C | C | C | A | A | T | T | G | A | T | G | A | A | C | T | G | T | T | A | A | T | T | - | - | - | T | A | T | G | T | A | G | T | T | - | - | C | A | T | C | C | G | T | - | - | - | - | - | C | C | A | T | C | G | C | A | A | A | C | T | C | A | A | A | C | A | T | G | T | C | A | T | A | G | C | C | C | A | A | A | T | A | A | T | A | C | A | A | T | - | - | - | - | - | - | - | - | - | - | - | - | A | T | - | - | - | - | A | T | T | A | T | T | C | - | - | - | - | - | - | - | - | - | - | - | - | - | - | - | - | - | - | - | A | A | A | T | T | G | A | T | A | T | T | G | A | C | A | A | C | A | T | G | A | A | A | T | A | A | A | A | C | A | G | C | C | T | C | T | T | T | - | - | - | - | - | - | - | - | - |
|  | MYZPE13164\_G006\_V2.0\_0001 | A | T | C | A | C | G | C | C | A | C | - | - | - | - | - | - | - | - | - | - | - | - | - | - | - | A | C | C | A | C | C | - | - | - | - | - | T | T | T | C | C | A | T | T | T | T | C | A | G | C | C | - | - | - | - | - | - | - | - | - | G | C | C | G | G | G | C | G | - | - | A | G | T | G | T | A | T | A | T | A | A | A | G | G | C | G | T | G | C | G | T | G | A | T | C | T | T | G | A | T | T | A | A | G | G | C | A | T | A | A | C | A | C | A | T | C | T | T | T | T | T | G | T | G | C | A | G | A | T | A | T | A | T | T | C | A | T | T | G | T | T | T | T | - | - | - | - | - | - | C | A | C | - | T | T | T | A | A | T | C | - | - | C | T | C | A | A | A | A | A | C | C | A | - | - | - | A | C | C | A | A | - | A | A | A | C | C | A | A | A | A | A | T | G | G | G | C | G | C | T | G | A | A | A | A | A | G | T | A | T | C | C | A | T | G | A | A | C | A | T | C | G | T | T | G | T | C | G | T | C | G | G | A | C | A | A | G | T | C | C | A | A | T | C | T | G | C | C | A | A | G | G | C | C | A | T | C | A | A | A | T | C | G | G | T | C | G | C | C | A | A | A | G | C | C | A | C | C | C | C | A | G | C | C | A | C | C | A | G | C | C | A | A | T | T | G | A | T | G | A | A | C | T | G | T | T | A | A | T | T | - | - | - | C | A | T | G | T | A | A | T | T | - | - | C | A | T | C | C | A | T | G | A | G | C | T | C | C | A | T | C | T | C | A | A | A | C | T | C | A | A | A | C | A | T | G | C | C | A | T | A | G | C | C | C | A | A | A | T | A | A | T | T | C | A | A | T | - | - | - | - | - | - | - | - | - | A | A | - | - | - | - | - | - | - | A | T | G | A | T | T | C | - | - | - | - | - | - | - | - | - | - | - | - | - | - | - | - | - | - | - | A | A | A | T | T | G | A | T | A | T | G | T | A | A | A | C | C | A | T | G | A | A | A | T | A | A | A | A | C | A | G | C | C | T | C | T | T | C | C | G | G | A | A | T | T | - | - |
|  | MYZPE13164\_G006\_V2.0\_0001 | A | T | C | A | C | G | C | C | A | C | - | - | - | - | - | - | - | - | - | - | - | - | - | - | - | G | C | C | A | C | C | - | - | - | - | - | T | T | C | T | C | A | C | T | T | T | C | A | G | C | C | - | - | - | - | - | - | - | - | - | G | C | C | T | G | A | C | G | - | - | A | G | T | G | T | A | T | A | T | A | A | A | G | G | C | A | T | G | C | T | T | G | A | T | C | T | T | G | A | T | T | A | A | A | A | C | A | T | A | A | C | A | C | A | T | C | T | T | T | T | T | G | T | G | C | A | G | A | T | A | A | A | T | T | C | T | T | T | G | T | T | T | T | - | - | - | - | - | - | C | A | C | - | T | T | T | A | A | C | C | - | - | C | T | C | A | A | A | A | A | C | C | A | - | - | - | A | C | C | A | A | - | A | A | A | C | C | A | A | A | A | A | T | G | G | G | C | G | C | A | G | A | A | A | A | T | G | T | A | T | C | C | A | T | G | A | A | C | A | T | C | G | T | T | G | T | C | G | T | C | G | G | A | C | A | A | G | T | C | C | A | A | T | C | T | G | C | C | A | A | G | G | C | C | A | T | C | A | A | A | T | C | T | G | T | C | G | C | C | A | A | A | G | C | C | A | C | C | C | C | A | G | C | C | A | C | C | A | G | C | C | A | A | T | T | G | A | T | G | A | A | C | T | G | T | T | A | A | T | T | - | - | - | C | A | T | G | T | A | G | T | T | - | - | C | A | T | C | C | A | T | - | - | - | - | - | C | C | A | T | C | G | C | A | A | A | C | T | C | A | A | A | C | A | T | G | C | C | A | T | A | G | C | C | A | A | A | A | T | A | A | T | A | C | A | A | T | - | - | - | - | - | - | A | C | G | A | G | T | A | T | - | - | - | - | A | T | T | A | T | T | T | - | - | - | - | - | - | - | - | - | - | - | - | - | - | - | - | - | - | - | A | G | A | T | T | G | A | T | A | T | T | G | A | C | A | C | C | A | T | G | A | A | A | T | A | A | A | A | C | A | G | C | T | T | C | T | T | - | - | - | - | - | - | - | - | - | - |
|  | MYZPE13164\_G006\_V2.0\_0001 | A | T | C | A | C | G | C | C | A | T | C | G | C | C | C | - | - | - | - | - | - | - | - | - | - | C | T | C | T | C | C | - | - | - | - | - | C | C | G | C | C | A | C | T | T | C | A | A | G | C | T | G | - | - | - | - | - | - | C | A | G | G | C | C | A | T | C | A | G | T | A | G | T | G | T | A | T | A | T | A | A | A | G | G | A | G | T | C | C | G | A | G | A | T | C | T | C | G | A | T | T | C | C | G | G | C | A | T | A | A | C | A | C | A | T | - | T | T | T | T | T | G | T | G | C | A | A | A | T | A | T | A | T | T | C | T | T | T | G | T | T | T | T | - | - | - | - | - | - | C | T | C | T | T | T | T | A | A | A | C | - | - | C | T | A | A | A | A | A | A | C | C | A | - | - | - | A | C | C | A | A | C | A | A | A | T | C | A | A | A | A | A | T | G | G | G | C | G | C | T | G | A | A | A | A | G | G | T | A | T | C | C | A | T | G | A | A | C | A | T | C | G | T | T | G | T | C | G | T | C | G | G | A | C | A | A | G | T | C | C | A | A | T | C | T | G | C | C | A | A | G | G | C | C | A | T | C | A | A | G | A | C | C | G | T | C | G | C | C | A | A | G | G | C | C | T | C | C | C | C | A | G | C | C | A | C | C | A | A | C | C | A | A | T | T | G | A | T | G | A | A | C | T | G | T | T | A | A | T | T | - | - | - | C | A | T | G | T | A | G | T | C | - | - | A | A | T | C | C | A | - | - | - | - | - | - | - | - | - | T | T | T | C | A | A | A | C | T | C | A | A | A | A | A | T | G | C | C | A | T | A | G | T | C | C | A | A | A | C | A | A | T | A | C | A | A | T | - | - | - | - | - | - | - | - | - | A | A | T | A | T | - | - | - | - | G | T | A | A | C | T | C | A | T | - | - | - | - | - | - | - | - | - | - | - | - | - | - | - | - | A | A | T | A | T | C | A | G | T | A | T | T | G | C | C | A | C | C | G | C | A | A | A | A | T | A | A | A | T | C | A | G | T | C | A | C | A | T | - | - | - | - | - | - | - | - | - | - |
|  | MYZPE13164\_G006\_V2.0\_0001 | A | T | C | A | C | G | C | C | A | C | - | - | - | - | - | - | - | - | - | - | - | - | - | - | - | T | T | T | G | C | T | - | - | - | - | - | T | C | C | C | C | A | C | T | T | G | C | A | A | G | C | G | G | C | C | - | - | G | A | C | G | A | C | G | A | G | T | G | C | G | T | A | G | G | T | A | T | A | T | A | A | A | G | G | T | G | T | A | C | G | T | G | T | T | C | A | C | G | A | T | T | C | C | G | G | C | A | T | A | A | C | A | C | A | T | A | T | T | T | T | T | G | T | G | C | A | G | A | T | A | T | A | T | T | C | T | T | T | G | T | T | T | T | - | - | - | - | - | - | C | A | C | - | - | T | C | A | A | C | A | - | - | C | T | C | A | A | A | A | A | C | C | A | - | - | - | A | C | C | A | A | C | A | A | A | T | C | A | A | A | A | A | T | G | G | G | C | G | C | A | G | A | A | A | A | G | G | T | A | T | C | C | A | T | G | A | A | C | A | T | C | G | T | T | G | T | C | G | T | C | G | G | A | C | A | A | G | T | C | C | A | A | T | C | T | G | C | C | A | A | A | G | C | A | G | T | C | A | A | G | A | C | C | G | T | C | G | C | C | A | A | G | G | C | C | G | C | C | C | C | A | G | C | C | A | C | C | A | G | C | C | A | A | T | T | G | A | T | G | A | A | C | T | G | T | T | A | A | T | T | - | - | - | C | A | C | G | T | A | G | T | T | - | - | C | A | T | C | C | A | - | - | - | - | - | - | - | - | - | T | C | A | C | A | A | A | C | T | C | A | A | A | A | A | T | G | C | C | A | T | A | G | A | C | C | A | A | A | C | A | A | T | A | C | A | A | T | - | - | - | - | - | - | - | - | - | A | T | T | A | T | - | - | - | - | G | T | A | A | A | T | C | T | T | A | T | - | - | - | - | - | - | - | - | - | - | - | - | - | - | - | - | - | - | - | - | T | G | A | T | T | T | G | A | C | G | C | C | A | T | G | A | A | A | T | A | A | A | A | C | A | G | C | C | A | C | A | T | C | C | G | A | T | A | T | T | C | A |
|  | MYZPE13164\_G006\_V2.0\_0000 | A | T | C | A | C | G | C | C | A | C | - | - | - | - | - | - | - | - | - | - | - | - | - | - | - | C | A | C | G | C | C | - | - | - | - | - | G | C | T | C | G | A | C | A | T | T | G | A | G | C | - | - | - | - | - | - | - | - | - | - | G | A | A | A | A | T | C | G | T | C | A | G | T | G | T | A | T | A | A | A | A | T | A | A | C | G | T | A | C | T | G | G | T | G | T | A | T | C | G | T | T | C | C | G | G | C | A | T | A | A | C | A | C | A | A | - | T | T | T | T | C | G | T | G | C | A | A | A | T | C | T | A | A | T | A | A | T | T | A | T | T | T | T | - | - | - | - | - | - | C | A | C | - | T | T | T | A | C | T | C | - | - | C | T | T | A | A | A | C | A | C | C | A | - | - | - | A | C | C | A | A | - | A | A | A | C | C | A | A | A | A | A | T | G | G | G | C | G | C | T | G | A | A | A | A | A | G | T | A | A | C | C | A | T | G | A | A | C | A | T | C | G | T | C | G | T | C | G | T | A | G | G | A | C | A | A | G | T | C | C | A | A | T | C | T | G | C | T | A | A | G | G | C | C | A | T | C | A | A | A | T | C | G | G | T | C | G | C | C | A | A | A | G | C | C | A | C | T | C | C | A | G | C | A | A | C | C | A | G | C | C | A | A | T | T | A | A | T | G | A | A | C | T | G | T | T | A | A | A | T | - | - | - | C | A | T | G | A | A | G | T | T | C | A | C | A | T | C | A | A | - | - | - | - | - | - | - | - | - | C | C | T | C | T | A | A | C | T | C | A | A | A | A | A | T | G | C | A | A | T | A | G | C | C | C | A | A | A | C | A | G | T | T | C | A | T | A | - | - | - | - | - | - | - | - | - | A | T | T | T | T | - | - | - | - | A | T | A | A | T | T | T | T | T | T | C | C | - | - | - | - | - | T | T | A | T | T | A | T | A | A | T | A | A | T | G | T | G | T | A | T | T | G | A | C | A | T | T | T | C | A | A | A | A | T | A | A | A | T | C | A | G | T | C | G | C | T | T | - | - | - | - | - | - | - | - | - | - |
|  | MYZPE13164\_G006\_V2.0\_0001 | A | T | C | A | C | G | C | C | A | C | - | - | - | - | - | - | - | - | - | - | - | - | - | - | - | C | A | C | G | C | C | - | - | - | - | - | G | C | T | C | C | A | C | A | T | T | T | A | T | C | - | - | - | - | - | - | - | - | - | - | G | A | A | A | A | T | C | A | C | C | G | G | T | G | T | A | T | A | A | A | A | T | A | A | C | G | T | A | C | C | A | G | T | G | C | T | T | A | G | T | T | T | C | G | G | C | A | T | A | A | C | G | C | A | A | - | T | T | T | T | C | G | T | G | C | A | A | A | T | C | T | A | A | T | C | T | T | C | A | T | T | T | T | - | - | - | - | - | - | C | A | C | - | T | T | T | A | C | T | C | - | - | C | T | C | A | A | A | A | A | C | C | A | A | - | - | A | C | C | A | A | - | A | A | A | C | C | A | A | A | A | A | T | G | G | G | C | G | C | T | G | A | A | A | A | A | G | T | A | A | C | C | A | T | G | A | A | C | A | T | C | G | T | C | G | T | C | G | T | C | G | G | A | C | A | A | G | T | C | C | A | A | T | C | T | G | C | C | A | A | G | G | C | C | A | T | C | A | A | A | T | C | G | G | T | C | G | C | C | A | A | A | G | C | C | A | C | C | C | C | A | G | C | C | A | C | C | A | G | C | C | A | A | T | T | G | A | T | G | A | A | C | T | G | T | T | A | A | T | T | - | - | - | C | A | T | G | A | A | A | T | C | C | A | C | A | T | C | A | A | - | - | - | - | - | - | - | - | - | T | C | T | C | A | A | A | C | T | C | A | A | A | A | A | T | G | C | C | A | T | A | G | C | C | C | G | A | A | C | A | G | T | A | C | A | T | A | - | - | - | - | - | - | - | - | - | A | T | T | T | T | - | - | - | - | A | T | A | A | T | T | T | G | T | T | T | C | - | - | - | - | - | T | A | A | T | T | A | T | A | A | T | A | A | T | G | C | G | T | A | T | T | G | A | C | A | T | T | G | C | A | A | A | A | T | A | A | A | T | C | A | G | T | C | G | T | T | T | - | - | - | - | - | - | - | - | - | - |
|  | MYZPE13164\_G006\_V2.0\_0000 | A | T | C | A | C | G | C | C | A | C | - | - | - | - | - | - | - | - | - | - | - | - | - | - | - | A | A | C | G | C | T | - | - | - | - | - | - | - | - | - | - | - | - | - | - | - | - | A | G | C | - | - | - | - | - | - | - | - | - | - | G | A | A | A | G | C | C | G | C | C | A | C | T | G | T | A | T | A | A | A | A | G | G | A | C | G | T | A | C | C | G | A | T | G | C | T | T | C | A | T | T | C | C | G | A | C | A | T | A | A | C | G | C | A | A | - | T | T | T | T | T | G | T | G | C | A | A | A | T | A | T | A | A | T | C | T | T | C | A | A | T | T | T | - | - | - | - | - | - | C | A | C | - | T | C | T | A | A | T | C | - | - | C | T | C | A | A | A | A | A | C | A | A | A | - | - | C | C | A | A | A | - | A | A | A | C | C | A | A | A | A | A | T | G | G | G | C | G | C | T | G | A | A | A | A | A | G | T | A | A | C | C | A | T | G | A | A | C | A | T | C | G | T | T | G | T | C | G | T | C | G | G | A | C | A | A | G | T | C | C | A | A | T | C | T | G | C | C | A | A | G | G | C | C | A | T | C | A | A | A | T | C | G | G | T | C | G | C | C | A | A | A | G | C | C | A | C | C | C | C | A | G | C | C | A | C | C | A | G | C | C | A | A | T | T | G | A | T | G | A | A | C | T | G | T | T | A | A | T | T | - | - | - | C | A | G | G | A | A | G | T | C | C | A | C | A | T | C | A | A | - | - | - | - | - | - | - | - | - | C | C | T | C | A | A | A | C | T | C | A | A | A | A | A | T | G | C | C | A | T | A | G | C | C | C | A | A | A | C | A | G | T | A | C | A | T | A | - | - | - | - | - | - | - | - | - | A | T | T | T | T | - | - | - | - | A | T | A | A | T | T | T | G | T | T | T | A | - | - | - | - | - | T | A | A | T | T | T | T | A | A | A | A | A | T | G | T | G | T | A | T | T | G | A | T | A | T | T | A | C | A | A | A | A | T | A | A | A | C | C | A | G | T | C | A | C | T | T | - | - | - | - | - | - | - | - | - | - |
|  | MYZPE13164\_G006\_V2.0\_0001 | A | T | C | A | C | G | C | C | A | C | - | - | - | - | - | - | - | - | - | - | - | - | - | - | - | A | A | C | G | C | C | - | - | - | - | - | - | - | - | - | - | - | - | A | T | T | A | A | G | C | - | - | - | - | - | - | - | - | - | - | G | A | A | A | G | C | C | G | C | C | G | C | T | G | T | A | T | A | A | A | A | G | G | A | C | G | T | G | T | T | G | A | T | G | C | T | T | C | A | T | T | C | C | G | G | C | A | T | A | A | C | G | C | A | A | - | T | T | T | T | T | G | T | G | C | A | A | T | T | A | C | A | A | T | C | T | T | C | A | T | T | T | C | - | - | - | - | - | - | C | A | C | - | T | T | T | A | A | T | C | - | - | C | T | C | A | A | A | A | A | C | C | A | - | - | - | A | C | C | A | A | - | A | A | A | C | C | A | A | A | A | A | T | G | G | G | C | G | C | T | G | A | A | A | A | A | G | T | A | A | C | C | A | T | G | A | A | C | A | T | C | G | T | T | G | T | C | G | T | C | G | G | A | C | A | A | G | T | C | C | A | A | T | C | T | G | C | C | A | A | G | G | C | C | A | T | C | A | A | A | T | C | G | G | T | C | G | C | C | A | A | A | G | C | C | A | C | C | C | C | A | G | C | C | A | C | C | A | G | C | C | A | A | T | T | G | A | T | G | A | A | C | T | G | T | T | A | A | T | T | - | - | - | C | A | T | G | A | A | T | T | C | C | A | C | A | T | C | A | A | - | - | - | - | - | - | - | - | - | T | C | T | C | A | A | A | C | T | C | A | A | A | A | A | T | G | C | C | A | T | A | G | C | C | C | A | A | A | C | A | G | T | A | C | A | T | A | - | - | - | - | - | - | - | - | - | A | T | T | T | T | - | - | - | - | A | T | A | A | T | T | T | G | T | T | T | C | A | A | - | A | T | T | T | A | T | T | A | T | A | A | A | A | A | T | G | T | G | T | T | C | T | G | C | C | A | T | T | A | C | A | A | A | A | T | A | A | A | T | C | A | G | T | C | A | T | T | T | - | - | - | - | - | - | - | - | - | - |
|  | MYZPE13164\_G006\_V2.0\_0001 | A | T | C | A | C | G | C | C | A | C | - | - | - | - | - | - | - | - | - | - | - | - | - | - | - | C | A | C | T | C | C | T | C | C | G | T | G | C | T | C | T | A | C | G | T | C | A | G | G | C | - | - | - | - | - | - | - | - | - | - | G | A | A | A | G | C | C | A | C | C | T | G | T | G | T | A | T | A | A | A | A | T | G | A | C | G | T | A | C | T | G | A | T | G | C | T | T | C | A | G | T | C | T | A | G | C | A | T | A | A | C | G | C | A | A | - | T | T | T | T | T | G | T | G | C | A | A | A | C | A | T | A | A | T | C | T | T | T | A | A | T | T | T | - | - | - | - | - | - | C | A | C | - | T | T | T | A | A | T | C | - | - | C | T | C | A | A | A | A | A | C | C | A | A | - | - | A | C | A | A | A | - | A | A | A | C | C | A | A | A | A | A | T | G | G | G | C | G | C | T | G | A | A | A | A | A | G | T | A | A | C | C | A | T | G | A | A | C | A | T | C | G | T | T | G | T | C | G | T | C | G | G | A | C | A | A | G | T | C | C | A | A | T | C | T | G | C | C | A | A | G | G | C | C | A | T | C | A | A | A | T | C | G | G | T | C | G | C | C | A | A | A | G | C | C | A | C | C | C | C | A | G | C | C | A | C | C | A | G | C | C | A | A | T | T | G | A | T | G | A | A | C | T | G | T | T | A | A | T | T | - | - | - | C | A | A | G | A | - | - | - | C | C | A | C | A | T | C | A | A | - | - | - | - | - | - | - | - | - | A | C | T | C | A | A | A | C | T | C | A | A | A | A | A | T | G | C | C | A | T | A | G | C | C | C | A | A | A | C | A | G | T | A | C | T | T | A | - | - | - | - | - | - | - | - | - | A | T | T | T | T | - | - | - | - | A | T | A | A | T | T | T | A | T | T | T | C | T | - | - | A | A | T | T | A | T | A | A | T | A | A | T | T | A | C | A | T | G | T | A | T | T | G | A | C | A | T | T | G | C | A | A | A | A | T | A | A | A | T | C | A | G | T | A | A | C | T | T | - | - | - | - | - | - | - | - | - | - |
|  | MYZPE13164\_G006\_V2.0\_0001 | A | T | C | A | C | G | C | C | A | C | - | - | - | - | - | - | - | - | - | - | - | - | - | - | - | C | A | C | G | C | T | - | - | - | - | - | G | C | T | C | T | A | T | A | T | C | A | G | G | T | - | - | - | - | - | - | - | - | - | - | G | A | A | A | A | T | C | G | - | - | G | C | T | G | T | A | T | A | A | A | A | A | G | A | C | G | C | A | C | C | G | A | T | G | C | T | A | C | A | T | T | C | C | A | G | C | A | T | A | A | C | G | C | A | A | - | T | T | T | T | T | G | T | G | C | A | A | T | T | A | C | A | A | T | C | T | T | C | A | T | T | T | T | - | - | - | - | - | - | C | A | C | - | T | C | T | A | A | C | C | - | - | C | T | C | A | A | A | A | A | A | C | C | A | - | - | A | C | C | A | A | - | A | A | A | C | C | A | A | A | A | A | T | G | G | G | C | G | C | T | G | A | A | A | A | A | G | T | A | A | C | C | A | T | G | A | A | C | A | T | C | G | T | C | G | T | C | G | T | C | G | G | A | C | A | A | G | T | C | C | A | A | T | C | T | G | C | C | A | A | G | G | C | C | A | T | C | A | A | A | T | C | C | G | T | C | G | C | C | A | A | G | G | C | C | G | C | C | C | A | A | G | C | C | A | C | C | A | G | C | C | A | A | T | T | G | A | T | G | A | A | C | T | G | T | T | A | A | T | T | - | - | - | C | A | T | G | A | A | G | T | C | C | A | C | A | T | C | A | A | - | - | - | - | - | - | - | - | - | T | C | T | C | A | A | G | C | T | C | A | A | A | A | A | T | G | C | C | A | T | A | G | C | C | C | A | A | A | C | A | G | T | A | C | A | T | T | - | - | - | - | - | - | - | - | - | A | T | T | T | T | - | - | - | - | A | T | A | A | T | T | T | T | T | T | T | T | A | - | - | - | A | T | T | A | T | A | A | T | A | A | T | T | A | C | G | T | G | T | A | T | T | G | A | C | A | T | T | G | C | A | A | A | A | T | A | A | A | T | C | A | G | T | C | A | C | T | T | - | - | - | - | - | - | - | - | - | - |
|  | MYZPE13164\_G006\_V2.0\_0001 | A | T | C | A | C | G | C | C | A | C | - | - | - | - | - | - | - | - | - | - | - | - | - | - | - | C | A | C | G | C | T | - | - | - | - | - | G | C | T | A | A | G | A | G | T | T | T | A | T | C | - | - | - | - | - | - | - | - | - | - | G | A | A | A | A | T | T | C | A | C | A | G | T | G | T | A | T | A | A | A | A | G | G | A | C | A | - | A | C | A | G | G | T | G | C | T | T | C | G | T | T | C | T | G | G | C | A | T | A | A | C | A | C | G | A | - | T | T | T | T | T | G | T | G | C | A | A | A | T | C | T | T | A | T | C | T | T | C | G | A | T | T | T | - | - | - | - | - | - | C | A | T | - | T | C | T | A | A | T | C | - | - | C | T | C | A | A | A | A | A | C | C | A | A | - | - | A | C | C | A | A | - | A | A | A | C | C | A | A | A | A | A | T | G | G | G | C | G | C | T | G | A | A | A | A | A | G | T | A | A | C | C | A | T | G | A | A | C | A | T | C | G | T | T | G | T | C | G | T | C | G | G | A | C | A | A | G | T | C | C | A | A | T | C | T | G | C | C | A | A | G | G | C | C | A | T | C | A | A | A | T | C | G | G | T | C | G | C | C | A | A | A | G | C | C | G | C | C | C | C | A | G | C | C | A | C | C | A | G | C | C | A | A | T | T | G | A | T | G | A | A | C | T | G | T | T | A | A | T | T | - | - | - | C | A | C | A | A | A | G | C | A | C | A | C | A | C | C | A | A | - | - | - | - | - | - | - | - | - | T | C | T | C | A | A | G | C | T | C | A | A | A | A | A | T | G | C | C | A | T | A | G | C | C | C | A | A | A | C | A | T | T | A | C | A | C | G | - | - | - | - | - | - | - | - | - | A | T | T | T | T | - | - | - | - | A | T | A | A | T | T | T | T | G | T | A | C | T | - | - | - | - | T | A | A | T | T | A | T | A | A | T | A | A | T | G | T | G | T | G | T | T | G | A | C | A | T | T | G | C | A | A | A | A | T | A | A | A | T | C | A | C | T | T | C | C | G | T | - | - | - | - | - | - | - | - | - | - |
|  | MYZPE13164\_G006\_V2.0\_0001 | A | T | C | A | C | G | C | C | A | C | - | - | - | - | - | - | - | - | - | - | - | - | - | - | - | C | T | T | A | T | C | - | - | - | - | - | G | T | C | C | C | A | C | T | T | T | C | G | G | C | - | - | - | - | - | - | - | - | - | - | G | G | C | A | A | T | C | G | A | G | A | G | T | G | T | A | T | A | A | A | A | A | G | G | C | G | T | C | T | C | A | G | T | G | C | T | T | G | A | G | T | C | C | A | G | C | A | T | A | A | C | A | C | A | - | - | T | T | T | T | T | G | T | G | C | A | A | T | T | T | T | A | T | T | C | A | T | T | G | T | T | T | T | - | - | - | - | - | - | C | A | C | - | T | T | T | G | T | T | C | - | - | A | A | C | A | A | A | A | A | C | C | A | - | - | - | A | C | C | A | A | - | A | A | A | T | C | A | A | A | A | A | T | G | G | G | C | G | C | T | G | A | A | A | A | A | G | T | A | A | C | C | A | T | G | A | A | C | A | T | C | G | T | T | G | T | C | G | T | T | G | G | A | C | A | A | G | T | C | C | A | A | T | C | T | G | C | C | A | A | G | G | C | C | A | T | C | A | A | A | T | C | G | G | T | C | G | C | C | A | A | A | G | C | C | A | C | C | C | C | A | G | C | C | A | C | C | A | G | C | C | A | A | T | T | G | A | T | G | A | A | C | T | G | T | T | A | A | T | T | - | - | - | C | A | C | C | A | T | G | T | C | - | - | C | A | T | - | T | A | - | - | - | - | - | - | - | - | - | T | C | T | C | A | A | A | C | T | C | A | A | A | C | A | T | G | C | C | A | T | A | G | C | C | C | A | A | A | C | A | G | T | A | C | A | T | T | - | - | - | - | - | - | - | - | - | - | - | - | - | - | - | - | - | - | - | - | - | - | - | - | - | - | - | - | - | - | - | - | - | - | - | - | - | - | - | - | - | G | T | A | A | T | A | T | G | T | G | T | A | T | T | G | A | C | A | T | T | G | C | A | A | A | A | T | A | A | A | T | T | G | A | A | T | A | T | C | C | - | - | - | - | - | - | - | - | - | - |
|  | MYZPE13164\_G006\_V2.0\_0001 | A | T | C | A | C | G | C | C | A | C | G | T | T | C | G | - | - | - | - | - | - | - | - | - | - | - | - | - | - | - | - | - | - | - | - | - | C | T | C | C | C | A | C | T | T | T | T | G | T | C | G | A | T | T | T | T | C | G | C | T | C | A | C | T | T | A | T | C | A | G | A | G | C | G | T | A | T | A | T | - | A | A | G | A | C | G | C | C | G | G | A | G | A | T | C | T | C | G | A | T | T | C | C | G | G | C | A | T | A | A | C | A | C | A | T | C | T | T | T | T | T | G | T | G | C | A | A | A | T | C | T | A | T | T | C | T | T | T | G | T | T | T | T | - | - | - | - | - | C | C | A | C | - | A | T | T | T | C | T | C | - | - | C | T | C | A | A | A | A | A | C | C | A | - | - | - | A | C | C | A | A | - | A | A | A | C | C | A | A | A | A | A | T | G | G | G | C | G | C | T | G | A | A | A | A | G | G | T | A | T | C | C | A | T | G | A | A | C | A | T | C | G | T | T | G | T | C | G | T | C | G | G | A | C | A | A | G | T | C | C | A | A | T | C | T | G | C | C | A | A | G | G | C | C | A | T | C | A | A | G | A | C | C | G | T | C | G | C | C | A | A | G | G | C | C | A | C | C | C | C | A | G | C | C | A | C | C | A | G | C | C | A | A | T | T | G | A | T | G | A | A | C | T | G | T | T | A | A | T | T | - | - | - | C | A | T | G | T | A | G | T | C | - | - | C | A | T | C | G | A | - | - | - | - | - | - | - | - | - | T | C | T | C | A | A | A | C | T | C | A | A | A | A | A | T | G | C | C | A | T | A | G | C | C | C | A | A | A | C | A | A | T | A | C | G | C | A | A | A | T | T | A | T | A | A | T | A | A | T | A | T | - | - | - | - | G | T | A | G | T | A | A | A | T | A | A | A | T | - | - | G | T | A | T | A | T | T | - | A | T | A | A | T | T | A | A | C | G | T | A | T | T | G | C | C | A | C | T | G | C | A | A | A | A | T | A | A | A | C | C | A | C | G | A | T | T | T | T | - | - | - | - | - | - | - | - | - | - |
|  | MYZPE13164\_G006\_V2.0\_0001 | A | T | C | A | C | G | C | C | A | T | G | C | C | - | - | - | - | - | - | - | - | - | - | - | - | - | - | - | - | - | - | - | - | - | - | - | G | C | C | C | G | A | C | T | C | T | A | A | T | C | A | G | C | G | - | - | - | - | - | - | - | - | - | - | - | - | - | - | - | - | - | - | - | G | T | A | T | A | A | A | T | A | G | T | A | G | T | C | G | G | A | T | T | C | C | T | C | G | A | T | T | C | C | G | G | C | A | T | A | A | C | A | C | A | T | - | T | T | T | T | T | G | T | G | C | A | A | G | T | C | T | A | T | T | C | T | T | T | G | T | C | T | T | - | T | A | C | T | C | C | A | T | - | T | T | C | A | C | T | C | - | - | C | A | C | G | A | A | A | A | C | C | A | - | - | - | A | C | C | A | - | - | A | A | A | C | C | A | A | A | A | A | T | G | G | G | C | G | C | A | G | A | A | A | A | G | G | T | A | T | C | C | A | T | G | A | A | C | A | T | C | G | T | T | G | T | C | G | T | C | G | G | A | C | A | A | G | T | C | C | A | A | T | C | T | G | C | C | A | A | G | G | C | C | A | T | C | A | A | A | T | C | G | G | T | C | G | C | C | A | A | A | G | C | C | G | C | C | C | C | A | G | C | C | A | C | C | A | G | C | G | A | A | T | T | G | A | T | G | A | G | C | T | G | T | T | A | A | T | T | C | A | A | C | A | T | G | T | A | A | C | T | - | - | C | A | T | C | C | A | - | - | - | - | - | - | - | - | - | T | T | G | C | A | A | A | C | T | C | A | A | A | A | A | T | G | C | C | A | T | A | G | C | C | C | A | A | A | C | A | A | T | A | C | A | T | G | - | - | - | - | - | - | - | - | - | T | A | T | C | T | - | - | - | - | - | - | - | - | - | - | - | - | - | - | - | - | - | - | - | - | - | - | - | - | - | A | A | T | C | A | A | T | A | T | C | A | T | G | A | T | T | G | A | C | A | T | T | A | C | A | A | A | A | T | A | A | A | A | T | A | T | C | A | A | C | C | T | - | - | - | - | - | - | - | - | - | - |
|  | MYZPE13164\_G006\_V2.0\_0001 | A | T | C | A | C | G | C | C | A | C | A | A | C | - | - | - | - | - | - | - | - | - | - | - | - | - | - | - | - | - | - | - | - | - | - | - | T | C | C | C | C | A | C | T | T | C | T | G | T | C | G | G | C | C | - | - | - | - | - | - | G | C | C | T | C | G | A | A | T | T | T | A | T | G | T | A | T | A | - | A | A | A | A | G | C | G | T | C | C | G | G | C | T | C | T | T | C | G | A | T | T | C | T | G | G | C | A | T | A | A | C | A | C | A | T | - | T | T | T | T | T | G | T | G | C | A | A | A | T | C | T | A | T | T | C | T | T | T | G | T | C | T | T | - | T | A | C | T | C | T | A | T | - | T | T | C | A | C | T | C | - | - | C | A | C | G | A | A | A | A | C | C | A | - | - | - | A | C | C | A | - | - | A | A | A | C | C | A | A | A | A | A | T | G | G | G | C | G | C | A | G | A | A | A | A | G | G | T | A | T | C | C | A | T | G | A | A | C | A | T | C | G | T | T | G | T | C | G | T | C | G | G | A | C | A | A | G | T | C | C | A | A | T | C | T | G | C | C | A | A | G | G | C | C | A | T | C | A | A | A | T | C | G | G | T | C | G | C | C | A | A | A | G | C | C | G | C | C | C | C | A | G | C | C | A | C | C | A | G | C | C | A | A | T | T | G | A | T | G | A | A | C | T | G | T | T | A | A | T | T | C | A | A | C | A | T | G | T | A | A | T | T | - | - | C | A | T | A | A | G | - | - | - | - | - | - | - | - | - | C | A | A | C | A | A | A | C | T | C | A | A | A | A | A | T | G | C | C | A | T | A | G | C | C | C | A | A | A | C | A | T | T | A | C | A | C | T | - | - | - | - | - | - | - | - | - | A | A | C | A | T | - | - | - | - | G | T | A | A | T | T | A | T | A | C | T | A | A | A | - | A | T | G | T | A | T | G | T | G | A | A | T | T | A | A | T | A | T | T | A | T | G | T | A | C | A | T | T | G | C | A | A | A | A | T | A | A | A | T | C | A | G | T | C | T | T | C | T | - | - | - | - | - | - | - | - | - | - |
|  | MYZPE13164\_G006\_V2.0\_0001 | A | T | C | A | C | G | C | C | A | C | G | T | C | C | G | T | C | C | - | - | - | - | - | - | - | - | - | - | - | - | - | - | - | - | - | - | G | T | C | C | C | A | T | T | T | C | T | A | T | C | A | A | T | C | C | - | - | - | - | - | T | C | G | A | G | A | G | A | T | T | A | G | C | G | T | A | T | A | A | A | A | G | G | C | C | G | C | T | C | G | G | G | A | T | C | T | C | G | A | T | T | T | C | G | G | C | A | T | A | A | C | G | C | A | T | - | T | T | T | T | T | G | T | G | C | A | A | T | T | C | T | A | T | T | C | T | T | T | T | T | C | T | T | C | C | A | C | T | T | C | G | T | - | T | T | C | A | C | T | C | - | - | C | A | C | G | A | A | A | A | C | C | A | - | - | - | A | C | C | A | - | - | A | A | A | T | C | C | A | A | A | A | T | G | G | G | C | G | C | T | G | A | A | A | A | G | G | T | A | T | C | C | A | T | G | A | A | C | A | T | C | G | T | T | G | T | C | G | T | C | G | G | A | C | A | A | G | T | C | C | A | A | T | C | T | G | C | C | A | A | G | G | C | C | A | T | C | A | A | A | T | C | G | G | T | C | G | C | C | A | A | A | G | C | C | G | C | C | C | C | A | G | C | C | A | C | C | A | G | C | C | A | A | T | C | G | A | T | G | A | T | C | T | G | T | T | A | A | A | T | - | - | - | A | A | T | G | T | A | G | T | T | - | - | C | A | T | C | C | A | - | - | - | - | - | - | - | - | - | T | C | A | C | A | A | A | C | T | C | A | A | A | A | A | T | G | C | C | A | T | A | A | T | C | C | A | A | A | C | A | A | T | A | T | A | T | G | - | - | - | - | - | - | - | - | - | T | A | A | T | T | - | - | - | - | G | A | T | A | T | G | T | - | - | - | - | - | - | - | - | - | - | - | - | - | - | - | - | - | - | - | - | T | A | T | A | T | G | T | A | T | T | G | A | C | A | T | C | G | C | T | A | A | A | T | A | A | A | A | C | A | G | A | T | C | C | T | T | - | - | - | - | - | - | - | - | - | - |
|  | MYZPE13164\_G006\_V2.0\_0001 | A | T | C | A | C | G | C | C | A | T | T | T | A | C | G | C | C | C | C | - | - | - | - | - | C | C | C | C | C | T | T | - | - | - | - | - | A | C | A | C | T | A | C | C | T | T | A | G | A | C | G | - | - | - | - | - | - | - | - | - | G | T | C | T | T | T | T | A | T | G | T | G | T | G | T | A | T | A | A | A | A | G | A | A | C | G | G | C | C | G | T | G | T | G | G | T | C | A | A | A | T | C | C | G | A | C | A | T | A | A | C | A | C | A | T | - | T | T | T | T | T | G | T | G | C | A | G | C | T | T | T | A | T | T | C | T | T | T | G | T | T | T | T | - | - | - | - | - | C | C | A | C | - | A | T | A | A | C | T | A | - | - | C | T | C | A | A | A | A | A | C | C | A | - | - | - | A | C | C | A | A | - | A | A | A | C | C | A | A | A | A | A | T | G | G | G | C | G | C | T | G | A | A | A | A | G | G | T | A | T | C | C | A | T | G | A | A | C | A | T | C | G | T | T | G | T | C | G | T | C | G | G | A | C | A | A | G | T | C | C | A | A | T | C | T | G | C | C | A | A | G | G | C | C | A | T | C | A | A | A | T | C | G | G | T | C | G | C | C | A | A | A | G | C | C | A | C | C | C | C | A | G | C | C | A | C | C | A | G | C | C | A | A | T | T | G | A | T | G | A | A | C | T | G | T | T | A | A | T | T | T | - | - | T | G | T | C | A | A | G | T | C | T | C | C | A | T | A | A | A | - | - | - | - | - | - | - | - | - | C | C | T | C | A | A | A | A | T | C | A | A | A | A | A | T | G | C | C | A | T | A | G | T | C | C | A | A | A | A | C | T | T | C | A | A | A | C | - | - | - | - | - | - | - | - | - | A | G | T | A | C | - | - | - | - | A | T | A | G | T | A | A | A | A | A | T | G | T | G | T | A | A | T | C | A | T | A | T | A | T | T | T | A | T | T | A | C | G | T | A | T | T | G | A | C | A | T | C | G | C | A | A | A | A | T | A | A | A | T | C | T | G | T | C | T | T | T | C | - | - | - | - | - | - | - | - | - | - |
|  | MYZPE13164\_G006\_V2.0\_0000 | A | T | C | A | C | A | C | C | A | T | G | A | C | A | T | G | G | C | A | - | - | - | - | - | T | A | T | C | A | C | T | - | - | - | - | - | C | G | A | T | T | A | A | T | C | T | T | A | T | A | C | - | - | - | - | - | - | - | - | - | - | A | C | T | T | T | T | A | A | A | A | A | C | G | T | A | T | A | T | A | A | A | C | A | A | A | A | G | C | T | T | A | - | C | G | G | T | C | G | A | A | T | C | G | T | C | A | T | A | A | C | A | - | - | - | - | A | T | T | T | C | G | T | A | C | A | A | G | T | C | A | A | T | T | G | T | T | T | G | A | T | T | T | - | - | - | - | - | - | C | A | A | A | T | T | C | A | A | T | C | A | A | C | A | C | A | A | A | A | A | C | C | A | - | - | - | A | C | C | A | - | - | A | A | A | T | C | A | A | A | A | A | T | G | G | G | C | G | C | A | G | A | A | A | A | A | G | T | A | T | C | C | A | T | G | A | A | C | A | T | C | G | T | C | G | T | C | A | T | C | G | G | G | C | A | A | G | T | C | C | A | G | T | C | A | G | C | C | A | A | G | G | C | C | A | T | C | A | A | G | A | C | T | G | T | C | G | C | C | A | A | G | G | C | C | A | C | A | C | C | A | G | C | C | A | C | C | A | G | C | C | A | A | C | T | G | A | T | G | A | A | C | T | G | T | T | A | A | A | T | - | - | - | C | A | T | G | T | A | A | T | C | - | - | C | A | T | C | T | A | - | - | - | - | - | - | - | - | - | T | C | T | C | A | A | A | C | A | C | A | A | A | A | A | T | G | C | C | A | T | A | A | C | C | T | A | A | A | C | A | A | A | A | T | A | T | G | - | - | - | - | - | - | - | - | - | T | A | T | T | T | - | - | - | - | T | T | A | A | T | G | T | - | - | - | - | - | - | - | - | - | - | - | - | - | - | - | - | - | - | - | - | - | - | T | A | T | G | T | T | T | T | G | A | C | A | A | T | A | C | C | A | A | A | T | A | A | A | T | A | A | G | T | A | T | A | T | T | - | - | - | - | - | - | - | - | - | - |
|  | MYZPE13164\_G006\_V2.0\_0001 | A | T | C | A | C | A | C | C | A | T | A | C | C | A | T | G | C | C | A | - | - | - | - | - | T | A | T | C | A | C | A | - | - | - | - | - | C | G | A | T | T | A | A | T | C | T | T | A | T | A | C | - | - | - | - | - | - | - | - | - | G | C | T | T | T | T | A | A | A | A | A | A | C | G | T | A | T | A | T | A | A | A | C | A | A | A | C | G | C | T | T | A | C | C | T | G | C | C | C | A | A | T | C | G | T | C | A | T | A | A | C | A | C | A | - | - | A | T | T | T | C | G | T | A | C | A | A | G | T | C | A | A | T | T | G | T | T | T | G | A | T | T | T | - | - | - | - | - | - | C | A | A | G | T | T | C | A | A | T | C | C | A | C | A | C | A | A | A | A | A | C | C | A | - | - | - | A | C | C | A | - | - | A | A | A | T | C | A | A | A | A | A | T | G | G | G | C | G | C | A | G | A | A | A | A | A | G | T | A | T | C | C | A | T | G | A | A | C | A | T | C | G | T | C | G | T | C | G | T | C | G | G | A | C | A | A | G | T | C | C | A | G | T | C | T | G | C | C | A | A | G | G | C | C | A | T | C | A | A | G | A | C | C | G | T | C | A | C | C | A | A | G | G | C | C | A | C | A | C | C | A | G | C | C | A | C | C | A | G | C | C | A | A | C | T | C | A | T | G | A | A | C | T | G | T | T | A | A | A | T | - | - | - | C | A | T | G | T | A | A | A | C | - | - | C | G | T | C | A | A | - | - | - | - | - | - | - | - | - | T | C | T | C | A | A | A | C | A | C | A | A | A | A | A | T | G | C | C | A | T | A | A | C | C | T | A | A | A | C | A | A | A | A | T | A | T | G | - | - | - | - | - | - | - | - | - | T | A | T | T | T | - | - | - | - | C | T | A | A | C | A | T | - | - | - | - | - | - | - | - | - | - | - | - | - | - | - | - | - | - | - | - | - | - | T | A | T | G | C | T | T | T | G | A | C | A | C | A | A | C | C | A | A | A | T | A | A | A | T | A | A | G | T | A | T | A | T | T | - | - | - | - | - | - | - | - | - | - |
|  | MYZPE13164\_G006\_V2.0\_0001 | A | T | C | A | C | A | C | C | A | T | A | C | C | A | T | G | C | C | A | - | - | - | - | - | T | A | T | C | A | C | A | - | - | - | - | - | C | G | A | T | T | A | A | T | C | T | T | A | T | A | C | - | - | - | - | - | - | - | - | - | G | C | T | T | T | T | A | A | A | A | A | A | C | G | T | A | T | A | T | A | A | A | C | A | A | A | C | G | C | T | T | A | C | C | T | G | C | C | C | A | A | T | C | G | T | C | A | T | A | A | C | A | C | A | - | - | A | T | T | T | C | G | T | A | C | A | A | G | T | C | A | A | T | T | G | T | T | T | G | A | T | T | T | - | - | - | - | - | - | C | A | A | T | T | T | C | T | A | T | C | C | C | C | A | C | A | A | A | A | A | C | C | A | - | - | - | A | C | C | A | - | - | A | A | A | T | C | A | A | A | A | A | T | G | G | G | C | G | C | A | G | A | A | A | A | A | G | T | A | T | C | C | A | T | G | A | G | C | A | T | C | G | T | C | G | T | C | G | T | C | G | G | A | C | A | A | G | T | C | C | A | G | T | C | T | G | C | C | A | A | G | G | C | C | A | C | C | A | A | G | A | C | C | G | T | C | A | C | C | A | A | G | G | C | C | A | C | A | C | C | A | G | C | C | A | C | C | A | G | C | C | A | A | C | T | C | A | T | G | A | A | C | T | G | T | T | A | A | A | T | - | - | - | C | A | T | G | T | A | G | T | C | - | - | C | G | T | C | C | A | - | - | - | - | - | - | - | - | - | T | C | T | C | A | A | A | C | A | C | A | A | A | A | A | T | G | C | C | A | T | A | A | C | C | T | A | A | A | C | A | A | A | A | T | A | T | G | - | - | - | - | - | - | - | - | - | T | A | T | T | T | - | - | - | - | A | T | A | G | T | A | T | - | - | - | - | - | - | - | - | - | - | - | - | - | - | - | - | - | - | - | - | - | - | T | A | T | A | T | T | T | T | G | A | C | A | T | A | A | T | C | A | A | A | T | A | A | A | T | C | A | G | T | A | T | A | T | T | - | - | - | - | - | - | - | - | - | - |
|  |  | 1 |  |  |  |  |  |  |  |  | 1 | 0 |  |  |  |  |  |  |  |  | 2 | 0 |  |  |  |  |  |  |  |  | 3 | 0 |  |  |  |  |  |  |  |  | 4 | 0 |  |  |  |  |  |  |  |  | 5 | 0 |  |  |  |  |  |  |  |  | 6 | 0 |  |  |  |  |  |  |  |  | 7 | 0 |  |  |  |  |  |  |  |  | 8 | 0 |  |  |  |  |  |  |  |  | 9 | 0 |  |  |  |  |  |  |  |  | 1 | 0 | 0 |  |  |  |  |  |  |  | 1 | 1 | 0 |  |  |  |  |  |  |  | 1 | 2 | 0 |  |  |  |  |  |  |  | 1 | 3 | 0 |  |  |  |  |  |  |  | 1 | 4 | 0 |  |  |  |  |  |  |  | 1 | 5 | 0 |  |  |  |  |  |  |  | 1 | 6 | 0 |  |  |  |  |  |  |  | 1 | 7 | 0 |  |  |  |  |  |  |  | 1 | 8 | 0 |  |  |  |  |  |  |  | 1 | 9 | 0 |  |  |  |  |  |  |  | 2 | 0 | 0 |  |  |  |  |  |  |  | 2 | 1 | 0 |  |  |  |  |  |  |  | 2 | 2 | 0 |  |  |  |  |  |  |  | 2 | 3 | 0 |  |  |  |  |  |  |  | 2 | 4 | 0 |  |  |  |  |  |  |  | 2 | 5 | 0 |  |  |  |  |  |  |  | 2 | 6 | 0 |  |  |  |  |  |  |  | 2 | 7 | 0 |  |  |  |  |  |  |  | 2 | 8 | 0 |  |  |  |  |  |  |  | 2 | 9 | 0 |  |  |  |  |  |  |  | 3 | 0 | 0 |  |  |  |  |  |  |  | 3 | 1 | 0 |  |  |  |  |  |  |  | 3 | 2 | 0 |  |  |  |  |  |  |  | 3 | 3 | 0 |  |  |  |  |  |  |  | 3 | 4 | 0 |  |  |  |  |  |  |  | 3 | 5 | 0 |  |  |  |  |  |  |  | 3 | 6 | 0 |  |  |  |  |  |  |  | 3 | 7 | 0 |  |  |  |  |  |  |  | 3 | 8 | 0 |  |  |  |  |  |  |  | 3 | 9 | 0 |  |  |  |  |  |  |  | 4 | 0 | 0 |  |  |  |  |  |  |  | 4 | 1 | 0 |  |  |  |  |  |  |  | 4 | 2 | 0 |  |  |  |  |  |  |  | 4 | 3 | 0 |  |  |  |  |  |  |  | 4 | 4 | 0 |  |  |  |  |  |  |  | 4 | 5 | 0 |  |  |  |  |  |  |  | 4 | 6 | 0 |  |  |  |  |  |  |
|  | GUIDANCE2 SCORE |  |  |  |  |  |  |  |  |  |  |  |  |  |  |  |  |  |  |  |  |  |  |  |  |  |  |  |  |  |  |  |  |  |  |  |  |  |  |  |  |  |  |  |  |  |  |  |  |  |  |  |  |  |  |  |  |  |  |  |  |  |  |  |  |  |  |  |  |  |  |  |  |  |  |  |  |  |  |  |  |  |  |  |  |  |  |  |  |  |  |  |  |  |  |  |  |  |  |  |  |  |  |  |  |  |  |  |  |  |  |  |  |  |  |  |  |  |  |  |  |  |  |  |  |  |  |  |  |  |  |  |  |  |  |  |  |  |  |  |  |  |  |  |  |  |  |  |  |  |  |  |  |  |  |  |  |  |  |  |  |  |  |  |  |  |  |  |  |  |  |  |  |  |  |  |  |  |  |  |  |  |  |  |  |  |  |  |  |  |  |  |  |  |  |  |  |  |  |  |  |  |  |  |  |  |  |  |  |  |  |  |  |  |  |  |  |  |  |  |  |  |  |  |  |  |  |  |  |  |  |  |  |  |  |  |  |  |  |  |  |  |  |  |  |  |  |  |  |  |  |  |  |  |  |  |  |  |  |  |  |  |  |  |  |  |  |  |  |  |  |  |  |  |  |  |  |  |  |  |  |  |  |  |  |  |  |  |  |  |  |  |  |  |  |  |  |  |  |  |  |  |  |  |  |  |  |  |  |  |  |  |  |  |  |  |  |  |  |  |  |  |  |  |  |  |  |  |  |  |  |  |  |  |  |  |  |  |  |  |  |  |  |  |  |  |  |  |  |  |  |  |  |  |  |  |  |  |  |  |  |  |  |  |  |  |  |  |  |  |  |  |  |  |  |  |  |  |  |  |  |  |  |  |  |  |  |  |  |  |  |  |  |  |  |  |  |  |  |  |  |  |  |  |  |  |  |  |  |  |  |  |  |  |  |  |  |  |  |  |  |  |  |  |  |  |  |  |  |  |  |  |  |  |  |  |  |  |  |  |  |  |  |  |  |  |  |  |  |  |  |  |  |  |  |  |  |  |  |  |  |  |  |  |  |  |  |  |  |

 **Column** 
  
**Legend:  
  
The alignment confidence scale:**  

9  8  7  6  5  4  3  2  1 

|  |  |  |
| --- | --- | --- |
| **Confident** | **<--->** | **Uncertain** |

|  |  |
| --- | --- |
|  | Insufficient Data |
