## Supplementary material for "An aphid host-responsive RNA transcript that migrates systemically in plants promotes aphid reproduction": Suppl. Fig. S9

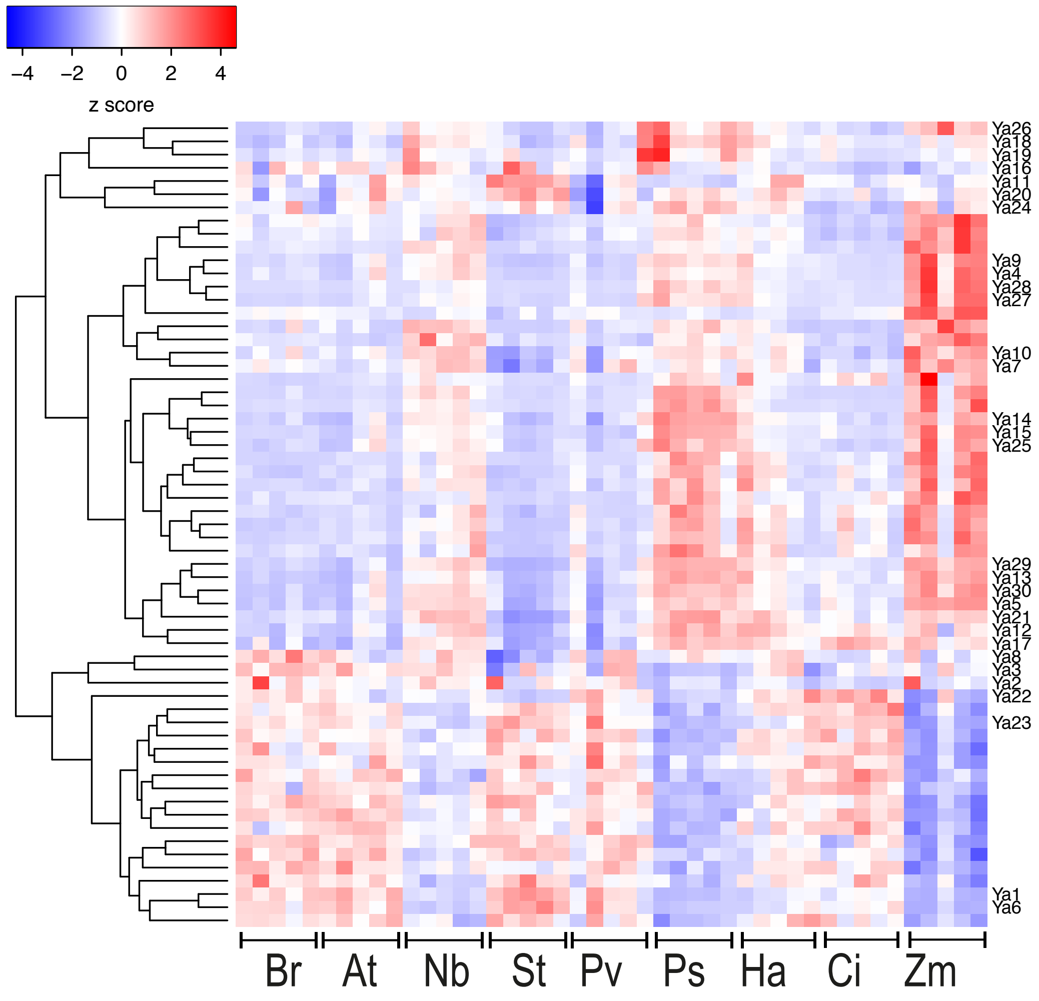


**Suppl. Fig. S9.** Heatmap of darkslateblue module after manual annotation of the *Ya* family. WGCNA analysis was repeated with the corrected set of *Ya* genes. Heatmap was generated using log-transformed TPM values of all genes in the module. Br, *B. rapa*; At, *A. thaliana*; Nb, *N. benthamiana*; St, *S. tuberosum*; Ci, *C. indicum*; Ha, *H. annuus*; Ps, *P. sativum*; Pv, *P. vulgaris*; Zm, *Z. mays*. Rows are scaled and represented as z-score.
