## Supplementary material for "An aphid host-responsive RNA transcript that migrates systemically in plants promotes aphid reproduction": Suppl. Fig. S10

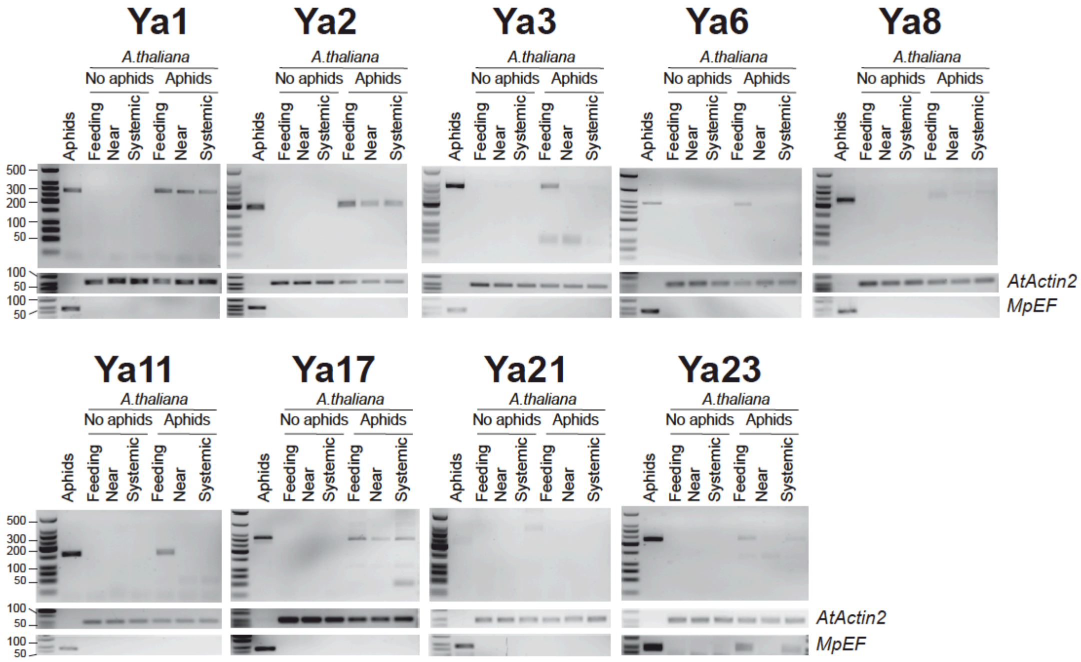


**Suppl. Fig. S10.** RT-PCR experiments showing translocation of 9 *Ya* transcripts into plants and systemic migration of *Ya1*, *Ya2*, and *Ya17* transcripts. Experimental setup is shown in Fig. 4A. The *Ya* transcripts were amplified by RT-PCR using gene-specific primers (**Suppl. Fig. S10; Suppl. Table S7**) and PCR products were separated by 3% agarose gel. The bands corresponding to the PCR products were extracted from the gel and sequenced directly by forward and/or reverse primers (**Suppl. Fig. S10**) to verify the identity of the Ya sequence.
