## Supplementary material for "An aphid host-responsive RNA transcript that migrates systemically in plants promotes aphid reproduction": Suppl. Fig. S11

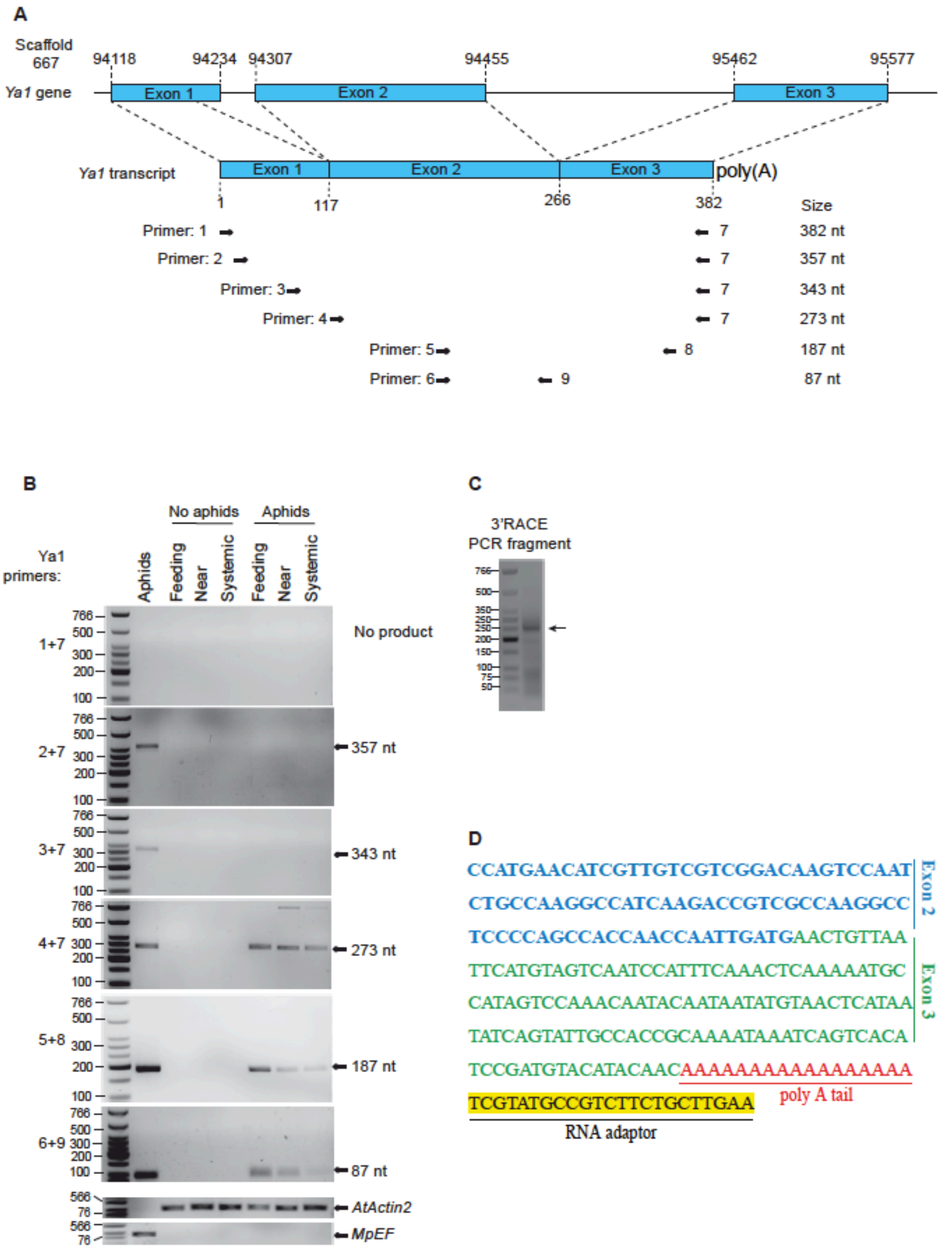


**Suppl. Fig. S11.** Identification of *Ya1* transcript sizes in aphids and plants. (**A**) Transcript model of the *Ya1* gene. The three exons of the *Ya1* gene that form the transcript are indicated as blue rectangles. The numbers above the Ya gene indicate the first and last nucleotide of each exon on scaffold 667 of the *M. persicae* genome assembly. The locations of the Ya1 primers 1-9 (**Suppl. Table S4**) used to assess the presence of *Ya1* transcripts in aphids and in plants by RT-PCR are indicated as black arrows. (**B**) A 357-nt *Ya1* transcript was detected in aphids and not in plants, whereas the largest *Ya1* transcript detected in plants at aphid feeding sites and systemic locations is 273-nt (primers 4 and 7). Experimental setup is shown in Fig. 4A. RT-PCR products generated with primers shown in A were separated on 3% agarose gels. Fragment sizes are indicated at right of the gels and correspond with the expected sizes as shown in A. Primers 1 and 7 did not generate a product in aphids nor in plants. (**C**) Results of a 3' RACE experiment. The 3' RACE PCR products were separated a 3% agarose gel. The band isolated from the gel and sequenced is shown with an arrow at right of the gel. (**D**) Sequence of the 3' RACE PCR fragment of the band shown in C. The sequence is identical to the sequences of the end of exon 2 (letters in blue font) and the entire exon 3 (letters in green font) of the annotated *Ya1* gene in the *M. persicae* genome. The *Ya1* transcript has a poly(A) tail (letters in red font). The RNA adaptor sequence used in 3' RACE protocol is indicated in yellow.
