## Supplementary material for "An aphid host-responsive RNA transcript that migrates systemically in plants promotes aphid reproduction": Suppl. Fig. S12

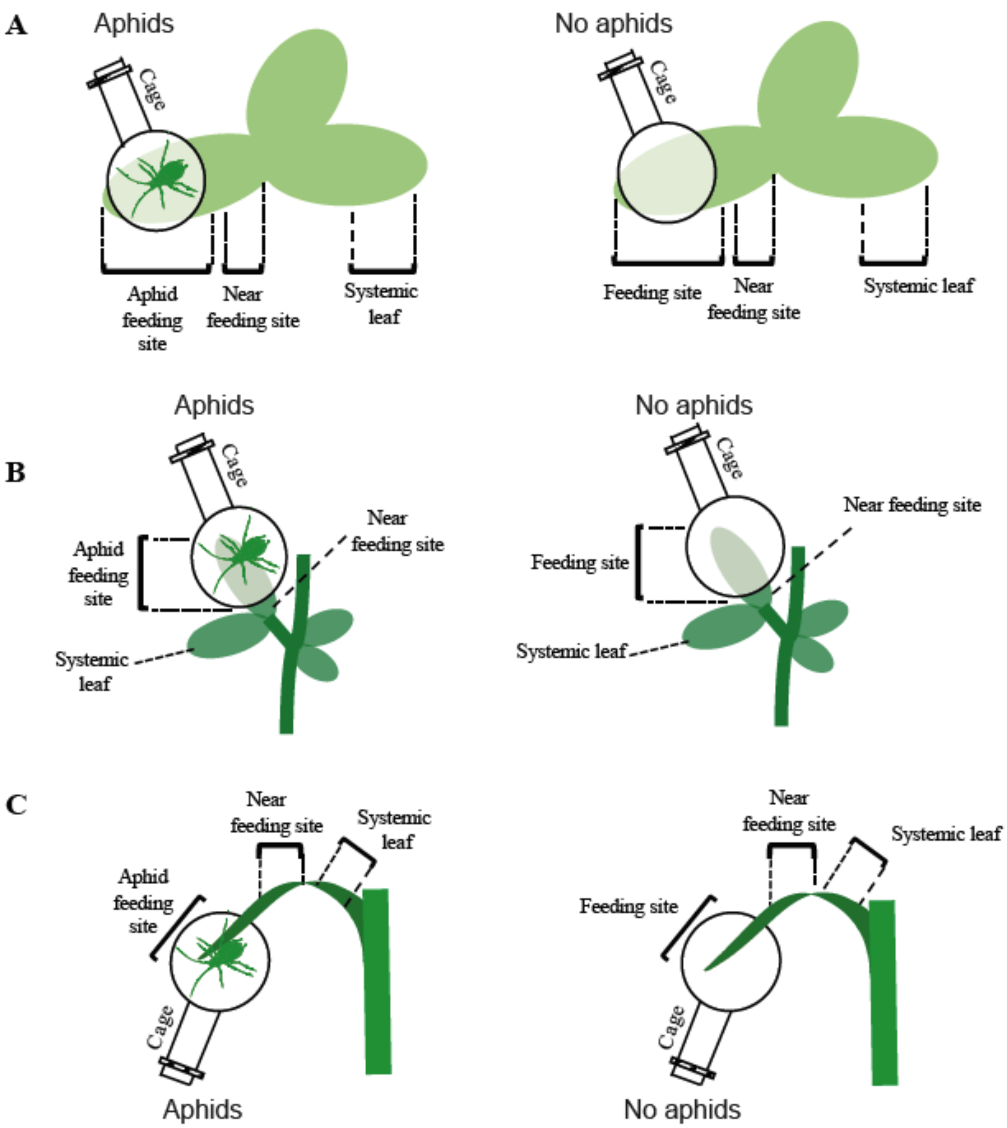


**Suppl. Fig. S12**. Schematic overview of experimental setups to investigate systemic migration of aphid *Ya1* transcripts in *Brassica rapa*, *Pisum sativum* and *Zea mays*. (A) *Brassica rapa.* (B) *Pisum sativum.* (C) *Zea mays*. Aphid feeding site is the caged parts of the leaf. Systemic sites include near-feeding sites that are located just externally of the caged leaf areas and more systemic locations that are different leaves for *B. rapa* and *P. sativum* and further away of the caged area on the same leaf of *Z. mays*.
