## Supplementary material for "An aphid host-responsive RNA transcript that migrates systemically in plants promotes aphid reproduction": Suppl. Fig. S13

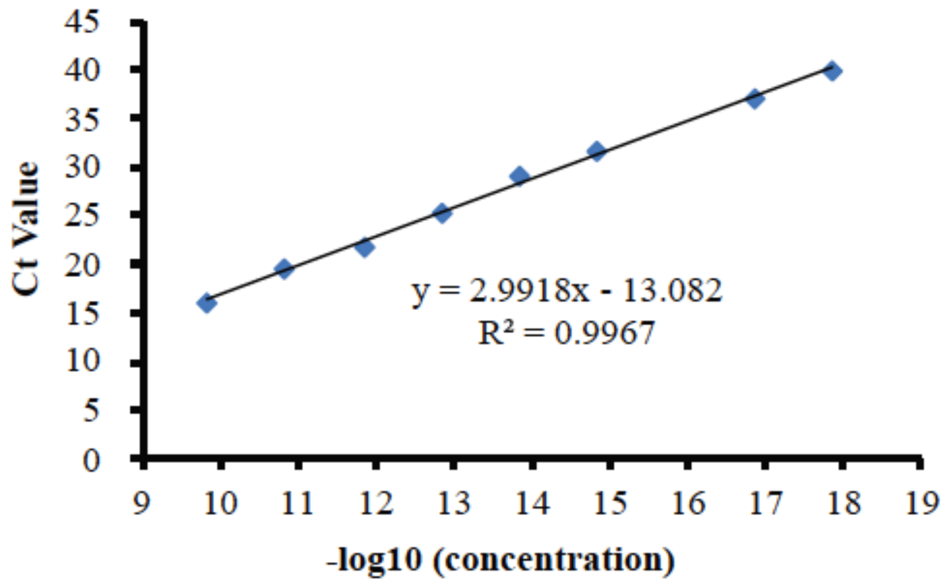


**Suppl. Fig. S13.** Standard concentration curve of *Ya1* and cycles of PCR amplification. The 273-nt Ya1 fragment in the plasmid pBI121 was used as the PCR template. qPCR were performed on a serial of dilution of the plasmid pBI121_35S::Ya1 from the highest concentration $1.44E^{-10}$ g/ μL to the lowest concentration $1.44E^{-18}$ g/ μL. x axis are minus log 10 transferred concentrations.
