## Supplementary material for "An aphid host-responsive RNA transcript that migrates systemically in plants promotes aphid reproduction": Suppl. Fig. S14

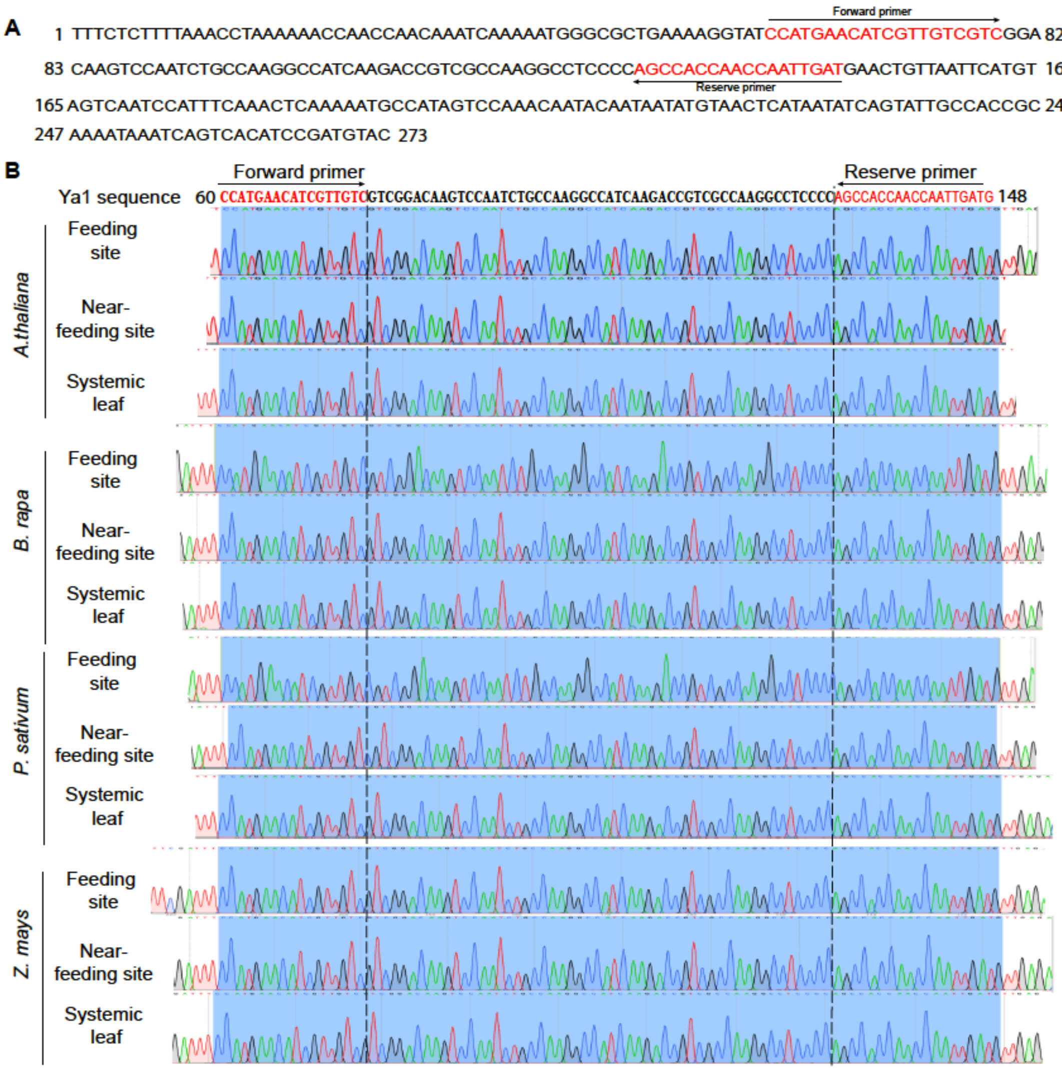


**Suppl. Fig. S14.** Sequences of RT-PCR products of *M. persicae Ya1* transcripts amplified from plants. The *Ya1* transcripts were amplified with Ya1 forward and reverse primers from feeding sites, near-feeding sites, and systemic sites of *A. thaliana*, *B. rapa*, *P. sativum* and *Z. mays* as per experimental setups shown in **Fig. 4A** and **Suppl. Fig. S12**. The sequences shown were identical to the corresponding region within the *Ya1* gene.
