## Supplementary material for "An aphid host-responsive RNA transcript that migrates systemically in plants promotes aphid reproduction": Suppl. Fig. S15

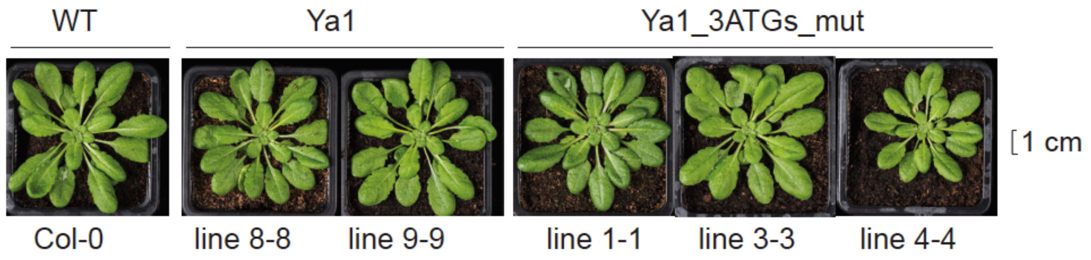


**Suppl. Fig. S15.** Comparison of phenotypes of *Ya1* transgenic *A. thaliana* plants. Transgenic *A. thaliana* (Col-0) plants that express *Ya1* and *Ya1_3ATGs_mut* under control of the *35S* promoter did not show obvious morphological differences compared to the non-transformed wild type Col-0 plants.
