## Supplementary material for "An aphid host-responsive RNA transcript that migrates systemically in plants promotes aphid reproduction": Suppl. Fig. S16

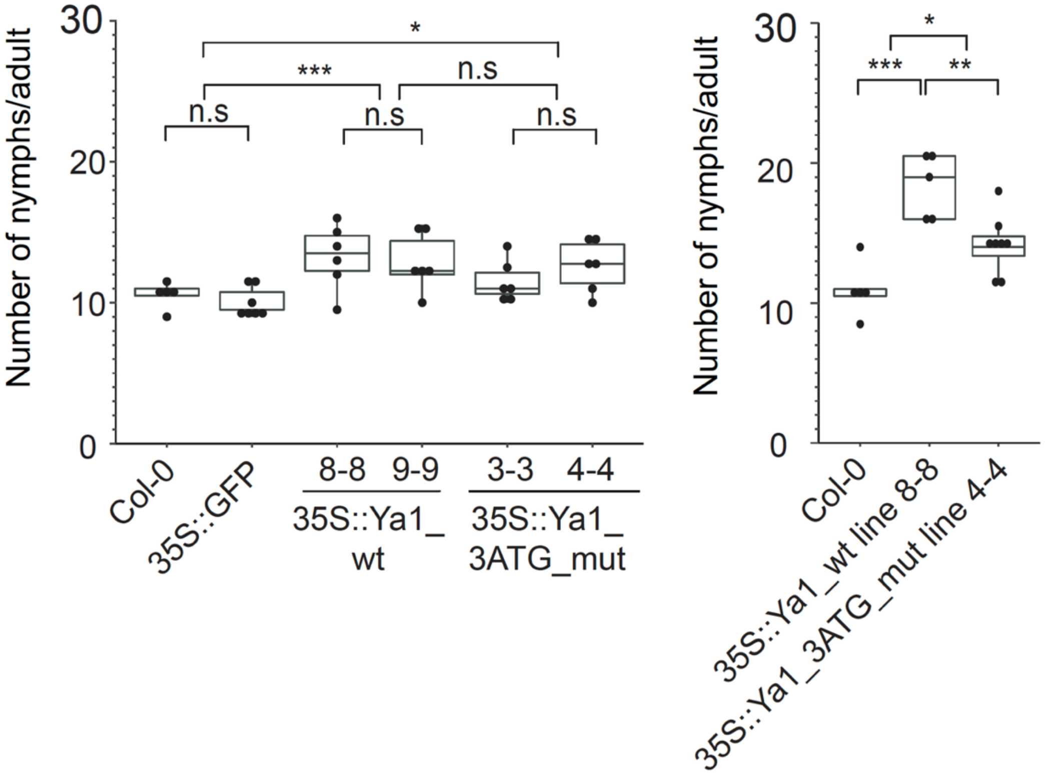


**Suppl. Fig. S16.** Repeats showing that *M. persicae Ya* lncRNA promotes *M. persicae* colonization on *A. thaliana*. Stable expression of *Ya1_wt* and *Ya1_3ATG* promotes *M. persicae* reproduction on plants. Each data point (black dot) represents number of nymphs produced by an adult female aphid per plant. Box plots show distribution of data points collected from n = 5-8 female aphids per *A. thaliana* line. $*p < 0.05$, $**p < 0.01$, $***p < 0.001$, ANOVA followed by a Tukey–Kramer post-hoc test.
