## Supplementary material for "An aphid host-responsive RNA transcript that migrates systemically in plants promotes aphid reproduction": Suppl. Table S1

**Table S1.** Transcriptome assembly statistics. RNA-seq reads were derived from samples LIB1777 (1) and colonies on 5 independent plants (1-5) of *Arabidopsis thaliana* (At); Br, *Brassica rapa* (Br), *Nicotiana benthamiana* (Nb), *Solanum tuberosum* (St), *Chrysanthemum indicum* (Ci), *Helianthus annuus* (Ha), *Pisum sativum* (Ps), *Phaseolus vulgaris* (Pv) and *Zea mays* (Zm). The reads were mapped to the *M. persicae* G006 genome assembly (1).

^a^ Total transcripts were assembled from mapped reads.

^b,c^ Transcripts that correspond to genes identified previously in Mathers et al. 2017 (1).

^d^ Novel transcript as identified in the transcriptome assembly reported herein.

| **Samples** | **Total reads** | **Mapped reads** | **Mapped reads (%)** | **Transcripts** | | | |
| --- | --- | --- | --- | --- | --- | --- | --- |
|  |  |  |  | **Total^a^** | **Known^b^** | **Genes^c^** | **Novel^d^** |
| LIB1777 | 182928264 | 117579937 | 64.28 | 87444 | 18750 | 11332 | 68694 |
| At1 | 35027735 | 31609619 | 90.24 | 61983 | 17781 | 10486 | 44202 |
| At2 | 30317626 | 27641464 | 91.17 |  |  |  |  |
| At3 | 32270976 | 29367540 | 91 |  |  |  |  |
| At4 | 33889504 | 30765104 | 90.78 |  |  |  |  |
| At5 | 29871796 | 27369312 | 91.62 |  |  |  |  |
| Ps1 | 33997244 | 31321585 | 92.13 | 60443 | 17737 | 10528 | 42706 |
| Ps2 | 33572880 | 30992391 | 92.31 |  |  |  |  |
| Ps3 | 30706147 | 28148247 | 91.67 |  |  |  |  |
| Ps4 | 38851016 | 35655100 | 91.77 |  |  |  |  |
| Ps5 | 23736763 | 20424890 | 86.05 |  |  |  |  |
| Br1 | 36105440 | 33390772 | 92.48 | 58293 | 17750 | 10489 | 40543 |
| Br2 | 27368493 | 23985300 | 87.64 |  |  |  |  |
| Br3 | 28484604 | 26110873 | 91.67 |  |  |  |  |
| Br4 | 27486534 | 25035464 | 91.08 |  |  |  |  |
| Br5 | 30044024 | 27584912 | 91.81 |  |  |  |  |
| Ci1 | 31213071 | 28395530 | 90.97 | 65917 | 17856 | 10621 | 48061 |
| Ci2 | 26059223 | 23430480 | 89.91 |  |  |  |  |
| Ci3 | 27469301 | 24621304 | 89.63 |  |  |  |  |
| Ci4 | 32534564 | 29376935 | 90.29 |  |  |  |  |
| Ci5 | 36510146 | 33342272 | 91.32 |  |  |  |  |
| Zm1 | 23898955 | 21330499 | 89.25 | 61157 | 17886 | 10610 | 43271 |
| Zm2 | 32663254 | 30362632 | 92.96 |  |  |  |  |
| Zm3 | 38833395 | 32998965 | 84.98 |  |  |  |  |
| Zm4 | 36223785 | 33199732 | 91.65 |  |  |  |  |
| Zm5 | 37985100 | 34900276 | 91.88 |  |  |  |  |
| Nb1 | 26749263 | 23973979 | 89.62 | 57533 | 17567 | 10444 | 39966 |
| Nb2 | 28476921 | 25844725 | 90.76 |  |  |  |  |
| Nb3 | 30481687 | 28348224 | 93 |  |  |  |  |
| Nb4 | 31075574 | 28936577 | 92.12 |  |  |  |  |
| Nb5 | 25055052 | 22819867 | 91.08 |  |  |  |  |
| Ps1 | 22202831 | 19882967 | 89.55 | 56431 | 17590 | 10488 | 38841 |
| Ps2 | 26907680 | 24899984 | 92.54 |  |  |  |  |
| Ps3 | 30923478 | 28678037 | 92.74 |  |  |  |  |
| Ps4 | 37835896 | 34845885 | 92.10 |  |  |  |  |
| Ps5 | 30745946 | 28130700 | 91.49 |  |  |  |  |
| St1 | 22647808 | 19265819 | 85.07 | 60662 | 17731 | 10537 | 42931 |
| St2 | 26239515 | 23358283 | 89.02 |  |  |  |  |
| St3 | 29668113 | 27521716 | 92.77 |  |  |  |  |
| St4 | 32534760 | 30241012 | 92.95 |  |  |  |  |
| St5 | 32905595 | 30331494 | 92.18 |  |  |  |  |
| Ha1 | 34475345 | 31538679 | 91.48 | 58152 | 17744 | 10490 | 40408 |
| Ha2 | 39135872 | 36179328 | 92.45 |  |  |  |  |
| Ha3 | 32762153 | 29883530 | 91.21 |  |  |  |  |
| Ha4 | 20135006 | 18438242 | 91.57 |  |  |  |  |
| Ha5 | 24340322 | 22404095 | 92.05 |  |  |  |  |
| **Total** | | | | 45972 | 30127 | 18529 | 15845 |
