## Supplementary material for "An aphid host-responsive RNA transcript that migrates systemically in plants promotes aphid reproduction": Suppl. Table S4

**Table S4.** Statistics of analyses of RNA-seq data retrieved from aphid-exposed (feeding sites) and non-exposed (control) leaves. RNA-seq reads derived from four biological replicates per treatment were mapped to both *A. thaliana* Col-0 (TAIR10) and *M. persicae* (Mp) genomes. *Reads mapping to Mp were realigned to At to find uniquely mapping Mp reads. The unique reads were then assigned to the transcripts obtained from the transcriptome assembly v2. ^ represents transcripts at TPM ≥ 50 in at least one biological replicate and presence in at least three replicates.

|  | Biological Replicate | Total reads  (million) | Reads mapped to At  (million) | Reads mapped to At (%) | Reads mapped to Mp | Reads mapped to Mp (%) | Reads mapped to At | Unique Mp reads* | Mp transcripts (TPM ≥ 50) | |
| --- | --- | --- | --- | --- | --- | --- | --- | --- | --- | --- |
|  |  |  |  |  |  |  |  |  | Number of transcripts | Number of low-coding potential transcripts |
| Aphid feeding  sites | 1 | 28 | 25 | 90.61 | 18007 | 0.0631 | 5 | 18002 | 2928 | 170 |
|  | 2 | 21 | 19 | 90.02 | 242628 | 1.1091 | 0 | 242628 | 2599 | 280 |
|  | 3 | 24 | 22 | 90.39 | 3847 | 0.0153 | 2 | 3845 | 1837 | 79 |
|  | 4 | 24 | 21 | 88.11 | 657024 | 2.6969 | 3 | 657021 | 3154 | 336 |
|  |  |  |  |  | | | | | **3186^** | **201^** |
| No aphid control  sites | 1 | 31 | 28 | 90.53 | 35 | 0.0001 | 23 | 12 | 0 | 0 |
|  | 2 | 22 | 20 | 91.08 | 9 | 0.0000 | 5 | 4 | 0 | 0 |
|  | 3 | 28 | 26 | 90.79 | 5 | 0.0000 | 5 | 0 | 0 | 0 |
|  | 4 | 21 | 19 | 90.55 | 17 | 0.0001 | 14 | 3 | 0 | 0 |
